# The Critical Role of the 2’-OH group in Phase Separation and Percolation Transitions of RNA

**DOI:** 10.1101/2025.02.26.638501

**Authors:** Gable M. Wadsworth, Dilimulati Aierken, George M. Thurston, Jerelle A. Joseph, Priya R. Banerjee

## Abstract

Mg^2+^ ion-dependent RNA liquid–liquid phase separation with lower critical solution temperatures is driven by the phosphate backbone and modulated by the solvation property of nucleobases. Here, we report a key role of the 2’-OH group of the ribose sugar in RNA condensation in the presence of divalent cations. We show that 2’-deoxyribose inhibits nucleic acid phase separation and suppresses the intra-condensate networking transition, known as percolation, that underlies condensate dynamical arrest. All-atom simulations reveal increased solvation and compaction of single-stranded DNA compared to RNA, suggesting an unintuitive role of chain flexibility in modulating heat-induced nucleic acid phase separation and percolation transitions. Further, 2’-O-methylation (2’-O-Me) of RNA, a common sugar modification, lowers the driving force of RNA phase transitions. These results highlight the diverse physicochemical parameters governing nucleic acid phase behavior and suggest how sugar modifications may have evolved to robustly tune the formation and dynamical arrest of RNA condensates.

---

In living cells, ribonucleic acids (RNA) and deoxyribonucleic acids (DNA) are both thought to participate in phase separation with proteins to form, for example, ribonucleoprotein (RNP) granules in the case of RNA ^1–4^, chromatin domains in the case of DNA ^5–10^, or nucleoli that contain both ^11–15^. Since many of these condensates contain both proteins and nucleic acids (NAs), it is conceivable that a molecular grammar has evolved that enables higher-order protein–NA interactions driving phase separation in a sequence- and structure-specific manner ^16–18^. Recent investigations of phase separation of both RNP granules and chromatin have in large part focused on the ability of proteins to undergo homo- and heterotypic condensation into liquid-like droplets. While many of these naturally occurring protein components undergo enthalpically driven phase separation with Upper Critical Solution Temperatures (UCSTs), desolvation entropy plays a central role in NA phase separation ^19^. This manifests in RNA ^19^ and DNA ^20,21^ phase transitions with Lower Critical Solution Temperatures (LCSTs) *in vitro* in the presence of divalent cations such as Mg^+2^. Mechanistically, desolvation of the phosphate backbone and a stronger coordination of Mg^+2^ ions upon heating drives LCST-type NA phase separation, whereas, the solvation properties and interaction specificity of nucleobases profoundly modulate the transition temperature and reversibility of the condensation process, leading to a rich variety of hysteretic NA phase behavior ^22^.

Reduced to the simplest description, the distinction between RNA and DNA is that of a single oxygen atom in the ribose sugar ^23^. These sixteen Daltons per monomer unit are sufficient to create profound differences between these two nucleic acids in terms of structure, thermodynamics, and function. Historically, it has been understood that in the presence of polyvalent ions DNA condenses ^24–27^ while RNA does not ^28–31^ except for special cases such as poly(rU) RNA ^29,32^. However, these studies were focused on the single-chain properties of nucleic acids in dilute solutions below the critical concentration (*c*_sat_) where mesoscopic condensates form. Moreover, by design, these studies precluded RNA condensates ^19,33^ as insoluble aggregates from their analysis ^29–31^. RNA phase separation in the presence of Mg^2+^ ions is ubiquitous and is often coupled to an intra-condensate percolation transition, leading to dynamically arrested irreversible condensates ^19,33,34^. Percolation is a connectivity transition defined by an exponential increase in the number of bonds formed above a critical parameter such as a critical concentration, *c*_prc_, or critical temperature, T_prc_. Percolation can arise either from physical or chemical bond formation ^35^. Phase separation drives percolation when *c*_prc_ exceeds the dense phase RNA concentration (*c*_dense_) ^22,36^. Here we leverage the concept of percolation in an operative sense through the phenomenological observation of percolation in two ways, first the persistence of condensates (irreversibility) below the lower cloud point temperature (LCPT) e.g. T_phase_ < T_percolation_, signifying thermo-reversible bond formation, and second as the remodeling of percolated condensates observed as shape relaxation upon heating above the percolation temperature, T_percolation_ ^19,22^. These observations of the thermos-reversible dynamical arrest implicate temperature-dependent intra-condensate RNA-RNA interactions leading to the percolation of the intra-condensate network, akin to rigidity percolation in disordered soft materials ^37,38^. Intra-condensate RNA percolation is RNA sequence-specific and may provide a thermodynamic framework for understanding the distinct physiological and pathological phase behaviors of RNA ^22^. RNA sequences associated with diseases, such as Huntington’s disease, Myotonic dystrophy, and ALS/FTD, have been observed both to form irreversible condensates and to recruit proteins in a gain-of-function manner within cells ^19,22,33,39,40^. These discoveries in RNA biophysics also have the potential to further DNA nano and soft matter technology ^41–43^. However, despite their many similarities in chemical properties and interactions (**Supplemental Figure S1**), it is not understood if single-stranded DNAs (ssDNAs) have similar intrinsic properties of tunable sequence-specific phase behavior.

In this study, we report experimental evidence and provide mechanistic insights gleaned from molecular simulations indicating that RNAs universally have higher tendencies to phase separate than ssDNA with the identical primary sequence. RNA phase separation is also tightly coupled to percolation in a sequence-specific manner, leading to the formation of large fractal-like networks and dynamically arrested condensates, which is not typically observed in ssDNA condensates. Using molecular dynamics (MD) simulations and small angle x-ray scattering (SAXS), we identify that ssDNA tends to be more compact than RNA while simultaneously retaining more water molecules in the solvation shell than RNA, providing insights into why RNA phase separates more readily upon heating. Furthermore, we demonstrate that both ssDNA and RNA can be switched from reversible phase separation to irreversible percolation via protonation of specific nucleobases. We also identify G-quadruplex forming sequences as strongly percolating systems for both ssDNA and RNA, suggesting that nucleic acid percolation can be engineered at the single-chain sequence level. The distinct phase behavior of ssDNA and RNA motivated us to probe the role of naturally occurring post-transcriptional sugar modifications (PTMs) in RNA on the phase behavior and percolation properties, leading us to discover that 2’-O-methylation (2’-O-Me) of RNA lowers the driving force of RNA phase transitions and makes RNA condensation reversible. This suggests sugar modification as a tunable switch to regulate RNA condensation.

## CAG repeat ssDNA is more compact than RNA

Classical theories of associative polymers predict that the density transition of the system is accompanied by a single-chain coil-to-globule transition for both UCST- and LCST-type transitions (**Figure 1A**) ^44^. For disordered protein systems, the strength of intra-chain interaction has been extensively used as a proxy for the likelihood to form physical crosslinks between protein chains at higher concentrations leading to the expectation that single-chain compaction scales with phase separation propensity ^45–48^. To uncover whether such a relationship exists for NAs, we employed all-atom molecular dynamics simulations and SAXS experiments. We observe that CAG-repeat ssDNA has systematically lower radii of gyration (*R*_g_) than CAG-repeat RNA when varying temperature (**Figure 1.B, C; Supplementary Figures S2-S4**). Additionally, we observe that R_g_ values from fitting SAXS data are not dependent on nucleic acid concentration confirming that these measurements are in the single-chain regime (**Supplementary Figure S3**; for more details see **Supplementary Note 1**). Importantly, both ssDNA and RNA become more compact upon increasing temperature, which signifies that phase separation is facilitated upon heating leading to an LCST-type transition (**Figure 1.B-C; Supplementary Figure S2.A**). R_g_ values from simulations are in reasonable agreement with SAXS results **(Supplementary Figure S2.B)**. Simulations predict that Mg^2+^ ions are closely associated with RNA in the first solvation shell, which is less frequently observed for ssDNA (**Figure 1.D, Supplementary Figure S5.A, E**) and is accompanied by a reduction of the number of water molecules in the first solvation shell (**Figure 1.E, Supplementary Figure S5.B, F**). This observation suggests that despite being less compact than ssDNA, RNA may have a greater tendency towards LCST-type phase separation than ssDNA in the presence of Mg^2+^ ions; that is, in RNA systems increased interactions with Mg^2+^ provide greater enthalpic stabilization upon phase separation, while reduced interactions with water lead to a smaller entropic penalty upon demixing. While in disordered protein systems, greater compaction suggests that homotypic interactions are stronger leading to a greater tendency to phase separate at higher concentrations, entropically driven desolvation leading to phase separation of structured nucleic acids does not seem to follow this phenomenological rule. This is likely due to a complex interplay between dominant enthalpic versus entropic contributions to phase separation and the stability of the folded structures ^49,50^.

**Figure 1.**
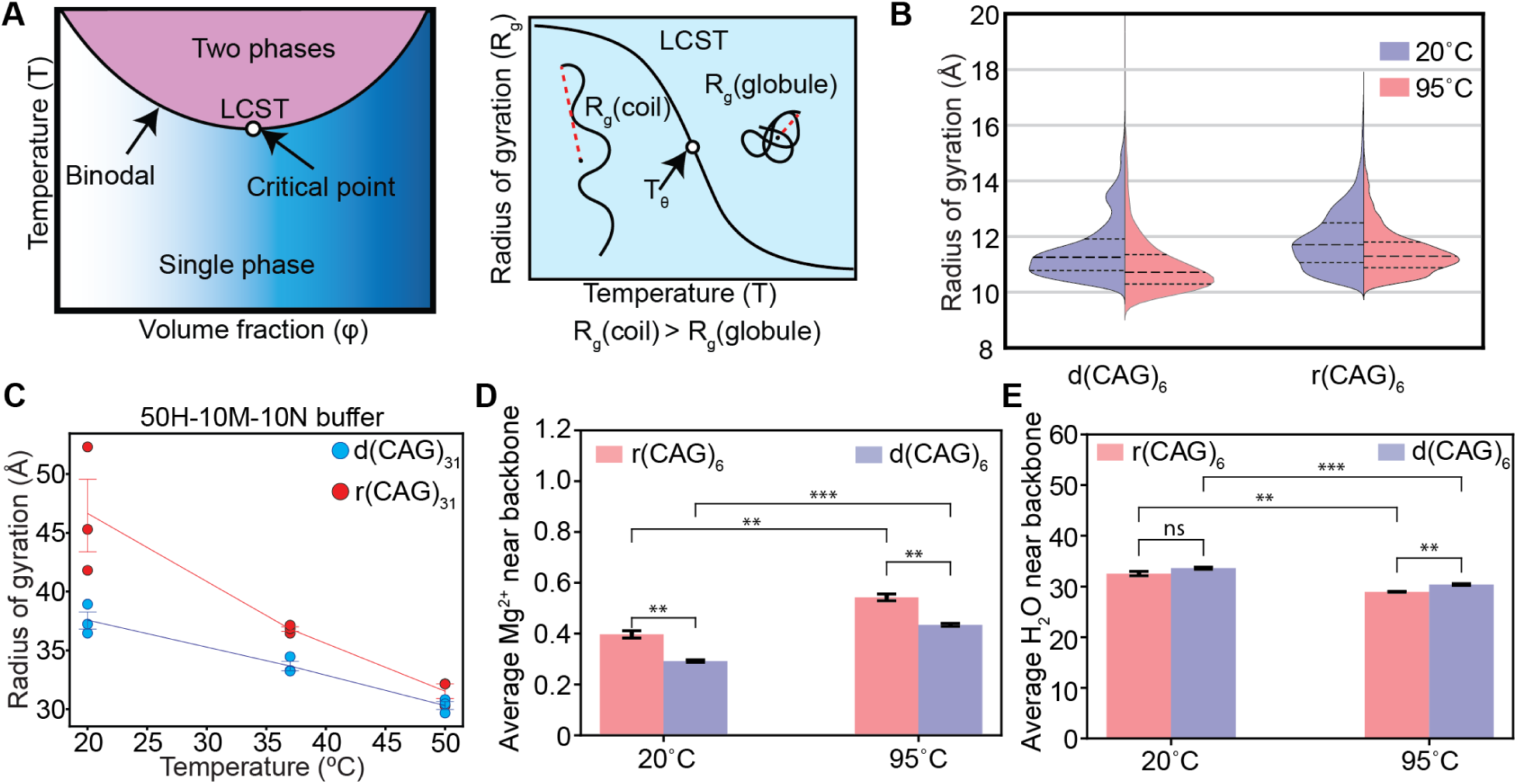
Simulations and Small Angle x-ray Scattering (SAXS) show ssDNA is more compact and interacts less with magnesium ions than RNA. **(A)** Lower Critical Solution Temperature (LCST)-type phase separation accompanies a corresponding coil-to-globule transition of the single chain upon heating. A schematic showing that as the solution temperature approaches the binodal (left), there is an associated collapse of single chains (right). **(B)** A violin plot of the distribution of the radius of gyration (*R*_g_) determined via all-atom molecular dynamics (MD) simulations for **r(CAG)_6_** and **d(CAG)_6_** at two different temperatures, 20°C (blue) and 95°C (red) in a box containing the equivalent of 150 mM MgCl_2_. The mean and upper and lower quartiles are shown as dashed lines in the distributions. We determine that the persistence lengths L_p_ for these RNA and ssDNA at 20°C are 6.42 ± 0.53 Å and 5.47 ± 0.36 Å respectively and at 95°C are 5.64 ± 0.47 Å and 4.69 ± 0.03 Å respectively. **(C)** Temperature dependence of *R*_g_ predicted by Guinier fits of SAXS data for **(CAG)_31_** ssDNA (blue) and RNA (red). **(D)** The average number of Mg^2+^ ions located in the 1^st^ and 2^nd^ solvation shells, i.e. within 5 Å of the phosphate oxygen (Op) in simulations at two temperatures, 20°C and 95°C, for r(CAG)_6_ (red bars) and d(CAG)_6_ (blue bars). P-values from left to right are p1 = 0.00213, p2 = 0.00166, p3 = 0.00004, p4 = 0.00163. **(E)** The average number of H_2_O molecules located in the 1^st^ and 2^nd^ solvation shells, i.e. within 5 Å of the phosphate oxygen (Op) in simulations at two temperatures, 20°C and 95°C, for r(CAG)_6_ (red bars) and d(CAG)_6_ (blue bars). P-values from left to right are p1 = 0.09721, p2 = 0.00130, p3 = 0.00032, p4 = 0.00119. Buffer notation used: the number in front of “H” indicates the [HEPES] in mM, the number in front of “M” indicates the [MgCl_2_] in mM, and the number in front of “N” indicates the [NaCl] in mM for each buffer. Error bars represent s.e.m. for n = 3 replicates.

## Deoxyribose suppresses the phase separation and percolation transition of CAG repeats

The interactions between nucleic acids and ions play a critical role in screening the negative charge of the phosphate backbone ^51^ as well as facilitating the desolvation of the backbone upon heating ^19^ leading to LCST-type phase separation ^19–21^. To uncover the role of the ribose sugar in thermoresponsive phase behavior, we compared r(CAG)_31_ and d(CAG)_31_ in otherwise identical conditions using temperature-controlled optical microscopy (**Figure 2.A**). While both systems displayed LCST-type phase separation, r(CAG)_31_ demonstrated systematically reduced lower cloud point temperatures (LCPTs) compared to d(CAG)_31_ by approximately 10°C through titration of both nucleic acid and Mg^2+^ ion concentrations (**Figure 2.B, C**). A further distinction between RNA and ssDNA emerges when we compare the propensity for percolation driven by phase separation. r(CAG)_31_ readily formed percolated condensates above a threshold concentration of MgCl_2_ (**Supplementary Figure S6**) while d(CAG)_31_ condensates remained reversible even at substantially higher concentrations of nucleic acid and Mg^2+^ ions (**Figure 2.D, E; Movies 1, 2**). These results indicate that while the percolation line overlaps with the two-phase coexistence line (i.e., binodal line) for RNA, it stays well-separated for ssDNA (**Figure 2.F**). The observed differences suggest a vital role in the 2’-OH group of ribose sugar in tuning both phase separation and percolation.

**Figure 2.**
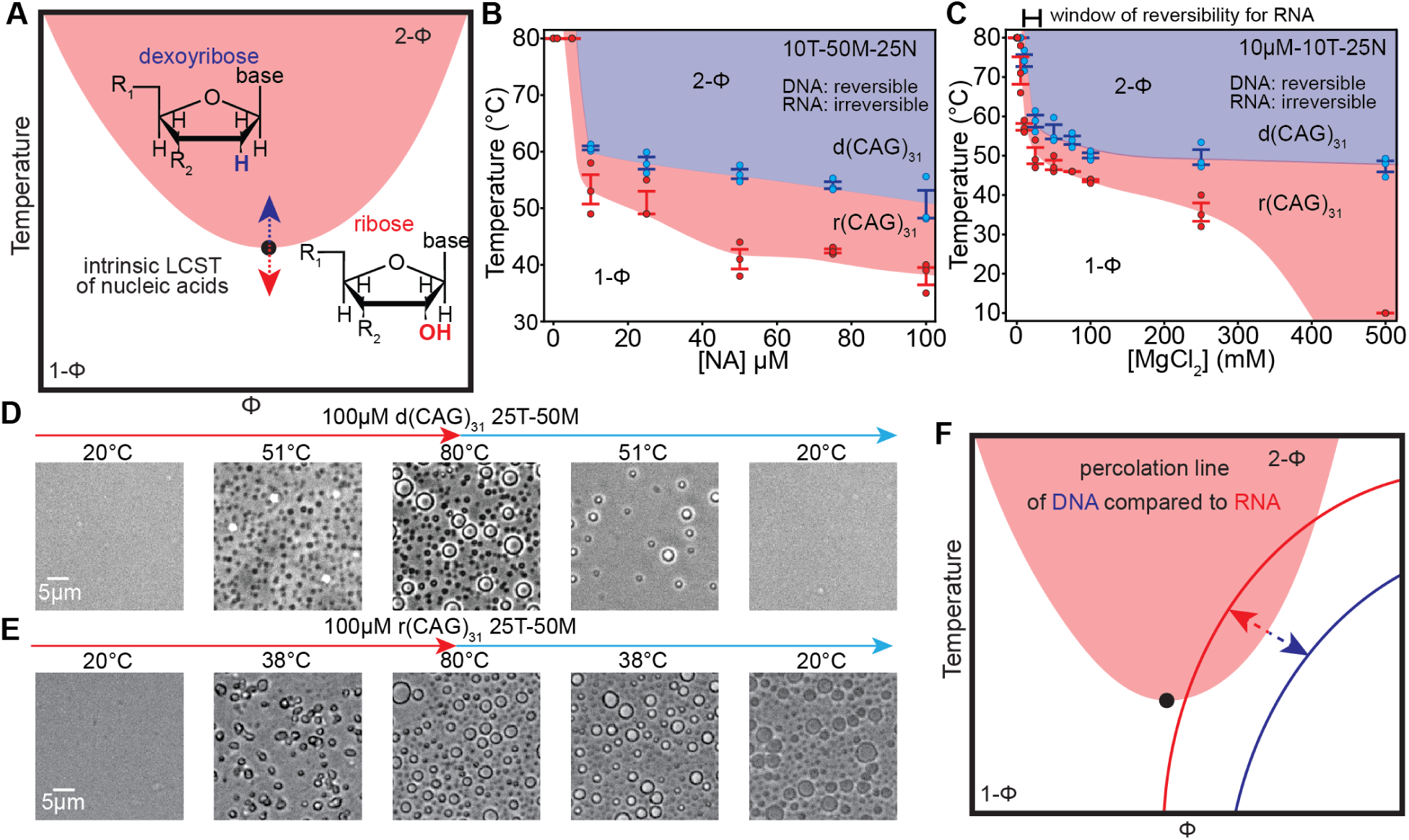
Thermoresponsive phase behavior of (CAG)_31_ repeat nucleic acids reveals distinct percolation coupled to phase separation behavior of RNA and ssDNA. **(A)** A schematic describing the contribution of the 2’-OH group of ribose to the intrinsic phase behavior of nucleic acids (NAs). Φ represents the volume fraction. 1-Φ and 2-Φ designate the single-phase and two-phase regimes. **(B)** A state diagram comparing the [NA] dependence of lower cloud point temperatures (LCPTs) for r(CAG)_31_ to d(CAG)_31_. The shaded region indicates the two phase (2-Φ) regime for r(CAG)_31_ (red) and d(CAG)_31_ (blue). **(C)** A state diagram comparing the [MgCl_2_] dependence of LCPTs for r(CAG)_31_ to d(CAG)_31_. The shaded region indicates the 2-Φ regime for r(CAG)_31_ (red) and d(CAG)_31_ (blue). r(CAG)_31_ data in **B** and **C** is reproduced for comparison purposes from reference ^19^. **(D)** Images of 100 µM d(CAG)_31_ in a buffer containing 10 mM Tris-HCl, pH 7.5 at RT, 50 mM MgCl_2_, and 25 mM NaCl. The LCPT is 50.1± 2.5°C. **(E)** Images of 100 µM r(CAG)_31_ in the same buffer as (D). The LCPT is 39.4 ± 1.5°C. **(F)** A schematic depicting the observed intersection (or a lack thereof) between the percolation line and the 2-Φ regime for DNA (blue) and RNA (red). Buffer notation used: the number in front of “T” indicates the [Tris-HCl] in mM, the number in front of “M” indicates the [MgCl_2_] in mM, and the number in front of “N” indicates the [NaCl] in mM for each buffer. Error bars represent s.e.m. for n = 3 replicates.

The percolation threshold is not only sequence-specific but also varies in a nucleic acid length-dependent manner. This argument is based on our observation that while (CAG)_n_ RNA formed percolated condensates at both n = 20 and n = 31 repeats, only r(CUG)_47_ but not r(CUG)_31_ displayed intra-condensate RNA percolation under identical experimental conditions (**Supplementary Figure S7**). We hypothesized that this may be due to chain stiffness and extracted persistence lengths, L_p_, from simulations of r(CUG)_6_ (**Supplementary Figure S7A**) and found that the rank order of L_p_ is r(CAG)_6_ > r(CUG)_6_ > d(CAG)_6_. Therefore, we next tested whether ssDNA condensate percolation can be induced through increasing repeat length. However, condensates formed by d(CAG)_n_ remained fully reversible at repeat lengths n = 20, 31, and 47 (**Supplementary Figure S8, Movies 2-4)**. These results further suggest that *c*_prc_ is higher and *T*_prc_ is lower than the binodal line for d(CAG)_n_, making the percolation regime non-overlapping with the 2-phase (2-Φ) regime (**Figure 2.F**) at the tested chain lengths.

## Systematic variations in CG-rich sequences show RNA universally phase separates with a greater propensity than ssDNA

It has been proposed ^19,22,33^ that the tendency of CG-rich RNA to phase separate into irreversible condensates may play a major role in the intra-cellular foci formation by the CAG and CUG repeat RNAs ^33^ that are linked to Huntington’s disease and myotonic dystrophy, respectively, and by the CGG repeats that are found in Friedrich’s ataxia ^52^. Based on our experimental results on CAG-repeat nucleic acids and all-atom simulations, we hypothesize that RNA in general has reduced LCPTs and percolation propensity compared to ssDNA due to the increased inner sphere contacts with magnesium ions in the phosphate backbone and the 2’-OH of the ribose sugar (**Figure 1.D; Supplementary Figures S5**). To systematically test this, we varied the middle base in our CG-rich sequence, C<u>X</u>G_31_ (<u>X</u>= <u>A</u>, <u>T/U</u>, <u>G</u>, <u>C</u>). We find that similar to r(CAG)_31_ compared with d(CAG)_31,_ RNA has reduced LCPTs for all four repeat sequences (**Figure 3.A.C)**. r(CUG)_31_ has LCPTs approximately 15°C lower compared to d(CTG)_31_ (**Figure 3.D.E; Movies 5-8**), which is the largest difference of the set and may indicate an important role for the methyl group in thymine in this process. (CCG)_31_ and (CGG)_31_ also show reduced LCPTs for RNA compared to ssDNA (**Figure 3.A, Movies 5-12**). A significant difference, however, between d(C<u>G</u>G)_31_ and d(C<u>C</u>G)_31_ condensates is that the former percolates throughout the concentration range tested while d(C<u>C</u>G)_31_ condensates remain reversible similar to r(CAG)_31_ versus d(CAG)_31_ (**Figure 3.F.I; Movies 9-12**). The irreversible condensation of d(CGG)_31_ suggests that, while the stacking contributions facilitate percolation in RNA via purines (A and G), G-tracts specifically play a primary role in facilitating percolation in ssDNA condensates.

**Figure 3.**
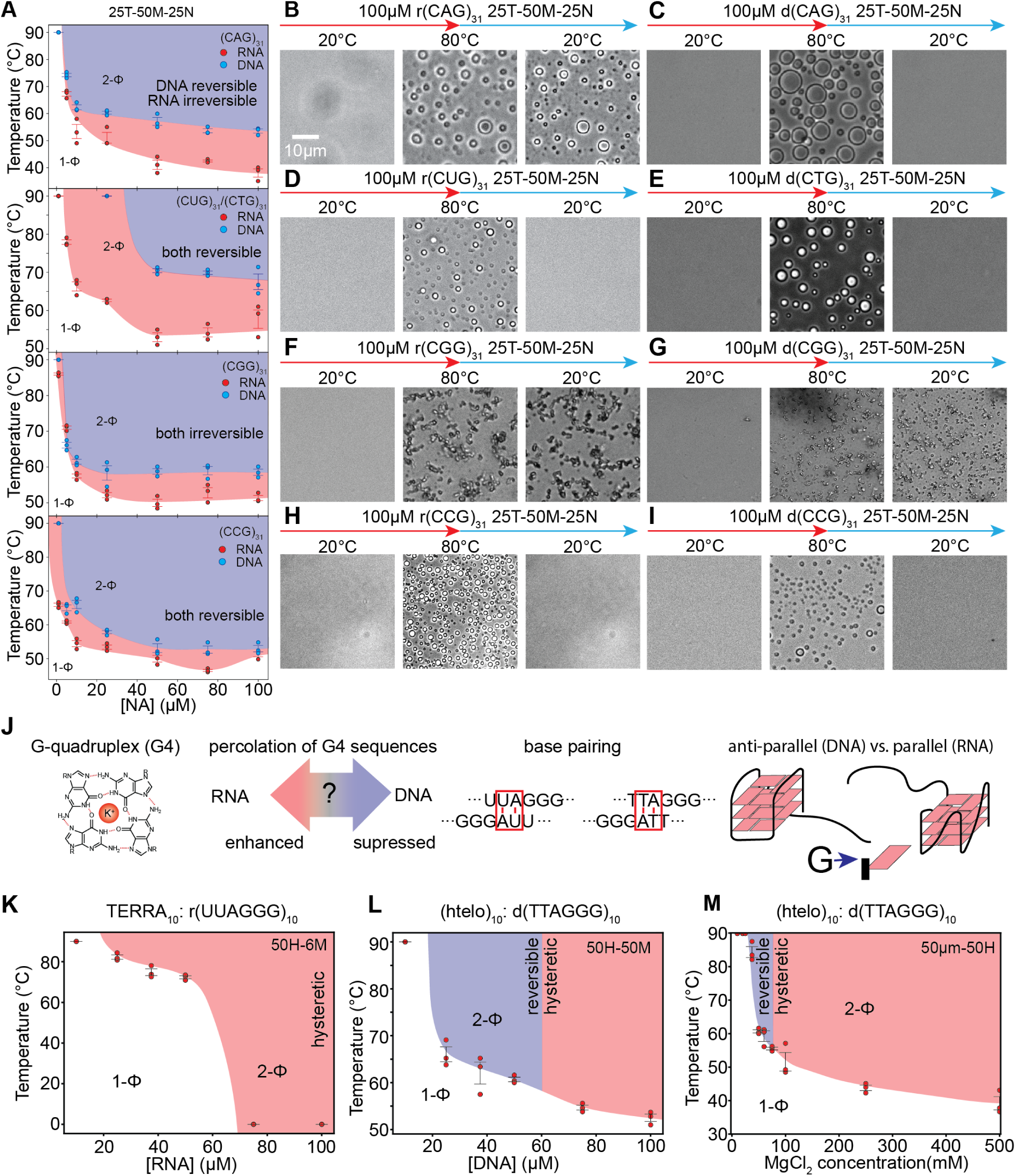
Phase separation and percolation transitions of nucleic acids (NAs) depend on their primary sequence and secondary structure. **(A)** State diagrams comparing the [NA] dependence of LCPTs for r(CAG)_31_ to d(CAG)_31_, r(CUG)_31_ to d(CTG)_31_, r(CGG)_31_ to d(CGG)_31_, and r(CCG)_31_ to d(CCG)_31_. The shaded regions indicate the phase separated (2-Φ) regime for RNA (red) and DNA (blue). All samples are shown in a buffer containing 25 mM Tris-HCl, pH 7.5 at RT, 50 mM MgCl_2_, and 25 mM NaCl. r(CAG)_31_ data is reproduced for comparison purposes from reference ^19^. **(B-I)** representative images of 100 µM nucleic acid in the order r(CUG)_31_, d(CTG)_31_, r(CGG)_31_, d(CGG)_31_, r(CCG)_31_, d(CCG)_31_, r(CAG)_31_, and d(CAG)_31_ with LCPTs in the same order, 39.4 ± 1.5°C, 50.1± 2.5°C, 59.2 ± 2.4°C, 67.5 ± 2.0°C, 50.5 ± 0.73°C, 58.5 ± 0.87°C, 51.8 ± 0.74°C, and 54.8 ± 0.96°C. All samples were prepared in the same buffer as in (A). The scale bar is identical for all panels. **(J)** A schematic showing the sequence features and base pairing potentials for TERRA and human telomeric DNA, htelo. Also shown is a schematic of a G-quadruplex in the anti-parallel and parallel configurations for DNA and RNA respectively. **(K)** A state diagram showing the [RNA] dependence of TERRA phase behavior in a buffer containing 50 mM HEPES, pH 7.5 at RT, and 6 mM MgCl_2_. **(L)** A state diagram showing the [DNA] dependence of htelo phase behavior in a buffer containing 50 mM HEPES, pH 7.5 at RT, and 50 mM MgCl_2_. The shading indicates reversibility (blue) or irreversibility (red) of the 2-Φ regime. **(M)** A state diagram showing the [MgCl_2_] dependence of htelo phase behavior in a buffer containing 50 µM (htelo)_10_ and 50 mM HEPES, pH 7.5 at RT. The shading indicates reversible condensation (blue) or irreversible percolation (red). Buffer notation used: the number in front of “T” indicates the [Tris-HCl] in mM, the number in front of “H” indicates the [HEPES] in mM, the number in front of “M” indicates the [MgCl_2_] in mM, and the number in front of “N” indicates the [NaCl] in mM for each buffer. Error bars represent s.e.m. for n = 3 replicates.

A further question arises whether this behavior is special to CG-rich sequences with patterned base pairing such as (C<u>X</u>G)_n_ and whether a random arrangement would also show the same trend for RNA versus ssDNA. We designed a scrambled (CAG)_31_ equivalent sequence (**Supplementary Table 1**), rsc(CAG)_31,_ and observed that this RNA had systematically higher LCPTs than an otherwise equivalent ssDNA, dsc(CAG)_31_ (**Supplementary Figure S9)**. This result supports the idea that the tendency of RNA to phase separate is likely generally higher than their ssDNA equivalents.

## G-quadruplex forming sequences promote networking transitions in both RNA and ssDNA

G-rich sequences are well known to form non-canonical nucleic acid structures known as G-quadruplexes (dG4s and rG4s) ^53–57^. Our observations that only (CGG)_31_ repeats form percolated condensates for ssDNA led us to hypothesize that G-tracts and potentially G4 structures may play a dominant role in percolation. We note that CCG and CGG repeats have similar degrees of base pairing and similar *T*_phase_ with the primary difference that only CGG has the possibility of forming G-quadruplex structures (**Figure 3.J**). Therefore, in addition to (CGG)_31_, we chose to examine four other G4 forming sequences that have well-established biological relevance, telomere derived RNA [TERRA; r(UUAGGG)_n_] and telomeric DNA [htelo; d(TTAGGG)_n_] (**Figure 3.J.M; Supplementary Figures S10-S11**) as well as (GGGGCC)_n_ (**Supplementary Figure S12**) which is linked to C9ORF72 Amyotrophic lateral sclerosis (ALS) and frontotemporal dementia (FTD) ^58^. We tested the structure of these nucleic acids in the single-phase regime through circular dichroism (CD) spectroscopy, which revealed the formation of G-quadruplexes for all putative dG4s and rG4s to various degrees (**Supplementary Figure S13)**. For (TERRA)_10_ and (htelo)_10_, we find both have LCST-type phase separation with significant differences in Mg^2+^ sensitivity and percolation threshold with the ssDNA displaying lower driving force for condensation under all conditions (**Figure 3.K.M; Supplementary Figure S10; Movies 13-16)**. The thermoresponsive transition of (TERRA)_10_ is extremely sharp going from no phase separation between 5-to-90 °C to percolated condensates within the range of 6 mM to 7 mM MgCl_2_ ^40^ (**Supplementary Figure S10.A**). Moreover, at a fixed MgCl_2_ concentration (6 mM), percolation (irreversible condensation) is always observed for (TERRA)_10_ (**Figure 3.K; Supplementary Figure S11**). Conversely, (htelo)_10_ form reversible condensates under identical buffer conditions, nucleic acid concentrations, and salinity (**Figure 3.L**). Percolation was observed only above 50 µM (htelo)_10_ and at an order of magnitude higher MgCl_2_ concentration (**Figure 3.M; Supplementary Figures S10.C-D; Movie 15**). In the case of (GGGGCC)_5_, we observe that RNA phase separates at lower temperatures than the corresponding ssDNA, however, both are very sensitive to Mg^2+^ ion concentration and form irreversible fractal clusters throughout the temperature range for [Mg^2+^] > 20 mM (**Supplementary Figure S12**). Additionally, we used Thioflavin T (ThT) as a reporter dye to probe the extent of intra-condensate G-quadruplex formation ^59–61^. We observed that condensates formed by putative G-quadruplex forming RNA and ssDNA display substantially higher ThT signal while condensates formed by r(CAG)_31_ and (UUAGUG)_10_, which do not form G-quadruplexes, remained largely ThT negative (**Supplementary Figure S14)**. Extending ThT measurements to other condensates formed by putative G-quadruplex RNA or ssDNA also revealed ThT positivity, albeit the absolute ThT signal varied between different condensates, suggesting variations in the nanoscale condensate structures.

## Post-transcriptional modifications tune the phase behavior of RNA

The 2’-OH group on the ribose sugar is a site of abundant chemical modifications in RNA ^62,63^. Modifications in the 2’-OH group on the ribose sugar can lead to altered nucleic acid conformation ^64^. Our results on the differential phase behavior of RNA and ssDNA provide a testable hypothesis that 2’-OH modifications may tune RNA condensation and reversibility of this process (**Figure 4.A**). One common RNA modification is the methylation of the 2’-OH group on the ribose sugar (2’-O-Me), which may effectively silence the chemical activity of the 2’-OH group ^65^. In addition, 2’-O-Me can bias sugar puckering towards the C3’-*endo* conformation found in the A-form DNA and RNA instead of the B-form helix which adopts a C2’-*endo* conformation ^66^. Modifications to this site which change the conformation of the sugar ^67^ may also perturb the local hydration of the backbone ^68^. Because of the dominant role of desolvation underlying the LCST-type nucleic acid phase separation and the potential for ion-mediated crosslinking interactions with the 2’-OH group of ribose but not the 2’-H of deoxyribose, we hypothesized that 2’-O-Me is a tunable switch to regulate the phase behavior and percolation transitions of RNA.

**Fig. 4.**
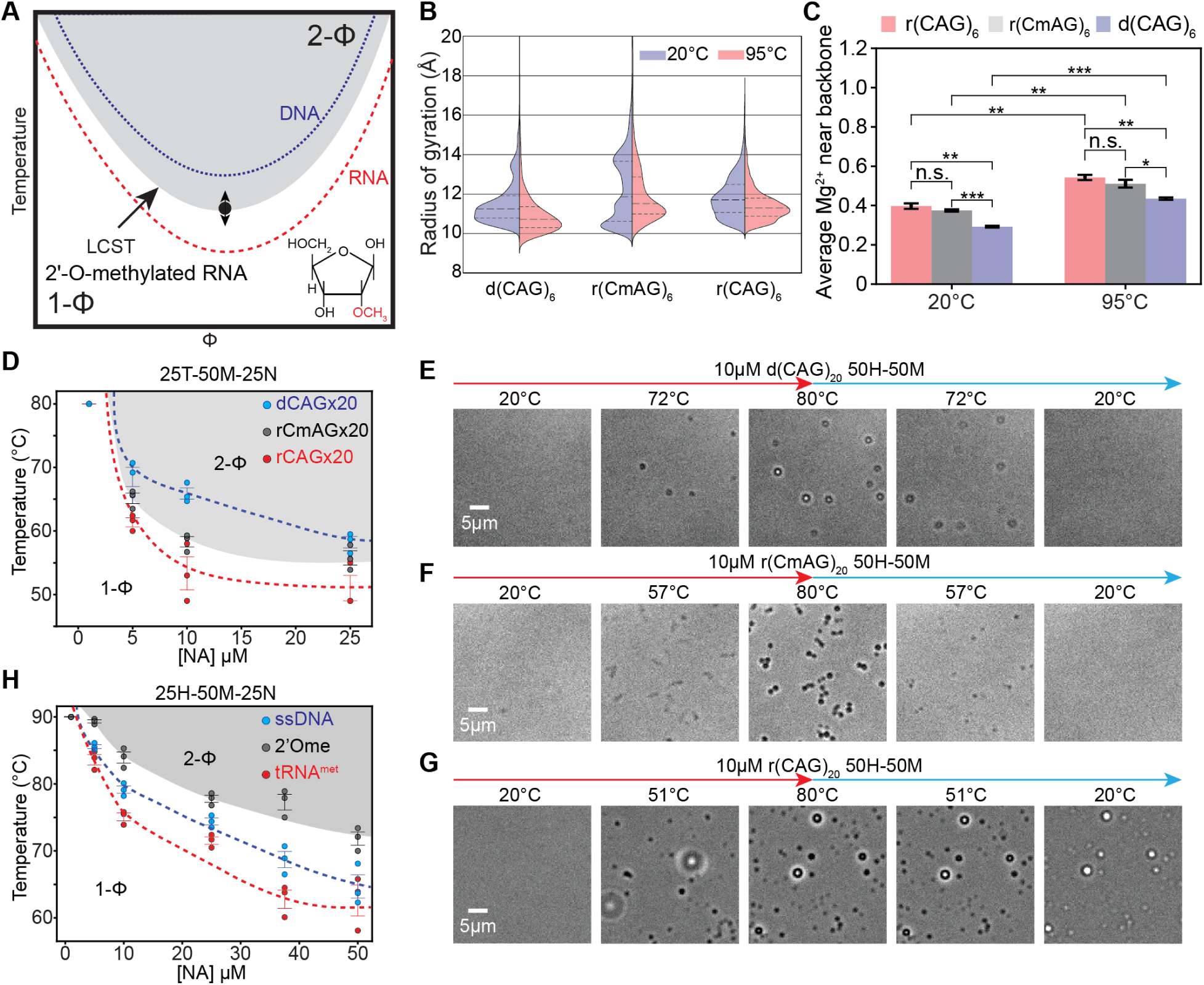
Methylation of the 2’-OH group on the ribose sugar (2’-O-Me) tunes percolation coupled to phase separation of RNA. **(A)** A schematic illustrates the modulation of the intrinsic phase separation of RNA versus ssDNA through methylation of the 2’-OH group of ribose sugar. **(B)** A violin plot of the distribution of the radius of gyration determined via simulations for r(CmAG)_6_, r(CAG)_6_, and d(CAG)_6_ at two different temperatures, 20°C (blue) and 95°C (red). Here mA refers to 2’-O-Me sugar modification in adenosine. We determine that the persistence length L_p_ for the r(CmAG)_6_ at 20°C is 5.76 ± 0.25 Å and at 95°C is 5.62 ± 0.30 Å. **(C)** The average number of Mg^2+^ located in the 1^st^ and 2^nd^ solvation shells, i.e. within 5 Å of the phosphate oxygen (Op) in simulations at two temperatures, 20°C and 95°C, for r(CAG)_6_ (red bars), r(CmAG)_6_ (gray bars), and d(CAG)_6_ (blue bars). P-values from left to right are 0.21528, 0.00213, 0.00029, 0.00166, 0.00252, 0.24997, 0.00163, and 0.01960. **(D)** A state diagram comparing the [NA] dependence of LCPTs for r(CmAG)_20_ to r(CAG)_20_ and d(CAG)_20_. The grey shaded region indicates the 2-Φ regime for r(CmAG)_20_. **(E)** Images of 10 µM d(CAG)_20_ in a buffer containing 50 mM HEPES, pH 7.5 at RT. The LCPT is 71.4 ± 0.87°C. **(F)** Images of 10 µM r(CmAG)_20_ in the same buffer as (E). The LCPT is 57.7 ± 0.81°C. **(G)** Images of 10 µM r(CAG)_20_ in the same buffer as (E). The LCPT is 51.4 ± 2.49°C. **(H)** A state diagram comparing the concentration dependence of LCPTs for a tRNA^met^ sequence derived from *Sulfolobus acidocaldarius* is shown. The phase behavior of RNA, ssDNA, and a modified RNA with the four native sites of 2’-O-methylation was compared in a buffer containing 25 mm HEPES, pH 7.5 at RT, 50 mM MgCl_2_, and 25 mM NaCl. Corresponding microscopy images are shown in **Supplementary Figure S16**. Buffer notation used: the number in front of “H” indicates the [HEPES] in mM, the number in front of “M” indicates the [MgCl_2_] in mM, and the number in front of “N” indicates the [NaCl] in mM for each buffer. Error bars represent s.e.m. for n = 3 replicates.

To test this idea, we designed a 2’-O-Me modified RNA r(CmAG)_n_ which we expected would best preserve the stability of C:G base pairing interactions while modulating the backbone hydration around the 2’-O-methylated Adenine (mA). All-atom MD simulations of r(CmAG)_6_ show that the chain has an *R*_g_ value that is multimodal and more widely distributed than that of ssDNA and RNA (**Figure 4.B**). Additionally, we observe that the probability of having inner sphere contacts with Mg^2+^ ions is significantly reduced compared to unmodified RNA (**Figure 4.C; Supplementary Figure S15.A**) accompanied by an increase in inner sphere water contacts (**Supplementary Figure S15.B-C**). Furthermore, we find that the probability of Mg^2+^ ions making contacts with the 2’-OH is greater than with 2’-Ome or 2’-H (**Supplementary Figure15.D-E**). Using temperature-dependent microscopy of r(CmAG)_20_, we observe that across the full range of concentrations, r(CmAG)_20_ demonstrated LCPTs that were systematically between those of d(CAG)_20_ and r(CAG)_20_ with the RNA having the lowest and DNA having the highest LCPTs (**Figure 4.D**). In some combinations of [NA] and [MgCl_2_], r(CmAG)_20_ displays reversible condensation like ssDNA (**Figure 4.E, F; Movies 3, 17**) while in other conditions, it percolates like RNA (**Figure 4.G; Supplementary Figure S16; Movie 18**).

In order to determine the effects of 2’-O-methylation in sequence positions and numbers that are biologically relevant, we tested the phase behavior of the tRNA for methionine (tRNA^met^) derived from the extremophilic archaeal bacteria Sulfolobus acidocaldarius that has four sites of 2’-Omethylation ^69^. This is a considerably smaller fraction than the number of modifications we introduced in r(CmAG)_20_. We compare an unmodified tRNA and a ssDNA of the same sequence to the methylated tRNA^met^ (**Figure 4.H**). We observe that this tRNA also undergoes LCST-type phase separation with the rank order of cloud point temperatures being RNA < ssDNA < 2’-Ome RNA under identical experimental conditions. Of note, the ssDNA has the intermediate LCPT, which may be due to the effect of 2’-Ome modification on the stability of the tertiary structure of the RNA ^66^. Combining the single chain properties from simulations with the bulk phase separation measurements, our results suggest that 2’-OMe weakens the driving force for RNA phase separation and percolation.

## The 2’-OH group of RNA facilitates phase separation and percolation via ion-mediated interactions

One consistent observation between RNA and ssDNA is that increased concentrations of magnesium ions lower *T*_phase_ and can induce intra-condensate RNA percolation **(Figures 2.C. and 3.L, M; Supplementary Figures S6-7)**. One possible mechanism is that these ions facilitate both crosslinking and charge neutralization of the backbone ^24,25,28,29,31,70^. The 2’-OH group may also interact with the divalent ions to influence the stiffness of the backbone ^71^. If true, replacing Mg^2+^ with other divalent cations such as Ca^2+^ may influence RNA but not ssDNA phase behavior substantially (**Figure 5.A**). Both ions are kosmotropes ^72^ meaning they possess well-ordered water shells ^73^. However, Mg^2+^ has a smaller ionic radius ^71^ and thus possesses a more ordered water shell than Ca^2+^ (**Figure 5.A; Supplementary Figure S18**). We hypothesize that the decreased stability of the water shell around Ca^2+^ should facilitate LCST-type phase behavior due to the lower energy cost to liberate the water around Ca^2+^ upon heating. To test this, we systematically compare the ion concentration-dependent nucleic acid phase behavior for Ca^2+^ and Mg^2+^. We find that at all concentrations of r(CAG)_20_ or r(CmAG)_20_, Ca^2+^ facilitates percolation coupled to phase separation better than Mg^2+^ ions (**Figure 5.B, C; Supplementary Figure S19.A**). In contrast, we see modest difference in the thermoresponsive phase behavior of d(CAG)_20_ through the same comparison (**Supplementary Figure S19.B**). This is recapitulated for r(CAG)_31_ versus d(CAG)_31_ where RNA shows dramatic differences in condensation behavior when Mg^2+^ is replaced by Ca^2+^, while ssDNA does not (**Supplementary Figure S19.C, D; Movie 19**). Furthermore, we find this trend holds even at 500 mM Ca^2+^, (**Supplementary Figure S19.E-F**). These data collectively confirm that interactions between the 2’-OH and divalent ions are important for tuning phase separation and percolation behavior in RNA.

**Fig. 5.**
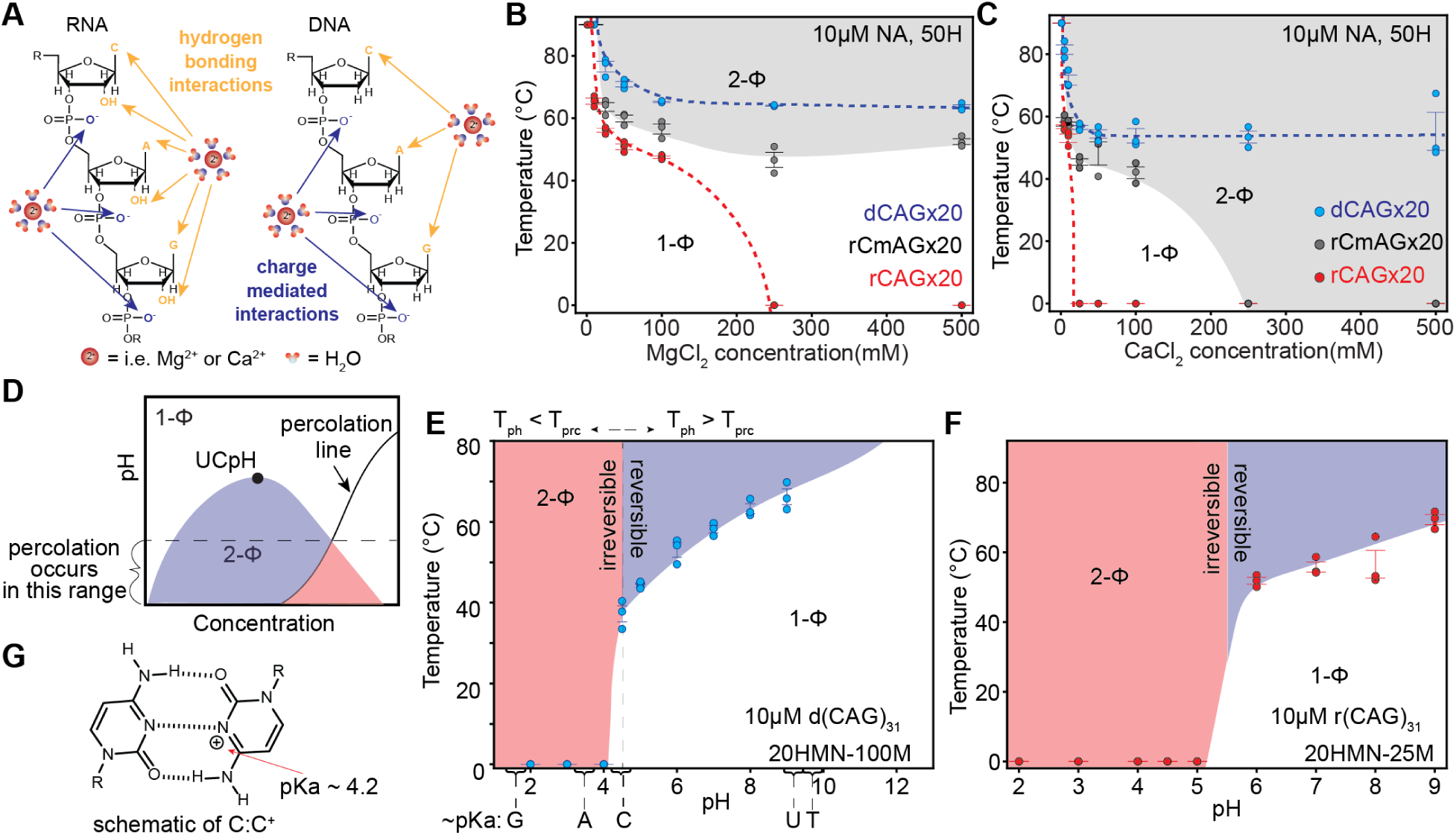
RNA and ssDNA condensation have differential responses to ionic conditions and solution pH. **(A)** A Schematic summarizing the ionic interactions in RNA and ssDNA. **(B)** A state diagram comparing the [MgCl_2_] dependence of the LCPTs for r(CmAG)_20_ to r(CAG)_20_ and d(CAG)_20_. The grey shaded region indicates the 2-Φ regime for r(CmAG)_20_. **(C)** A state diagram comparing the [CaCl_2_] dependence of the LCPTs for r(CmAG)_20_ to r(CAG)_20_ and d(CAG)20. The grey shaded region indicates the 2-Φ regime for r(CmAG)_20_. **(D)** A schematic of the relationship between an Upper Critical pH (UCpH) and the percolation line. Shading indicates the reversible (blue) versus irreversible (red) condensation regimes. **(E)** A state diagram demonstrating a switch from reversible to irreversible condensation upon decreasing the pH of the system for d(CAG)_31_. Shading indicates the reversible (blue) versus irreversible (red) regimes. **(F)** A state diagram demonstrating the onset of irreversible condensation upon decreasing the pH of the system for r(CAG)_31_. Shading indicates the reversible (blue) versus irreversible (red) regimes. Buffer notation used: the number in front of “H” indicates [HEPES], the number in front of “HMN” indicates the [HEPES]; [MES]; and [NaAc] in mM, and the number in front of “M” indicates [MgCl_2_] in mM in each buffer. Error bars represent s.e.m. for n = 3 replicates.

## Differences in single-chain stiffness may explain observed variations in the percolation behaviors of RNA and ssDNA condensates

In addition to ion-mediated interactions, we also hypothesize that differences in chain stiffness can regulate chain percolation in the condensed phase. It was shown previously that r(U)_40_ is stiffer than d(T)_40_ through a combination of single-molecule fluorescence and SAXS measurements ^74^ comparing their response to Mg^2+^ titration. We observe consistently larger *R*_g_ for r(CAG)_6_ compared to d(CAG)_6_ in MD simulations (**Figure 1; Supplementary Figure S5, S15**) and via SAXS (**Supplementary Figures S2-3**). Chain stiffness of nucleic acids has also been correlated with a number of factors in biological settings including nucleosome positioning ^75,76^, transcription factor affinity ^77^, and interactions with the ribosome ^78^. Based upon our observation that chain stiffness varies between r(CAG)_6_ and r(CUG)_6_ where the former is the stronger percolating RNA, we posit that chain stiffness may influence the percolation propensity of nucleic acids. We find that chain stiffness decreases in the order r(CAG)_6_ > r(CmAG)_6_ ∼ r(CUG)_6_ > d(CAG)_6_ which also corresponds to the rank order of percolation propensity determined experimentally (**Figure 1.B**, **Supplementary Figure S20.B**).

It has been shown that the variation in stiffness of nucleic acids can regulate the stability of secondary structures and can result in different folding behavior ^79,80^. We hypothesized that differences in chain stiffness, at least in part, may result in the observed disparities in percolation behavior for RNA and ssDNA condensates. To test this, we designed a coarse-grained model for a generic semiflexible polymer. Here, we employed an AB-type coarse-grained model to mimic the effect of base pairing and stacking in the CAG systems, where A–B can form favorable crosslinks (analogous to C–G base pairing). This minimal model allows us to isolate the effect due to stiffness (*κ*) and directly probe orderness (*S*), morphology, and number of intermolecular crosslinks (*N*) within condensates (**Supplementary Figure S20.C**). For low chain stiffness (*κ* ≤ 6), condensates exhibit low structural ordering (*S*) and low shape anisotropy (*A*), resembling behaviors of liquid-like condensates. In contrast, at higher chain stiffness (*κ ≥* 8), the droplets become more ordered and depict less spherical shapes. At intermediate stiffness we observe a sharp transition in condensate properties, implying that there is a phase boundary where a slight increase in chain stiffness leads to the emergence of dynamically arrested states. At the sub-molecular level, the emergence of dynamically arrested states corresponds to an increase in intermolecular crosslinks (**Supplementary Figure S20**). Collectively, these results suggest that differences in single-chain stiffness between RNA and DNA can also strongly contribute to their varied percolation in condensates.

## RNA and ssDNA display an upper critical pH (UCpH)

Similar to a UCST or an LCST, phase separation can be defined by an upper or lower critical pH (UCpH/LCpH) where the phase behavior is determined by the relationship between the pH, pK_a_ of associative sites, and the isoelectric point ^48,81^. We hypothesize that the aggregates that sediment as pH is lowered for nucleic acids ^82^ in the presence of divalent ions may stem from a UCpH-type percolation coupled to phase separation ^48^ (**Figure 5.D)**. To test this, we take r(CAG)_31_ and d(CAG)_31_ at conditions where they undergo reversible condensation at physiological pH (7.5) and fix all experimental parameters except pH. Using temperature-controlled microscopy, we observe that variation of pH lowers the *T*_phase_ for r(CAG)_31_ until between pH 5 and pH 6, at which point the condensates switch from being reversible to irreversible implicating the occurrence of percolation (**Figure 5.E**). At pH 5, RNA forms irreversible clusters immediately upon mixing, which persist at all experimentally accessible temperatures. Likewise, for d(CAG)_31_, we observe that phase separation becomes irreversible at pH 4.5 with irregular clusters forming at room temperature at pH 4 (**Figure 5.F)**.

The pH of the solution can modify the electrostatic properties of nucleic acids through the protonation reaction of the backbone, the sugar, and/or the nucleobases; however, within the range of our buffer pH only some of the pK_a_’s are relevant ^83^. We hypothesize that this pH dependent switch from phase separation to percolation may be driven by the protonation of cytosine, pK_a_ ∼ 4.2, and the phosphate groups, pK_a2_ ∼ 6-7 (**Supplementary Figure S21.A; Movies 20-22**) ^83^. To test if the percolation is driven by the cytosine protonation, we studied the phase behavior of polycytidylic acid and polyuradylic acid, poly(rC) and poly(rU) respectively. The pK_a_ of uracil, 9.2, lies outside the pH range of our experiments. We find that poly(rU) condensate cloud point temperatures do not vary substantially with the change in buffer pH from 7.5 to 3.5 (**Supplementary Figure S21.B; Movie 23**). These results suggest that pH-mediated percolation of CAG-repeat nucleic acid condensates did not arise from the protonation of the phosphate groups. Intriguingly, although poly(rU) condensate did not register any significant changes, poly(rC) condensates were observed to sharply transition from reversible to irreversible between pH 5.5 and 5.75 indicating the onset of intra-condensate percolation through the protonation of cytosine (**Supplementary Figure S21.C**). The conversion of C to C^+^ at lower pH has been reported to form i-motif structures comprised of intercalated strands with C-C^+^ base pairing ^84,85^, which may be a driver of percolation. These findings indicate the potential for the engineered design of phase separation and percolation in nucleic acid condensates based on sequence-specific ionization.

## Summary

Through experiments supported by insights obtained from atomistic simulations, we show that despite their similar charge and chemistry, the 2’-OH group of ribose sugar in RNA encodes substantial differences in phase behavior compared to ssDNA. It has been long known that individual chains of DNA and RNA have distinct conformational properties. We reveal that although ssDNA chains are more compact, RNA has a greater tendency to form condensates (**Figures 1-2**). This runs counter to the intuition derived from extensive experiments and simulations on disordered protein systems showing that single-chain compaction is a good proxy for the phase separation driving force ^12,36,44,86–88^. Our observation is consistent with prior reports for protein systems that display LCST-type behavior such as elastin-like polymers, which do not demonstrate a clear correlation ^47^ underscoring the need for further studies on the link between single-chain properties of LCST-type polymers and their phase behavior. Our work also helps establish that a sequence-specific percolation transition of nucleic acid condensates can be engineered along with phase separation, leading to a programmable temperature-dependent and pH-controlled hysteretic phase behavior.

While the role of RNA modifications in determining the structural ensemble of RNA has been recognized from the perspective of folding ^89^, it remains largely unknown how the diversity of RNA modifications influences the landscape of their mesoscale biophysics ^22^. Our results with 2’-O-methylation of RNA (**Figure 4**) suggest that modifications to nucleic acids may act as a tunable switch for regulating RNA condensation and their transitions between reversible and irreversible states. 2′-O-methylation is a common PTM of ribosomal RNA (rRNA), which must undergo highly coordinated steps of synthesis, processing, and assembly in the nucleolus ^15,90^, a multiphasic biomolecular condensate in the nucleus ^13^. Our work showing modulation of RNA condensation by 2’-O-methylation may suggest that this modification could influence how rRNA behaves within the nucleolus and may help prevent excessive RNA entanglement during rRNA processing as it undergoes directional transport ^91^ from the fibrillar component (FC) and to the granular component (GC); thereby ensuring efficient ribosome assembly. It has been observed that the extent of 2’-O-methylation increases as rRNA is processed and transported out of the nucleolus ^92^. In combination with the pH gradient that exists between the nucleolar subphases ^93^, 2’-O-methylation may facilitate vectorial rRNA ejection from the nucleolus leading to testable hypotheses regarding the interplay between percolation, phase separation, pH gradient, modification status, and compartmentalization of RNA.

Ion identity is central to proper folding and function of nucleic acids. Metal ion concentrations are tightly controlled within the cellular environment ^94^. We observe that the impact of changing from Mg^2+^ to Ca^2+^ is more profound on the condensation behavior of RNA than ssDNA. Ca^2+^ ions induce robust percolation of RNA at an order of magnitude lower concentrations than Mg^2+^ ions (**Figure 5**), which can inhibit RNA function via the formation of dynamically arrested condensates. This observation raises the possibility that Ca^2+^ induced percolation coupled to phase separation may explain the previously observed inhibition of certain classes of ribozymes by Ca^2+ 95,96^ suggesting a testable relationship between ion-mediated irreversible condensation and enzymatic activity. Nucleic acid percolation is also engendered under acidic pH due to protonation of cytosines (**Figure 5**). In the cellular environment, energy depletion can lead to acidification and a glass-like phase transition in yeast cytoplasm ^97^. Our data provides a testable hypothesis that acidification-induced RNA percolation could be a driver of this observed dynamical arrest of the cytoplasm in yeast and cellular extracts ^98^.

## Acknowledgments

This work was supported by National Institutes of Health grant R35 GM138186 (PRB), Cornell High Energy Synchrotron Source, proposal 3292-B (PRB, GMT, GMW), St. Jude Research Collaborative on Biophysics of RNP granules (PRB), National Institutes of Health grant R35 GM155259 (JAJ), National Science Foundation (NSF) through the Princeton University (PCCM) Materials Research Science and Engineering Center DMR-2011750 (JAJ), and Chan Zuckerberg Initiative DAF, an advised fund of Silicon Valley Community Foundation, grant 2023-332391 (JAJ). The authors acknowledge members of the Banerjee Lab and Joseph Group for valuable discussions during various stages of the manuscript preparation. The authors would also like to acknowledge Meet Raval, Michael Pierce, Gabriel Wilber and Christopher Voskinarian for assistance with SAXS data collection. Simulations reported on in this manuscript were performed using the Princeton Research Computing resources at Princeton University, which is a consortium of groups led by the Princeton Institute for Computational Science and Engineering (PICSciE) and Office of Information Technology.

## Author contributions

Conceptualization: PRB, GMW, Methodology: PRB, GMW, JAJ, DA, GMT, Investigation: GMW, DA, Visualization: GMW, DA, Funding acquisition: PRB, JAJ, GMT, Project administration: PRB, JAJ, Supervision: PRB, JAJ, Writing – original draft: GMW, PRB, DA, JAJ, Writing – review & editing: PRB, GMW, JAJ, DA, GMT.

## Data availability

All data are included in the main text or the supplementary materials.

