## Supplementary Information for "The Critical Role of the 2’-OH group in Phase Separation and Percolation Transitions of RNA"

† Lead author for the experimental part of this work; ‡ Lead author for the simulation part of the work.

*Correspondence for the experimental part of this work should be addressed to: P.R.B; Correspondence for the simulation part of this work should be addressed J.A.J.

**The PDF file includes:**

Materials and Methods
Figs. S1 to S26
Tables S1
Supplementary Note 1 containing Figs. SN1-SN31
Movie legends S1 to S23
References

**Other Supplementary Materials for this manuscript include the following:**

Movies S1 to S23

### Materials and Methods

#### Simulations

#### Atomistic simulations of CAG repeats:

We perform atomistic molecular dynamics (MD) simulations of CAG repeats using CHARMM36m [1,2] force field with TIP3P [3] water model. CHARMM36m includes parameterizations of post-transcriptionally modified ribonucleotides [4], which aligns well with the goal of this study. For all simulations, we use GROMACS [5,6] package (Version 2023.3) on GPUs via Princeton Research Computing Clusters. Initial states of the systems are generated using CHARM-GUI [7,8]. To prepare the systems, r(CAG)$_6$, r(CmAG)$_6$, and d(CAG)$_6$ are solvated in an 80Å×80Å×80Å cubic box. $Mg^{2+}$ or $Ca^{2+}$ ions, along with $Cl^-$ ions, are added to achieve the target divalent salt concentration of 150mM (~55,000 atoms in total). We then perform energy minimization via the steepest descent method, followed by a short equilibration (500–700 ps) in the *NVT*-ensemble using V-rescale algorithm for temperature coupling ($T$ = 293 K or 368 K) to obtain different initial structures. Finally, the 600-ns production run is performed in the *NPT*-ensemble, employing V-rescale algorithm for temperature coupling and C-rescale algorithm for pressure coupling at $T$ = 293 K/368 K and $p$= 1 bar. The data from the last 500ns is used for data analysis. In all simulations, periodic boundary conditions are imposed, the PME [9] algorithm is used for long-range electrostatic interactions, and LINCS [10] algorithm is employed for H-bond constraints. Each polymer is run for three independent simulations, yielding aggregate simulation times of ca. 1,500 ns. MDAnalysis package [11] is used for all trajectory analysis, and Pymol [12] and UCSF ChimeraX [13] are used for visualizations and schematics. Further simulations of r(CAG)$_{10}$ and d(CAG)$_{10}$ followed the same simulation procedure.

#### Minimal model simulations of AB systems:

To study the impact of chain stiffness on condensate properties, we conduct minimal model MD simulations. We design an AB-type model, where only A and B form associative interactions mimicking complementary base pairing. The bonded interactions are modeled by a harmonic potential, the excluded volume potential of same-type interactions (A–A and B–B) are modeled via the Weeks–Chandler–Andersen (WCA) potential, and the attractive interaction is modeled by Lennard-Jones (LJ) potential. We simulate identical chains with the primary sequence of (AB)$_{25}$ and systems are composed of 125 chains. The bending stiffness is modeled by the typical cos potential, and we only vary the relative stiffness in our simulations to evaluate the effect on condensate properties. In the MD simulations, we use Lennard-Jones units and integration of equations of motion is carried out in the LAMMPS [14] package (23 June 2022 Version). Each system is simulated in the *NVT*-ensemble, employing a Langevin thermostat. The masses of the monomers are set to $m$=1, the monomer size and equilibrium bond length are set to $\sigma$=1, and the LJ potential minimum is set to $\varepsilon$=1. The excluded volume strength is also set to $\varepsilon$, and the bonded interaction is set to 10$\varepsilon$, and the bending stiffness $\kappa$ is varied from 2$\varepsilon$ to 12$\varepsilon$. Finally, the simulation temperature is set to $kT/\varepsilon$ = 0.8, with damping constant 0.5$\tau$. The simulations are performed for at least for 60,000,000 steps with the step size of 0.005$\tau$ to ensure proper equilibration. 1000 snapshots from the last 10,000,000 steps, which are sampled every 10,000 steps, are taken for analysis. For the droplet shape and neighbor analysis Ovito [15] is used. For the local order analysis, Q-tensor analysis is performed using custom codes.

#### Nucleic acid sample preparation for temperature-controlled microscopy:

Nucleic acids are obtained from Integrated DNA Technologies using salt-free purification and are stored in RNase free water (Ambion) at stock concentrations at either 200 µM (RNA) or 1 mM (ssDNA). RNA homopolymers were obtained from Sigma-Aldrich [poly(rC), Sigma #P4903-

25MG, Batch # 0000091053; poly(rU), Sigma #P9528-10MG, Lot #089M4052V] and reconstituted in RNase free water. 2'-O-methylated RNA were obtained from Gene Link and stored in RNase free water. Nucleic acids were prepared in buffer as described in the text by diluting RNA into RNase free water mixed with a 4x buffer stock to a final working concentration described in the main text. DNA samples were observed to have some aggregates in the stock, which were removed by spinning at 10,000 rpm and retaining the supernatant. The concentration of DNA in the supernatant was recorded.

Nucleic acid samples for phase separation assays were prepared as described previously [16,17]. In brief, glass slides were cut to fit the temperature stage to be approximately 1"x 0.8" using a carbide glass scribe (Thorlabs). Slides were cleaned using ethanol and kimwipes and then two strips of double-sided tape (Scotch) are placed approximately 1 mm apart with 18 mm square #1.5 coverslip placed on top. These channels can contain approximately 5 µL of sample. A small drop of 518N oil is placed at either end of the channel to prevent evaporation after filling. Samples were imaged immediately after preparation.

For samples that were tested with variation of pH, buffers containing equimolar concentrations of HEPES, MES, and sodium acetate were prepared at evenly spaced intervals of one pH unit by titrating with HCl. This buffer has previously been reported to have a linear behavior in the range of pH 2-8 [18]. Samples were then subjected to measurements using temperature-controlled microscopy.

**Temperature-dependent microscopy of RNA and DNA:**
Samples were imaged via temperature-controlled microscopy as previously described [16,17]. In brief, an Instec temperature-controlled stage was used to sweep temperatures between 0°C to 90°C. Temperature was set manually using the ramp setting which heats at ~5°C/min. Near $T_{phase}$ of a sample, the temperature was increased in 2°C increments every 5 minutes. The sample was cycled around $T_{phase}$ at least once beyond the initial measurement to confirm (or a lack thereof) reversibility. Temperature values were extracted using a K-type thermocouple in combination with a USB digital to analog converter. This outputs temperatures with time stamps that are aligned with each frame time using image metadata through custom python code. A Zeiss primovert microscope was used with a 40x air objective and a Blackfly S camera (Teledyne FLIR) controlled via Micromanager to acquire images at 1 second intervals. $T_{phase}$ values were extracted from three technical replicates.

**Estimation of the percolation temperature ($T_{prc}$) via shape relaxation:**
Videos of samples with percolation coupled to phase separation were identified and the temperatures of the start and end of shape relaxation were extracted manually through time-lapse image analysis. Here we define shape relaxation to be a change in particle size and/or morphology starting from an irregular cluster and ending in an energetically minimized circular shape. The width of these clusters was determined using Fiji to draw a line to reference the difference in size between two frames. All condensates of similar size relax at similar temperatures in a single image. A minimum of three trials were averaged to determine the start and end temperatures for shape relaxation and $T_{prc}$ was assigned to be the average of these temperatures.

**Small-angle X-ray scattering experiments and analysis:**
Small-angle x-ray scattering experiments were conducted using the BioSAXS facilities at the Cornell High Energy Synchrotron Source (CHESS). SAXS experiments were performed following the protocols described in the literature and the guidance of the staff overseeing the BioSAXS station at CHESS with a full description as follows. These experiments entailed preparation of

RNA and ssDNA in equivalent buffer conditions with varying concentrations of magnesium ions. These conditions were chosen to be close to but not within the two-phase regime for $r(CAG)_{31}$. Concentration of RNA was chosen to have sufficient contrast during the experiment.

Samples were prepared by spinning concentrated stocks of nucleic acids for 10 minutes before dilution in buffer containing 50 mM HEPES, pH 7.5 at room temperature, and either 1 mM, 5 mM, or 10 mM $MgCl_2$. Due to the sensitive nature of SAXS measurements and the fact that it is a contrast technique, samples were dialyzed against a master stock of buffer at the working concentration of buffer and ions using dialysis cups with an 8.5 kDa molecular weight cutoff. Samples were recovered from the dialysis cup with approximately 80% yield after overnight dialysis (approximately 18 hrs) with gentle agitation at 20°C. The master buffer stock and samples were kept on dry ice and transported to CHESS (5 hours transit time). Samples were thawed and inserted into a 96-well plate, which was subsequently sealed with foil seals (Agilent) and spun for two minutes at low speed to remove air bubbles. The 96-well plate was then placed on the pedestal inside the hutch where robotic handling transferred the solution to the capillary where it was imaged.

Data at the BioSAXS station at CHESS is collected and subsequently visualized using the BioXTAS RAW software during acquisition [19]. In order to have absolute scaling, scattering images of the empty capillary tube and water-filled capillary tube were collected. These were subsequently utilized in the RAW software in the advanced options to correct the data using reference standards for glassy carbon and water incorporated into the software. Units for I(q) after correction are $cm^{-1}$. During acquisition the sample was dithered and 100 images were captured at 1s exposures without any meaningful changes in signal that would indicate aggregation. Additionally, images of the empty and water filled capillary were collected for each temperature point and the correction was applied based on the temperature. Temperature of the capillary was verified independently using a thermocouple. Temperature of the capillary chamber was equilibrated for at least 15 minutes prior to collecting data after changing the temperature of the water bath.

Prior to collecting data for experimental samples at a given temperature, a set of images was collected for the buffer without the nucleic acids. The data was averaged and then used for subtraction from subsequent data points. For each experimental data point, these data were averaged and the subtracted curve was calculated small after a DC-offset chosen to place the high-q data slightly above the buffer [20]. This offset was only performed, if necessary, due to factors such as evaporation. We note that this was primarily mitigated by dialysis beforehand. The subtracted data was verified to be linear in the low-q regime, implying mono-dispersity. No sample with significant bending in the Guinier plot was used to estimate the radius of gyration as this would imply aggregation of the polymers [21]. A Guinier fit was applied to estimate the radius of gyration using the freely available RAW software for SAXS analysis. For those data which had upturns at low q Guinier fitting was constrained at the most to have $q_{min}*R_g \sim 0.65$ where the residuals for the fit were found to be randomly distributed.

The Guinier approximation [19,22] for a spherical scattering object is represented by the equation below, which was fit by using the RAW software.

$$I(q) \approx I(0)e^{\frac{-q^2 {R_g}^2}{3}}$$

We have included further details about the SAXS experiments in Supplementary Note 1.

**Circular dichroism (CD) spectroscopy on G-quadruplex forming nucleic acids:**
Nucleic acids were prepared following a protocol from the literature [23]. The working concentration was 10 μM in a buffer containing 20% w/v PEG (8kDa), 20 mM HEPES, pH 7.5 at room temperature, and either 150 mM of LiCl,150 mM of KCl or 5 mM $MgCl_2$. Samples were transferred into a 1mm quartz cuvette (Hellma). The cuvette was placed inside the CD instrument (Jasco J815 CD spectrometer). Circular dichroism measurements were taken at 20°C. Data was acquired over a range of 200 nm - 340 nm at 0.1 nm data pitch with a band width of 1 nm. The scanning mode was set to continuous, and the speed was 50 nm/min. Data was saved as the result of four accumulations. Data was exported as a text file and background was subtracted and the result was plotted using a custom python scripts.

**Thioflavin T (ThT) fluorescence measurements of G-quadruplex forming nucleic acids:**
Sample conditions were chosen where stable condensates could be observed at room temperature after heating and cooling except for the $(UUAGUG)_{10}$ which forms condensates at room temperature without heating. $r(CAG)_{31}$ and $(UUAGUG)_{10}$ were chosen as non-G-Quadruplex forming negative controls. We note that ThT also functions as a viscosity sensor and may not only report on G-quadruplex structure making it necessary to complement this data with other verifications of G-quadruplex structure such as through CD spectroscopy. This has been observed in the literature where non-G-quadruplex forming sequences may bind ThT but demonstrate much less enhancement of ThT signal [24,25]. Samples were prepared as described above and sealed in a custom chamber using oil and double-sided tape. Samples were subsequently placed with the cover-glass facing down on a heat block set to 95°C for approximately five minutes to induce condensate formation. Chambers were removed from heat and samples were imaged at room temperature (~ 24°C). Samples were mounted on an Andor Dragonfly spinning disc confocal microscope and imaged using the CF40 spinning disc for fluorescence images and the widefield mode for brightfield images. Z-stack acquisitions were performed using the Andor iXon 880 camera at 100 ms exposure with 30 gain for confocal and 10 gain for brightfield images. Figures of these images were prepared using Fiji and Adobe Illustrator.

**Data analysis and preparation:**
All experimental data was plotted using custom python scripts relying on the Numpy, Scipy, and Matplotlib libraries including the data processed using RAW except the Guinier fits which were exported from RAW. Figures in the main text were prepared using Adobe Illustrator and for chemical structures, ChemDraw, was used.

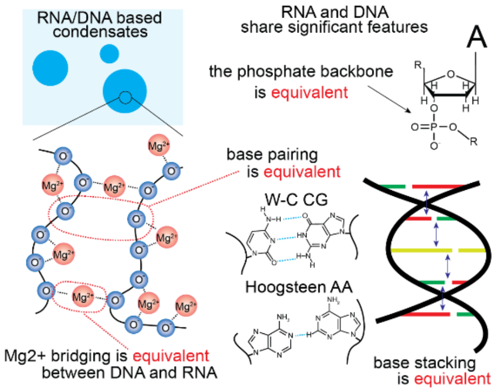


**Fig. S1. Cartoon representations of the different modes of interactions of single stranded nucleic acids.** While often described through their differences, many features of DNA and RNA are shared. The means of crosslinking are identical between ssDNA and RNA. The sole difference is the 2'-OH of ribose.

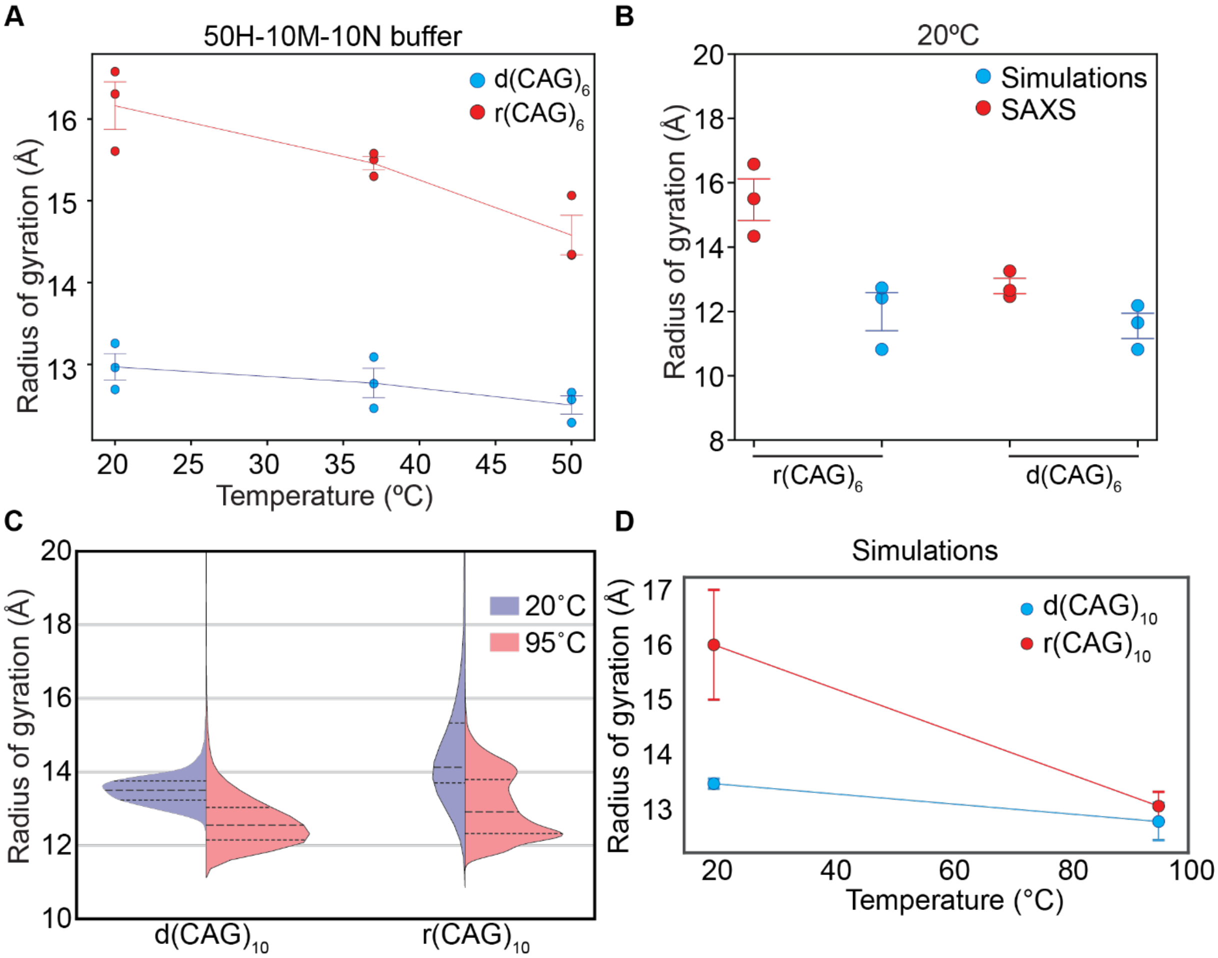


**Fig. S2. Radius of gyration ($R_g$) estimation of short $(CAG)_6$ RNA and ssDNA using Small Angle x-ray Scattering (SAXS). (A)** Comparison of the temperature dependence of $R_g$ estimated through Guinier fitting of SAXS data for 50 μM $(CAG)_6$ RNA (red) with ssDNA (blue) in a buffer containing 50 mM HEPES, pH 7.5 at RT, with 10 mM $MgCl_2$ and 10 mM NaCl. **(B)** Comparison of the $R_g$ for $(CAG)_6$ derived from SAXS data (red) and atomistic simulations (blue) for RNA and ssDNA. We find that without any calibration, the $R_g$ values at 20ºC from SAXS are on average 13.57 Å for ssDNA versus 11.26 Å from simulations, while for RNA, the values are 15.46 Å for SAXS and 11.70 Å for simulations. **(C)** A violin plot of the distribution of the radius of gyration ($R_g$) determined via all-atom molecular dynamics (MD) simulations for r$(CAG)_{10}$ at two different temperatures, 20°C (blue) and 95°C (red) in a box containing the equivalent of 150 mM $MgCl_2$. The mean and upper and lower quartiles are shown as dashed lines in the distributions. **(D)** The mean radius of gyration from three replicates of the simulations shown in **C** are plotted at two temperatures for DNA (blue) and RNA (red).”Buffer notation used: the number in front of “H” indicates the [HEPES] in mM, the number in front of “M” indicates the [$MgCl_2$] in mM, and the number in front of “N” indicates the [NaCl] in mM for each buffer. Error bars represent s.e.m. for n = 3 replicates.

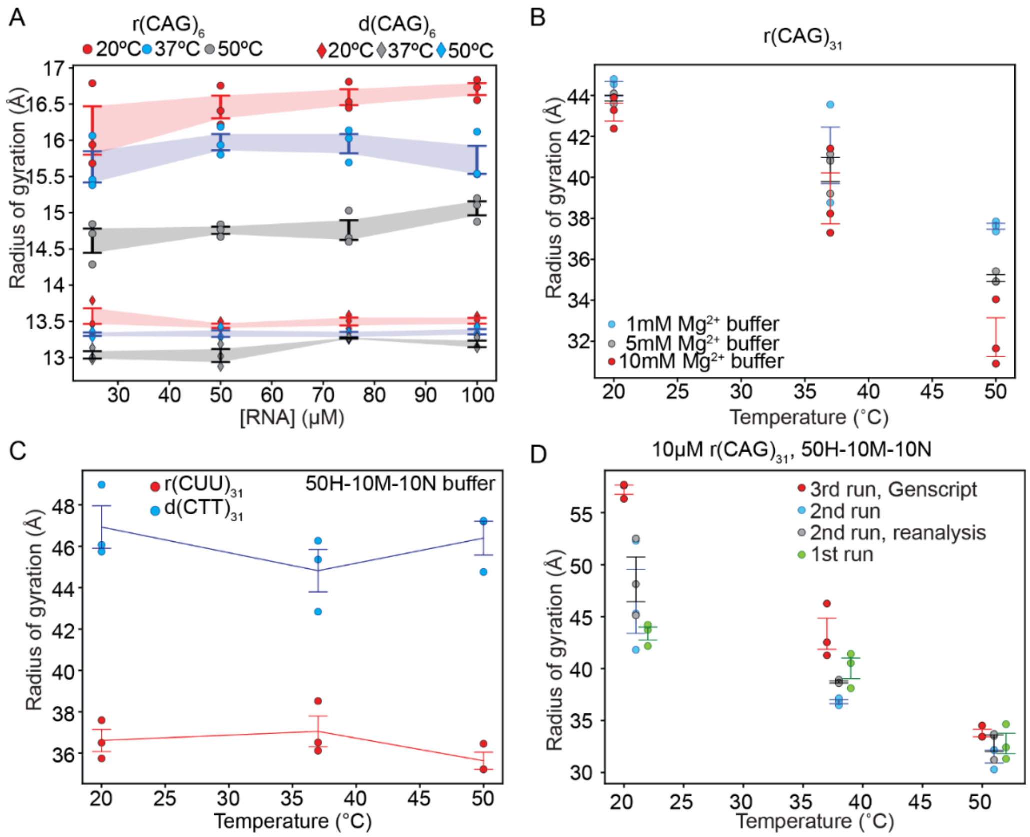


**Fig. S3. Controls for SAXS experiments. (A)** Radius of gyration ($R_g$) derived from SAXS data collected at different concentrations of $(CAG)_6$ RNA and ssDNA**.** Comparison of the $R_g$ at four concentrations of $(CAG)_6$ RNA (circles) to ssDNA (diamonds) in a buffer containing 50 mM HEPES, pH 7.5 at RT, with [$MgCl_2$] = 10 mM at three temperatures T = 20ºC (red), 37ºC (blue) or 50ºC (black). **(B)** Magnesium ion dependent coil-to-globule transition of $(CAG)_{31}$ RNA. Comparison of the $R_g$ of 10 μM r$(CAG)_{31}$ in a buffer containing 50 mM HEPES, pH 7.5 at RT, with [$MgCl_2$] = 1 mM (blue), 5 mM (black), or 10 mM (red). **(C)** Radius of gyration ($R_g$) invariance of U/T rich $(CUU/CTT)_6$ RNA/ssDNA. Shown here are temperature dependence of SAXS-derived $R_g$ values for 10 μM $(CUU)_{31}$ RNA or $(CTT)_{31}$ ssDNA at three temperatures in a buffer containing 50 mM HEPES, pH 7.5 at RT, with 10 mM $MgCl_2$ and 10 mM NaCl. **(D)** Radius of gyration ($R_g$) derived from SAXS data of $(CAG)_{31}$ RNA and ssDNA comparing the values from three independent SAXS experiments using RNA from three separate syntheses from two different vendors (two from IDT and one from Genscript) on three separate dates. Also, shown are two separate analyses of the SAXS data from the same run. Buffer notation used: the number in front of "H" indicates the [HEPES] in mM, the number in front of "M" indicates the [$MgCl_2$] in mM, and the number in front of "N" indicates the [NaCl] in mM for each buffer. Error bars represent s.e.m. for n = 3 replicates.

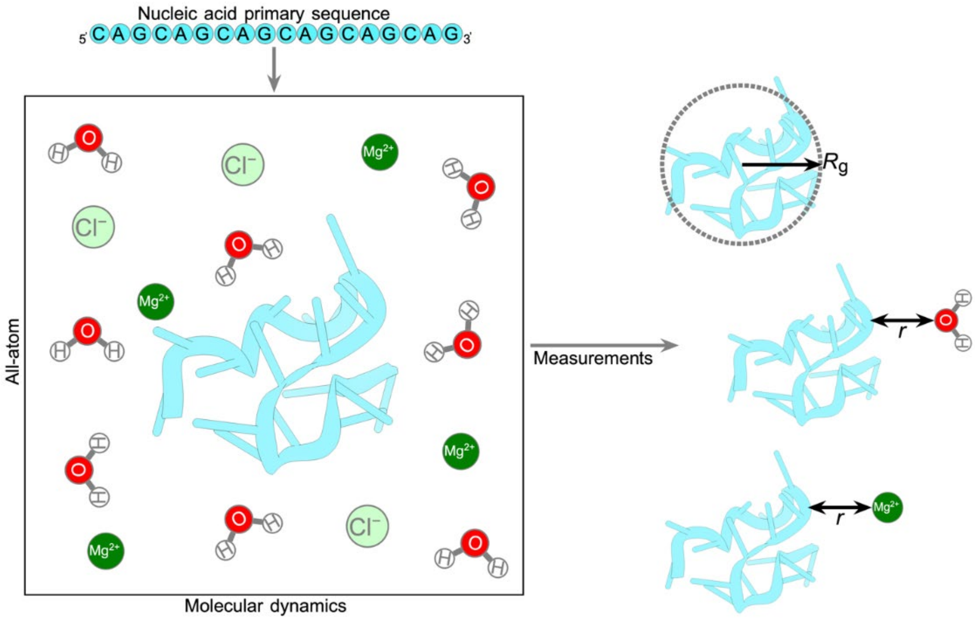


**Fig. S4. Schematics of atomistic simulations.** A schematic depicting the configuration of molecules in all-atom simulations is shown. The corresponding parameters radius of gyration, $R_g$, and the distance between phosphate oxygen (Op) in the backbone and the water or ions, $r$, are also depicted.

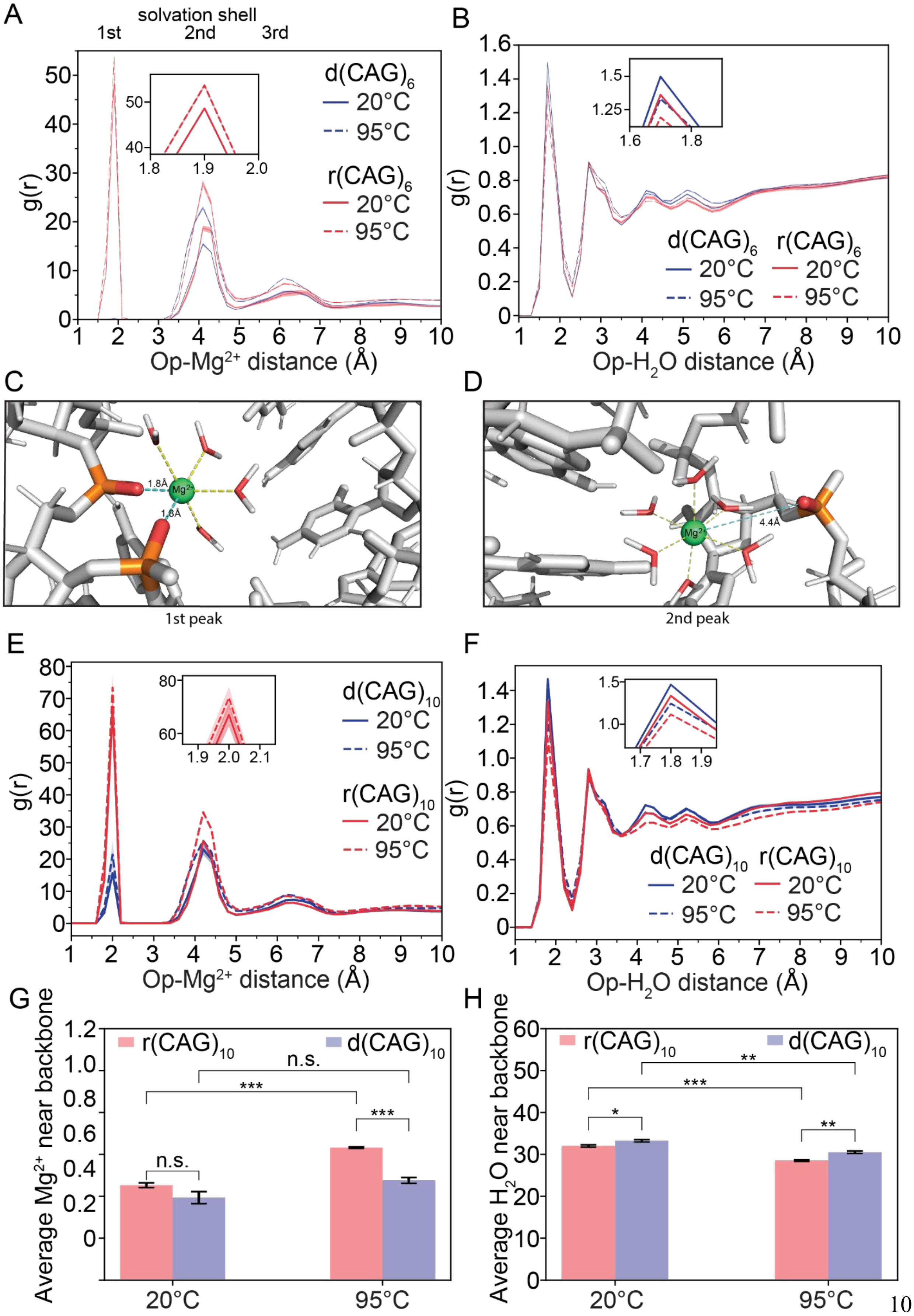
A
solvation shell
1st
2nd
3rd
d(CAG)6
20°C
95°C
r(CAG)6
20°C
95°C
g(r)
Op-Mg2+ distance (Å)
B
d(CAG)6
r(CAG)6
Op-H2O distance (Å)
C
1.8Å
Mg2+
1st peak
D
4.4Å
2nd peak
E
d(CAG)10
r(CAG)10
F
G
r(CAG)10
d(CAG)10
n.s.
***
Average Mg2+ near backbone
20°C
95°C
H
Average H2O near backbone
*
**

**Fig. S5. Radial distribution and representative visualization of the distance between Op and $Mg^{2+}$ or $H_2O$ in simulations with 150 mM $MgCl_2$. (A)** The radial distribution [g(r)] of distances between the phosphate oxygen (Op) and $Mg^{2+}$ ions in simulations at two temperatures, 20°C (solid) and 95°C (dashed), for $r(CAG)_6$ (red) and $d(CAG)_6$ (blue). **(B)** The distribution of radial distances between phosphate oxygen (Op) and water molecules in simulations at two temperatures, 20°C (solid) and 95°C (dashed), for $r(CAG)_6$ (red) and $d(CAG)_6$ (blue). The errors are shown as error bands corresponding to three independent simulations. Red or blue shading represents the error bands for the data. **(C-D)** Relative proximity of $Mg^{2+}$ ions from phosphate oxygen (Op). The distance between the $Mg^{2+}$ ions and the waters (yellow dashes) and Op (blue dashes) is shown for $Mg^{2+}$ ions within the first solvation shell, **C,** and the second solvation shell, **D**. **(E)** The radial distribution [g(r)] of distances between the phosphate oxygen (Op) and $Mg^{2+}$ ions in simulations at two temperatures, 20°C (solid) and 95°C (dashed), for $r(CAG)_{\mathbf{10}}$ (red) and $d(CAG)_{\mathbf{10}}$ (blue). **(F)** The distribution of radial distances between phosphate oxygen (Op) and water molecules in simulations at two temperatures, 20°C (solid) and 95°C (dashed), for $r(CAG)_{\mathbf{10}}$ (red) and $d(CAG)_{\mathbf{10}}$ (blue). The errors are shown as error bands corresponding to three independent simulations. Red or blue shading represents the error bands for the data. **(G)** The average number of $Mg^{2+}$ ions located in the $1^{st}$ and $2^{nd}$ solvation shells, i.e. within 5 Å of the phosphate oxygen (Op) in simulations at two temperatures, 20°C and 95°C, for $r(CAG)_{10}$ (red bars) and $d(CAG)_{10}$ (blue bars). P-values from left to right are p1 = 0.12631, p2 = 0.00011, p3 = 0.06096, p4 = 0.00039. (H) The average number of $H_2O$ molecules located in the $1^{st}$ and $2^{nd}$ solvation shells, i.e. within 5 Å of the phosphate oxygen (Op) in simulations at two temperatures, 20°C and 95°C, for $r(CAG)_{10}$ (red bars) and $d(CAG)_{10}$ (blue bars). P-values from left to right are p1 = 0.03486, p2 = 0.00044, p3 = 0.00242, p4 = 0.00377.

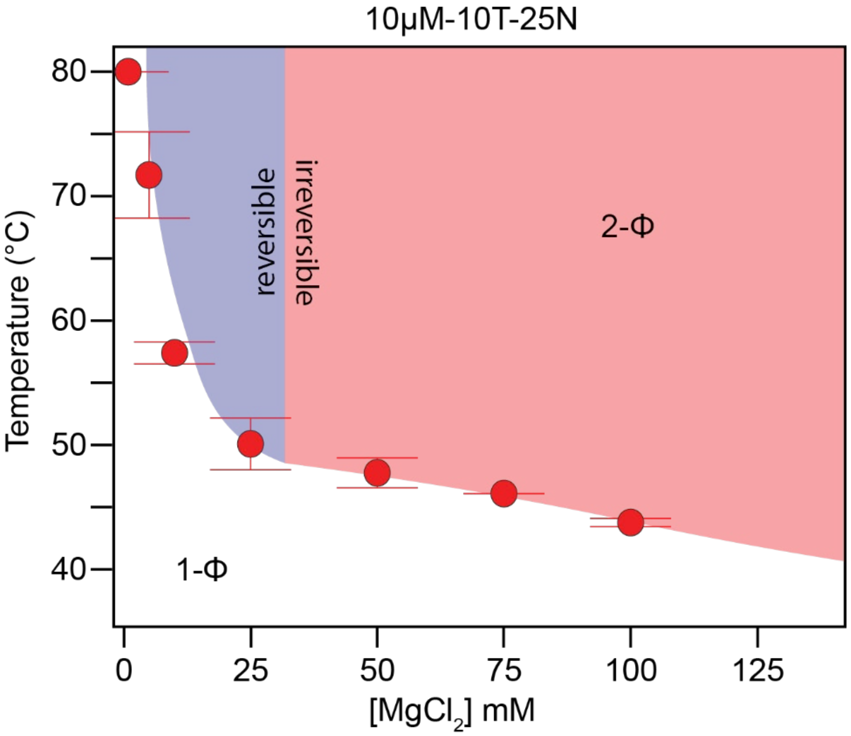


**Fig. S6. $Mg^{2+}$ ion dependence of percolation transition for $r(CAG)_{31}$ condensates.** Data replotted from the SI of Wadsworth, et al 2023 [17]. Shown here is a zoomed-in view of RNA state diagram from Figure 2C to highlight the window of reversibility. The shaded region indicates the 2-Φ regime for $r(CAG)_{31}$ while the color indicates the reversibility (blue) or irreversibility (red). Buffer notation used: the number in front of "T" indicates the [Tris-HCl] in mM, the number in front of "M" indicates the [$MgCl_2$] in mM, and the number in front of "N" indicates the [NaCl] in mM in the buffer. Error bars represent s.e.m. for n = 3 replicates.

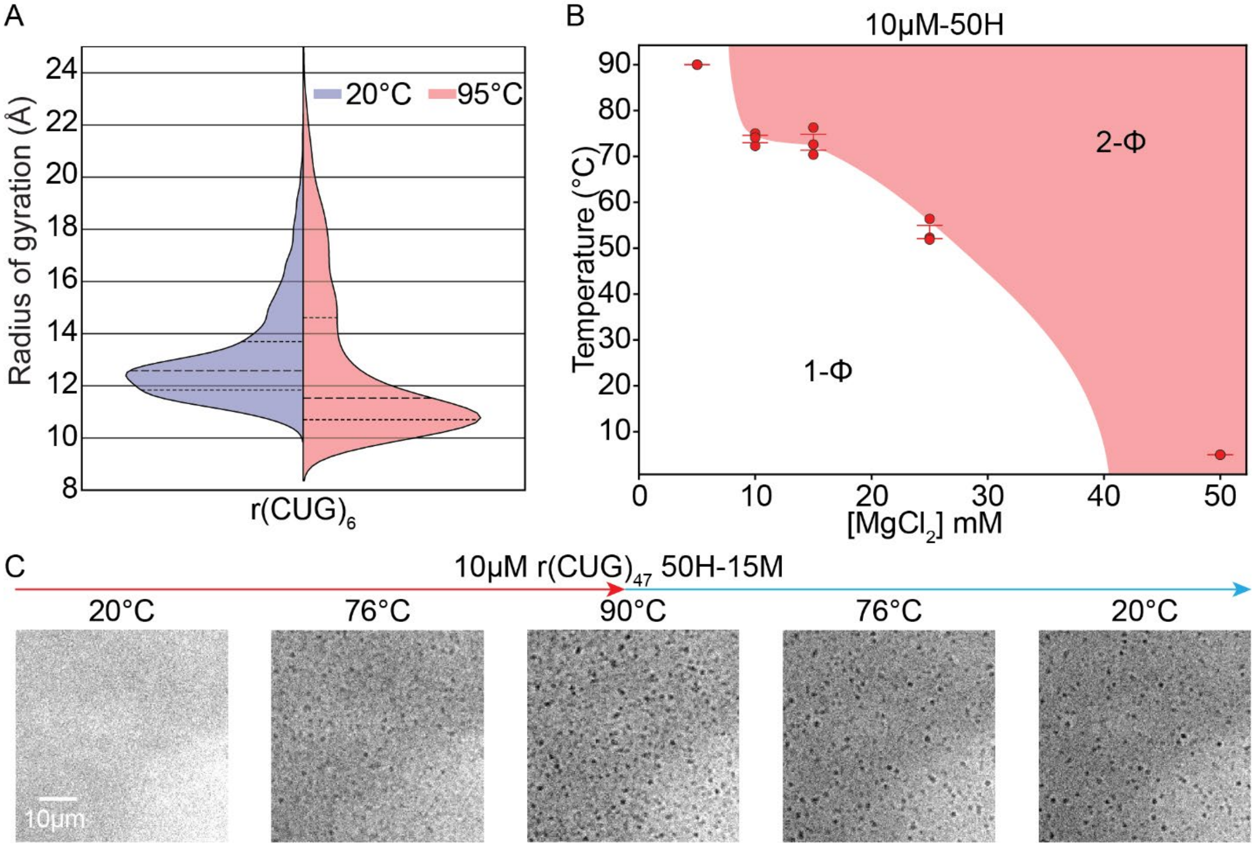


**Fig. S7. Temperature-dependent compaction and phase separation of $(CUG)_n$ RNA. (A)** A violin plot of the distribution of the radius of gyration ($R_g$) determined via all-atom molecular dynamics (MD) simulations for $r(CUG)_6$ at two different temperatures, 20°C (blue) and 95°C (red) in a box containing the equivalent of 150 mM $MgCl_2$. The mean and upper and lower quartiles are shown as dashed lines in the distributions. We determine that the persistence length $L_p$ for these two conditions is 6.07 ± 1.22 Å and 5.85 ±0.82 Å respectively. **(B)** A state diagram showing the [$MgCl_2$] dependence of $r(CUG)_{47}$ phase behavior in a buffer containing 10 μM $r(CUG)_{47}$ and 50 mM HEPES, pH 7.5 at RT. The color and shading indicate irreversibility (red) of the 2-Φ regime (shaded). Error bars represent s.e.m. for n = 3 replicates. **(C)** Representative images of thermoresponsive phase separation of 10 μM $r(CUG)_{47}$ in a buffer containing 50 mM HEPES, pH 7.5 at RT, and 15 mM $MgCl_2$. The LCPT of the sample is 66.3 ± 1.0°C. Buffer notation used: the number in front of "H" indicates the [HEPES] in mM and the number in front of "M" indicates the [$MgCl_2$] in mM in the buffer.

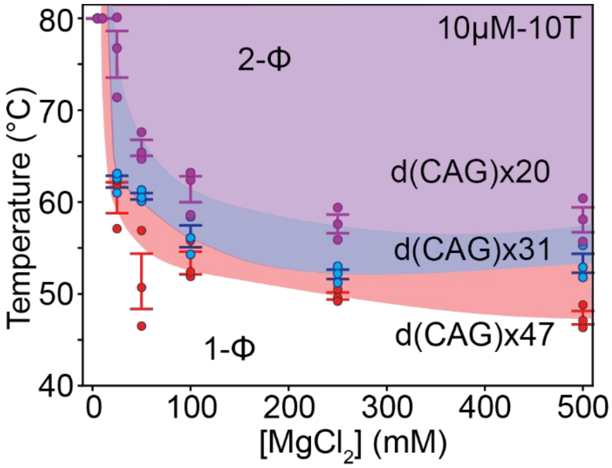


**Fig. S8. Length dependence of phase separation for d(CAG)$_n$**. A state diagram comparing the [$MgCl_2$] dependence of LCPTs for d(CAG)$_n$, where n = 20, 31, or 47. The shaded region indicates the 2-Φ regime for d(CAG)$_{47}$ (red), d(CAG)$_{31}$ (blue), and d(CAG)$_{20}$ (purple). We observe no percolation (CAG)$_n$ DNA. Buffer notation used: the number in front of "T" indicates the [Tris-HCl] in mM in the buffer. Error bars represent s.e.m. for n = 3 replicates.

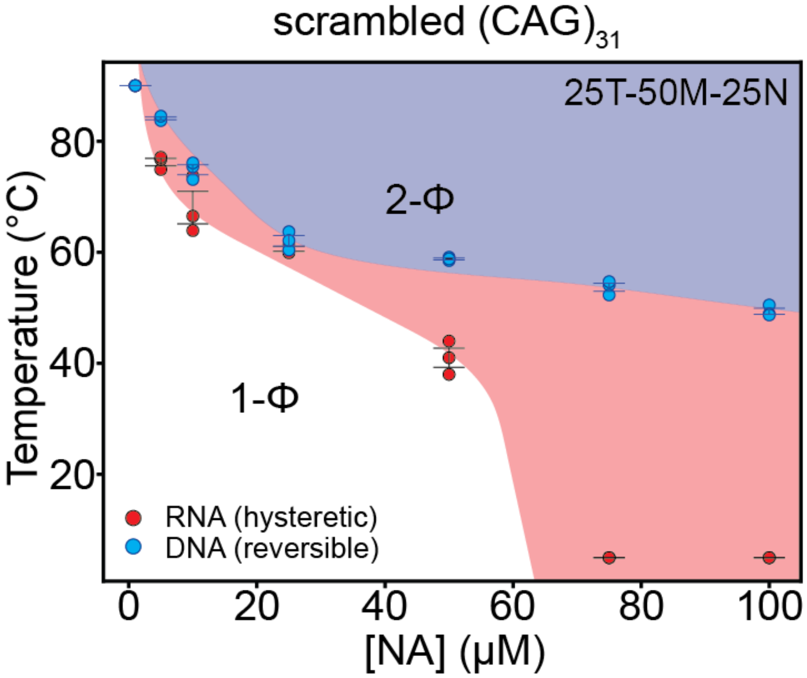


**Fig. S9. Comparison of the phase behavior of a scrambled $(CAG)_n$ equivalent RNA with the corresponding ssDNA.** Their respective sequences are shown in Supplementary table 1. A state diagram comparing the [NA] dependence of LCPTs for a scrambled sequence with equivalent amounts of C, A, and G to $(CAG)_{31}$, where the order was randomized (see SI Table 1). The shaded region indicates the 2-Φ regime for scrambled $r(CAG)_{31}$ (red), and scrambled $d(CAG)_{31}$ (blue). Buffer notation used: the number in front of "T" indicates the [Tris-HCl] in mM in the buffer. Error bars represent s.e.m. for n = 3 replicates.

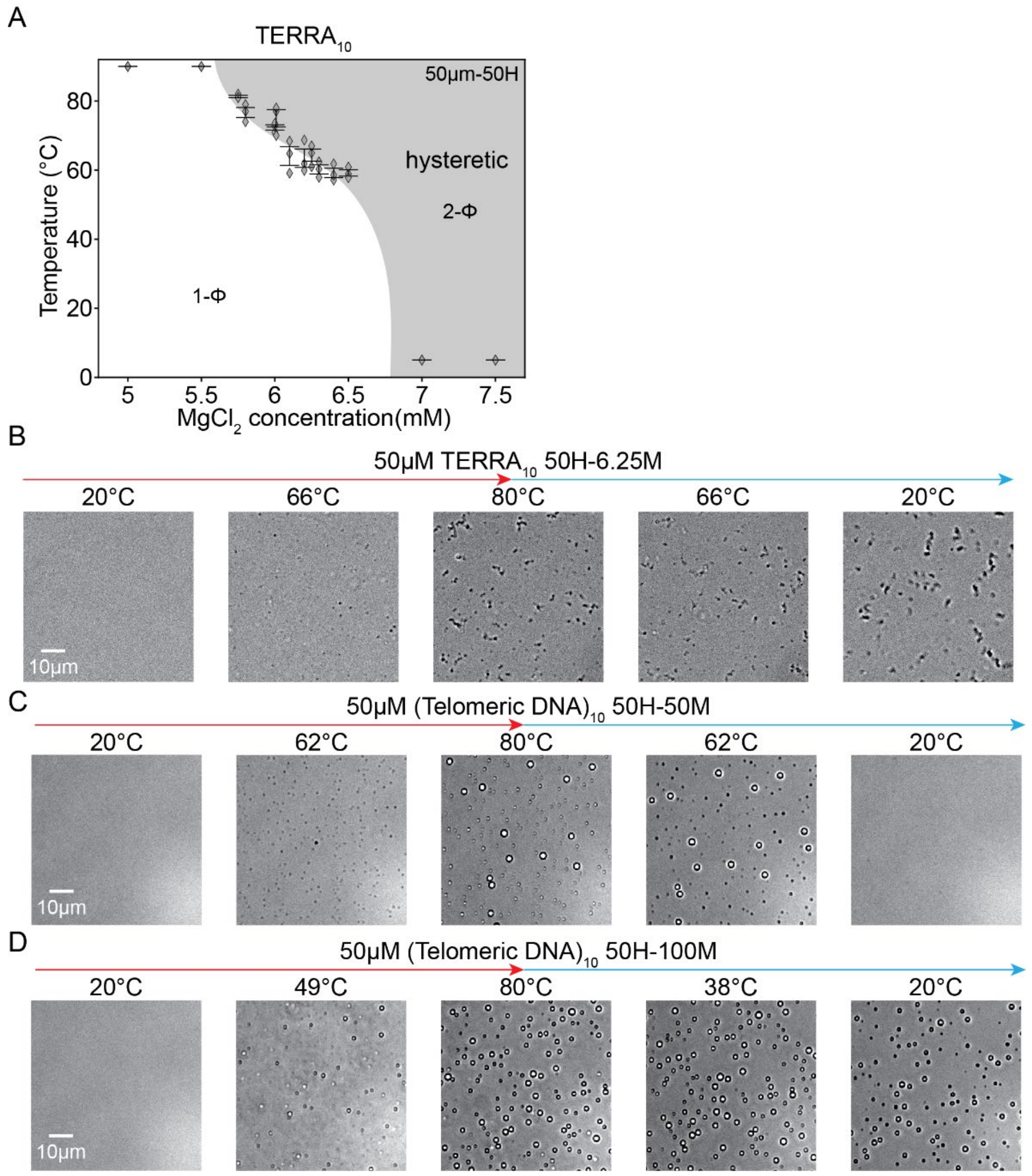


**Fig. S10. Magnesium dependence of phase separation and percolation of $(TERRA)_{10}$ and $(hTelo)_{10}$. (A)** A state diagram showing the $[MgCl_2]$ dependence of $TERRA_{10}$ phase behavior in a buffer containing 50 µM $(TERRA)_{10}$ and 50 mM HEPES, pH 7.5 at RT. The shading indicates the 2-Φ regime. Data is replotted from Mahendran et al. 2024 [16]. Error bars represent s.e.m. for n = 3 replicates. **(B)** Percolation of $(TERRA)_{10}$. Representative brightfield microscopy images of irreversible thermoresponsive phase separation of 50 µM $(TERRA)_{10}$ in a buffer containing 50 mM HEPES, pH 7.5 at RT, and 6 mM $MgCl_2$. The LCPT is 66.1 ± 1.8°C. **(C)** Representative brightfield microscopy images of thermoresponsive phase separation of 50 µM $(htelo)_{10}$ in a buffer containing 50 mM HEPES, pH 7.5 at RT, and 50 mM $MgCl_2$. The LCPT is 60.4 ± 0.48°C. **(D)** Representative brightfield microscopy images of thermoresponsive phase separation of 50 µM $(htelo)_{10}$ in a buffer containing 50 mM HEPES, pH 7.5 at RT, and 100 mM $MgCl_2$. The LCPT is 49.2 ± 0.78°C. Buffer notation used: the number in front of "H" indicates the [HEPES] in mM and the number in front of "M" indicates the $[MgCl_2]$ in mM in the buffer.

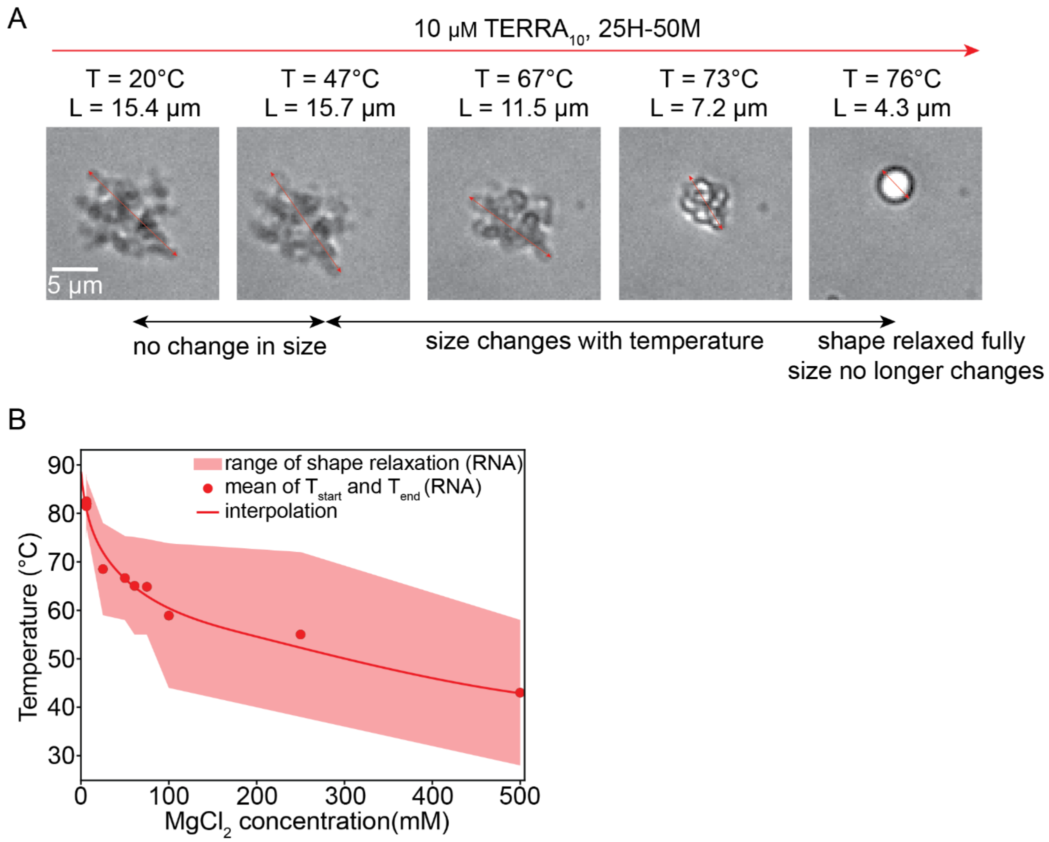


**Fig. S11. Shape relaxation of percolated $(TERRA)_{10}$ condensates. (A)** Representative images of shape relaxation of $(TERRA)_{10}$ condensates during a thermal ramp. $(TERRA)_{10}$ condensates were formed utilizing 10 μM $TERRA_{10}$ in a buffer containing 25 mM HEPES, pH 7.5 at RT, and 50 mM $MgCl_2$. **(B)** A plot showing the upper and lower bounds of shape relaxation (boundaries of red shading) with the mean (red data) representing the percolation temperature ($T_{prc}$). Buffer notation used: the number in front of "H" indicates the [HEPES] in mM and the number in front of "M" indicates the [$MgCl_2$] in mM in the buffer.

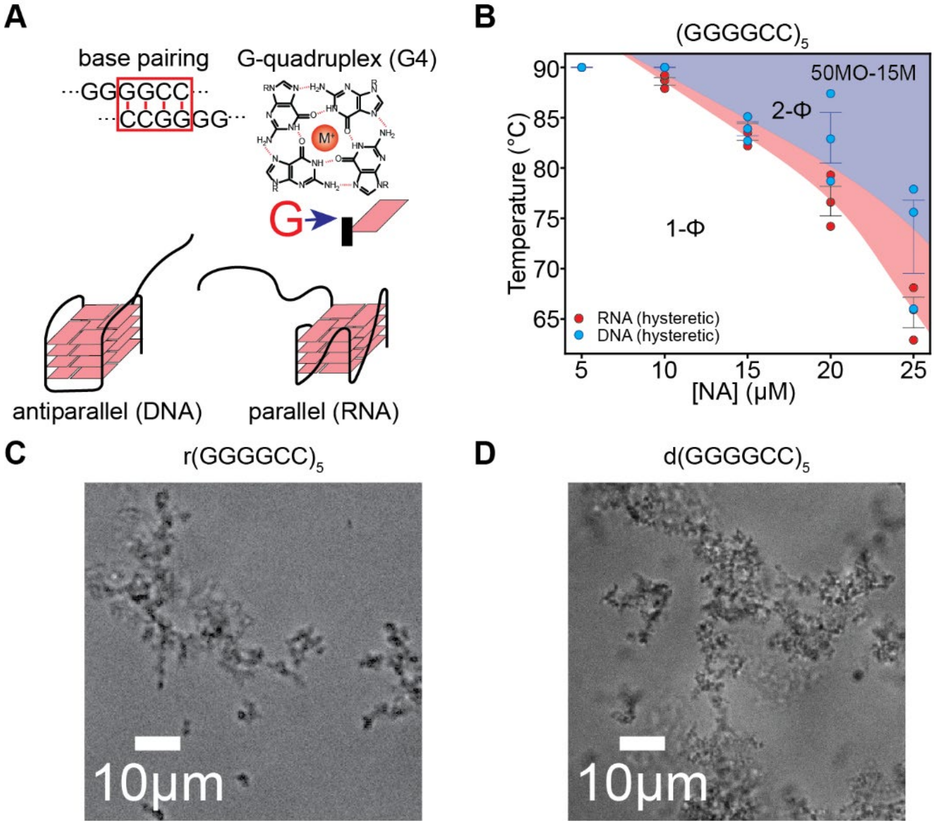


**Fig. S12. Comparison of the phase separation of G-quadruplex forming $(GGGGCC)_5$ RNA vs. ssDNA. (A).** A schematic showing the base pairing and possible G-quadruplex structure of $(GGGGCC)_5$ ssDNA or RNA. **(B)** A state diagram comparing the [NA] dependence of LCPTs in a buffer containing 50 mM MOPS, pH 7.5 at RT 15 mM $MgCl_2$.The shaded region indicates the 2-Φ regime for r$(GGGGCC)_5$ (red), and d$(GGGGCC)_5$ (blue). In both cases, the phase separation is irreversible. **(C)** Representative image of the fractal clusters formed at RT by 50 µM r$(GGGGCC)_5$ in a buffer containing 50 mM MOPS, pH 7.5 at RT, and 50 mM $MgCl_2$. **(D)** Representative image of the fractal clusters formed at RT by 50 µM d$(GGGGCC)_5$ in a buffer containing 50 mM MOPS, pH 7.5 at RT, and 50 mM $MgCl_2$. Clusters in **C** and **D** were not reversed. Buffer notation used: the number in front of "MO" indicates the [MOPS] in mM and the number in front of "M" indicates the [$MgCl_2$] in mM in the buffer. Error bars represent s.e.m. for n = 3 replicates.

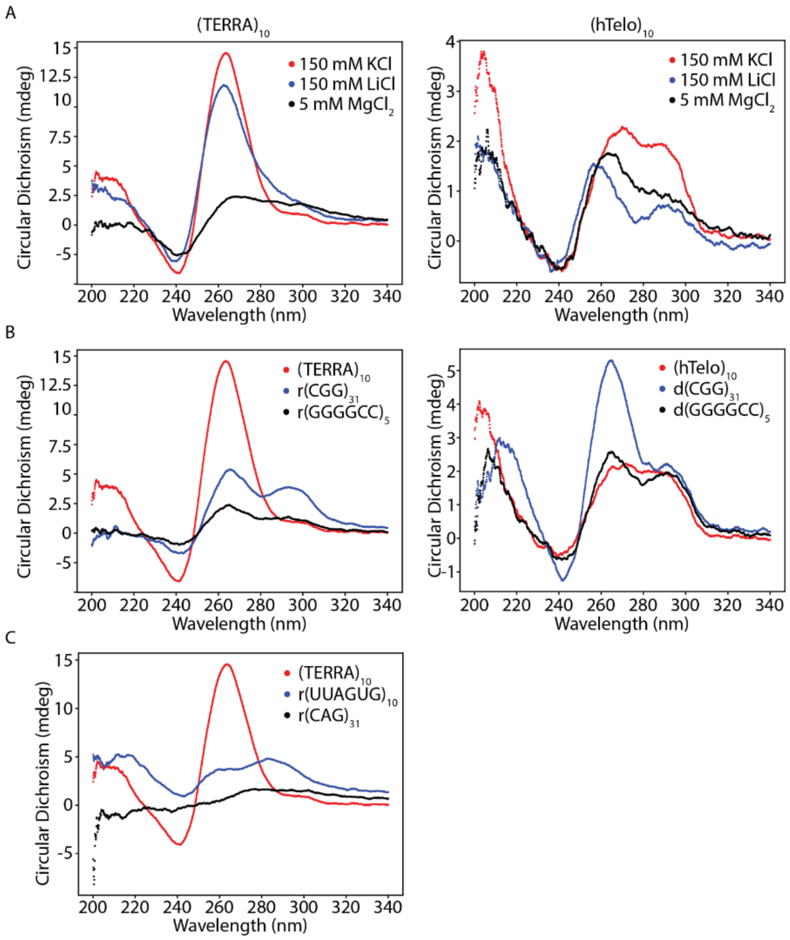


**Fig. S13. Circular dichroism (CD) measurements of putative G-quadruplex forming RNA and ssDNA**. **(A)** Representative circular dichroism spectra for 10 µM $(TERRA)_{10}$ (left) or $(hTelo)_{10}$ (right) in a buffer containing 50 mM Tris-HCl, pH 7.5, 20% PEG (8kDa) and either 150 mM KCl (red), 150 mM LiCl (blue) or 5 mM $MgCl_2$ (black). **(B)** A comparison of representative circular dichroism spectra 10 µM putative G-quadruplex forming RNA and DNA G-quadruplex forming RNA to $(TERRA)_{10}$ (left) or $(hTelo)_{10}$ (right) from (A) in the 150 mM KCl condition. **(C)** A comparison of 10 µM $r(CAG)_{31}$ and $(UUAGUG)_{10}$ which function as negative controls for G-quadruplex formation. We note that all curves except $r(CAG)_{31}$ can qualitatively fit into one of the three categories of G-quadruplex structures defined by a curated data set in the literature [26].

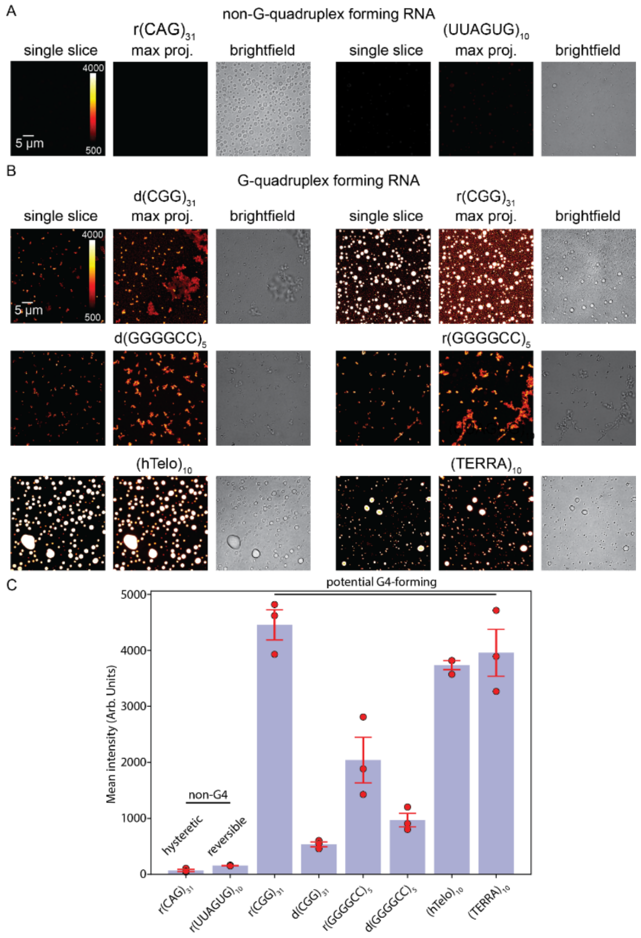


**Fig. S14. Thioflavin T (ThT) fluorescence of condensates formed by putative G-quadruplex forming RNA and ssDNA**. **(A)** ThT fluorescence images of condensates formed by two non-G-quadruplex forming RNA. r(CAG)$_{31}$ condensates at 100 µM RNA in a buffer containing 50 mM Tris-HCl, pH 7.5, and 50 mM $MgCl_2$ are shown for a single z-slice (left), a maximum intensity projection (middle) and a brightfield image (right) of the same field of view. We note that RNA condensates were generated by heating at 95ºC for 5 minutes and cooling to room temperature before imaging which is below the cloud point temperature for this condition indicating they are dynamically arrested. (Right). Images of condensates formed by a mutant of TERRA that is deficient in G-quadruplex formation. r(UUAGUG)$_{10}$ condensates at 50 µM RNA in a buffer containing 50 mM HEPES, pH 7.5, and 25 mM $MgCl_2$ are shown. **(B)** ThT fluorescence images of three pairs of putative G-Quadruplex ssDNA (left set) and RNA (right set). The top left set of images shows d(CGG)$_{31}$ condensates at 100 µM DNA in a buffer containing 50 mM Tris-HCl, pH 7.5, and 50 mM $MgCl_2$. The top right set shows r(CGG)$_{31}$ condensates at100 µM RNA in a buffer containing 50 mM Tris-HCl, pH 7.5, and 50 mM $MgCl_2$. The middle left shows d(GGGGCC)$_5$ condensates at 25 µM DNA in a buffer containing 50 mM MOPS, pH 7.5, and 15 mM $MgCl_2$. The middle right shows r(GGGGCC)$_5$ condensates at 25 µM RNA in a buffer containing 50 mM MOPS, pH 7.5, and 15 mM $MgCl_2$. The bottom left shows (hTelo)$_{10}$ condensates at 50 µM DNA in a buffer containing 50 mM Tris-HCl, pH 7.5, and 250 mM $MgCl_2$. The bottom right shows (TERRA)$_{10}$ condensates at 50 µM RNA in a buffer containing 50 mM Tris-HCl, pH 7.5, and 25 mM $MgCl_2$. Conditions were chosen for all cases in A and B to have condensates that exist at room temperature. **(C)** A bar chart showing the mean fluorescence intensity of condensates in A and B. Error bars represent s.e.m. for n = 3 replicates.

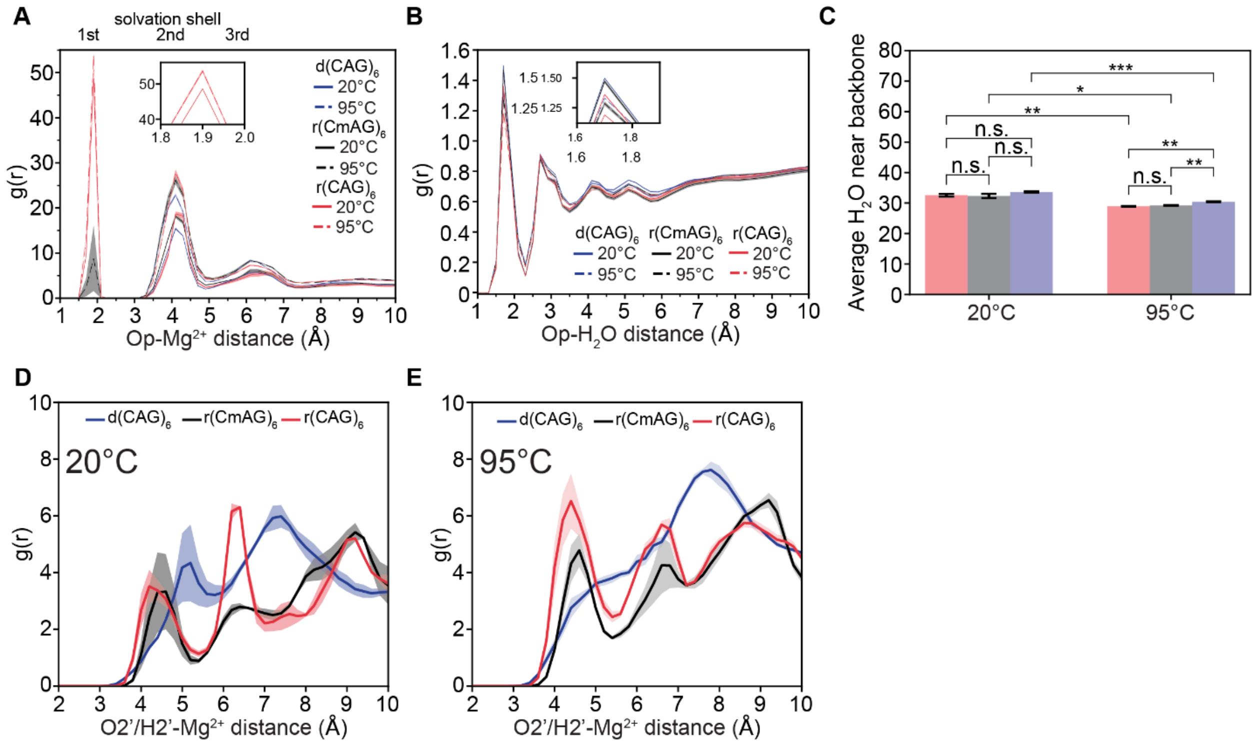


**Fig. S15. Radial distribution of the distance between phosphate oxygen (Op) and $H_2O$ for r(CmAG)$_6$ in simulations with 150 mM $MgCl_2$. (A)** The radial distribution [$g(r)$] of distances between phosphate (Op) and $Mg^{2+}$ ions in simulations at two temperatures, 20°C (solid) and 95°C (dashed), for r(CAG)$_6$ (red**),** r(CmAG)$_6$ (black), and d(CAG)$_6$ (blue). **(B)** The radial distribution [$g(r)$] of distances between phosphate (Op) and water molecules in simulations at two temperatures, 20°C (solid) and 95°C (dashed), for r(CAG)$_6$ (red**),** r(CmAG)$_6$ (black), and d(CAG)$_6$ (blue). **(C)** A bar plot representing the integrated number of water molecules in the first and second solvation shell. P-values are from right to left 0.27458, 0.27458, 1.0000, 0.81490, 1.0000, 0.56144, 0.53908, 0.27944, and 0.69176. **(D)** The radial distribution [$g(r)$] of distances between the 2'-group of RNA or DNA (O2'/H2') and $Mg^{2+}$ ions in simulations at 20°C for r(CAG)$_6$ (red**),** r(CmAG)$_6$ (black), and d(CAG)$_6$ (blue). **(E)** The radial distribution [$g(r)$] of distances between the 2'-group of RNA or DNA (O2'/H2') and $Mg^{2+}$ ions in simulations at 95°C for r(CAG)$_6$ (red), r(CmAG)$_6$ (black), and d(CAG)$_6$ (blue). The errors are shown as error bands corresponding to three independent simulations. Red, gray, or blue shading represents the error bands for the data in A, B, D, and E.

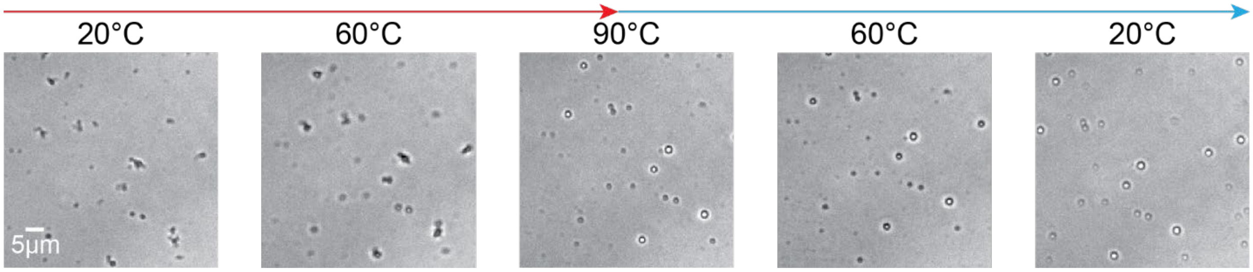


**Fig. S16. Percolation of r(CmAG)$_{20}$ condensates in the presence of high concentrations of calcium ions**. Shown is 10µM r(CmAG)$_{20}$ in a buffer containing 50 mM HEPES, pH 7.5 at RT, with 250 mM $CaCl_2$. The small clusters undergo shape relaxation as the temperature increases.

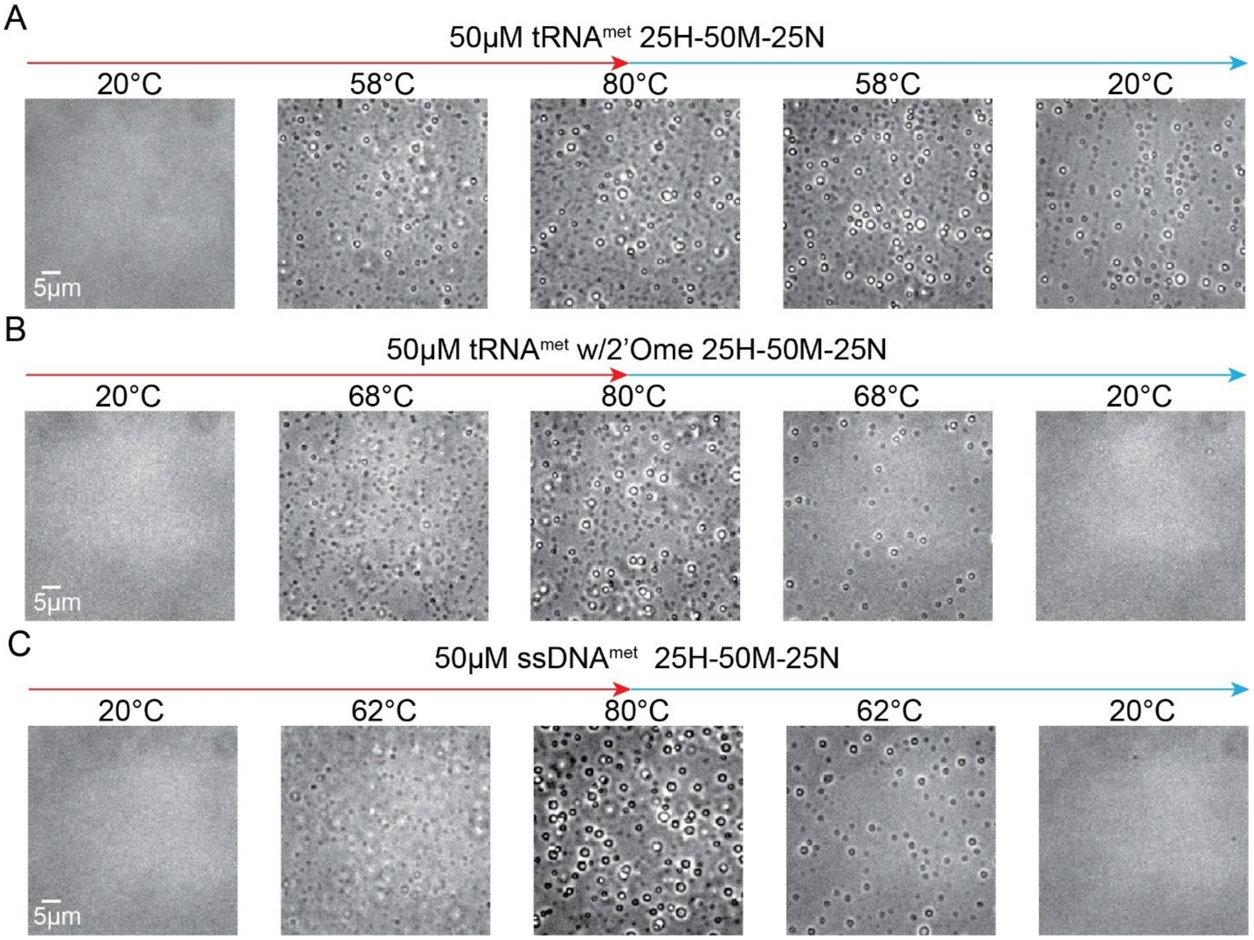


**Fig. S17. Condensation behavior of a tRNA$^{met}$ derived from *S. Acidocaldarius*. (A)** Shown here are condensates formed by 50 µM tRNA$^{met}$ in a buffer containing 25 mM HEPES, pH 7.5 at RT, with 50 mM $MgCl_2$ and 25 mM NaCl. Percolation is observed for this RNA at all concentrations of RNA. **(B)** Shows condensates formed by 50 µM 2'-Ome modified tRNA$^{met}$ in a buffer containing 50 mM HEPES, pH 7.5 at RT, with 50 mM $MgCl_2$ and 25 mM NaCl. Percolation is not observed. **(C)** Shows are condensates formed by 50 µM ssDNA$^{met}$ in a buffer containing 25 mM HEPES, pH 7.5 at RT, with 50 mM $MgCl_2$ and 25 mM NaCl. Percolation is not observed. Buffer notation used: the number in front of "H" indicates the [HEPES] in mM, the number in front of "M" indicates the [$MgCl_2$] in mM, and the number in front of "N" indicates the [NaCl] in mM for each buffer.

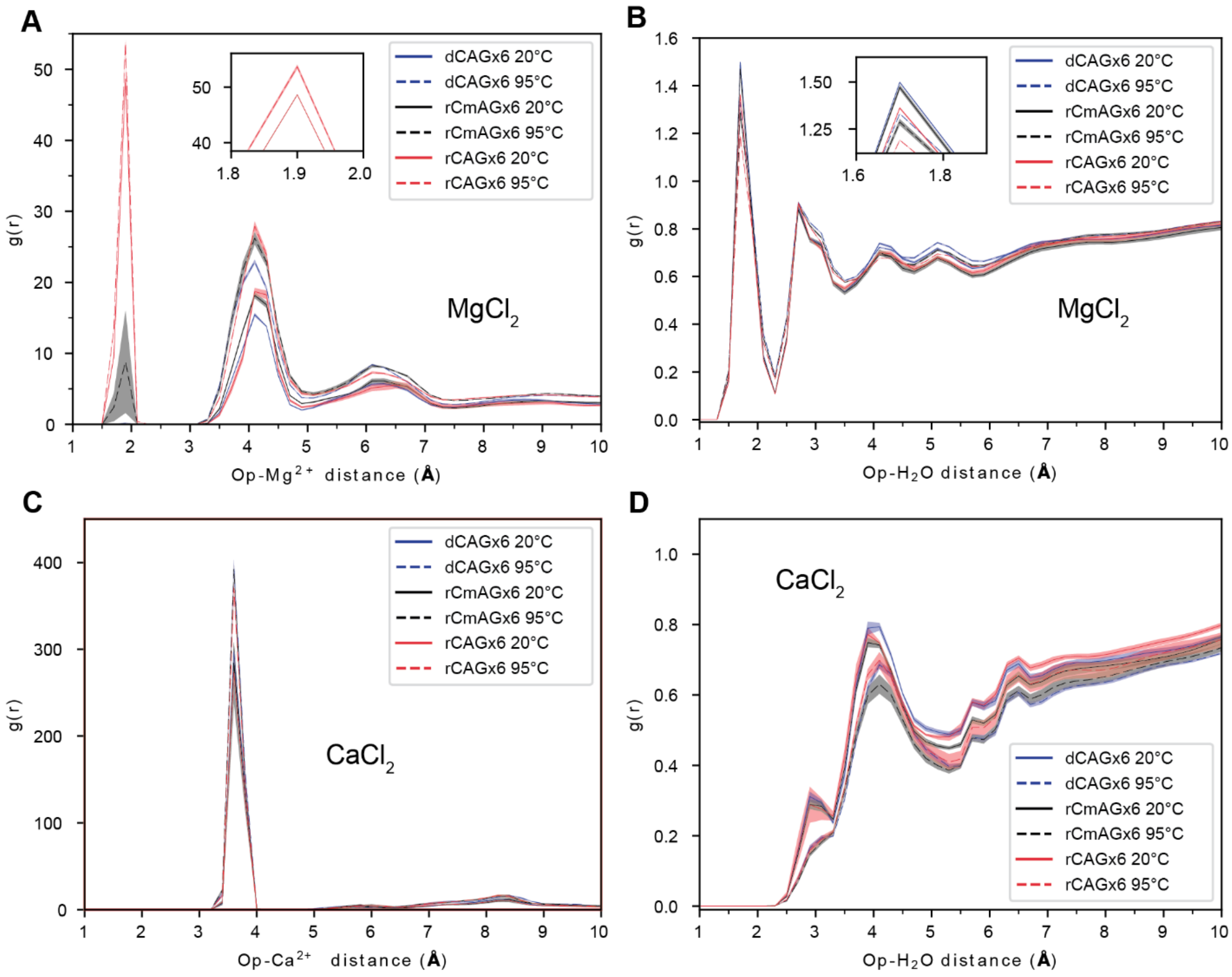


**Fig. S18. Radial distribution of different ion types ($Mg^{2+}$/$Ca^{2+}$) relative to the backbone of nucleic acids (NAs), from atomistic simulations.** The ion distribution (**A, C**) around the phosphate backbone and the water distribution (**B, D**) around the phosphate backbone. Whereas water molecules are more ordered around the backbone in the presence of $Mg^{2+}$, waters are more excluded from the backbone when $Ca^{2+}$ is present. The errors are shown as error bands corresponding to three independent simulations.

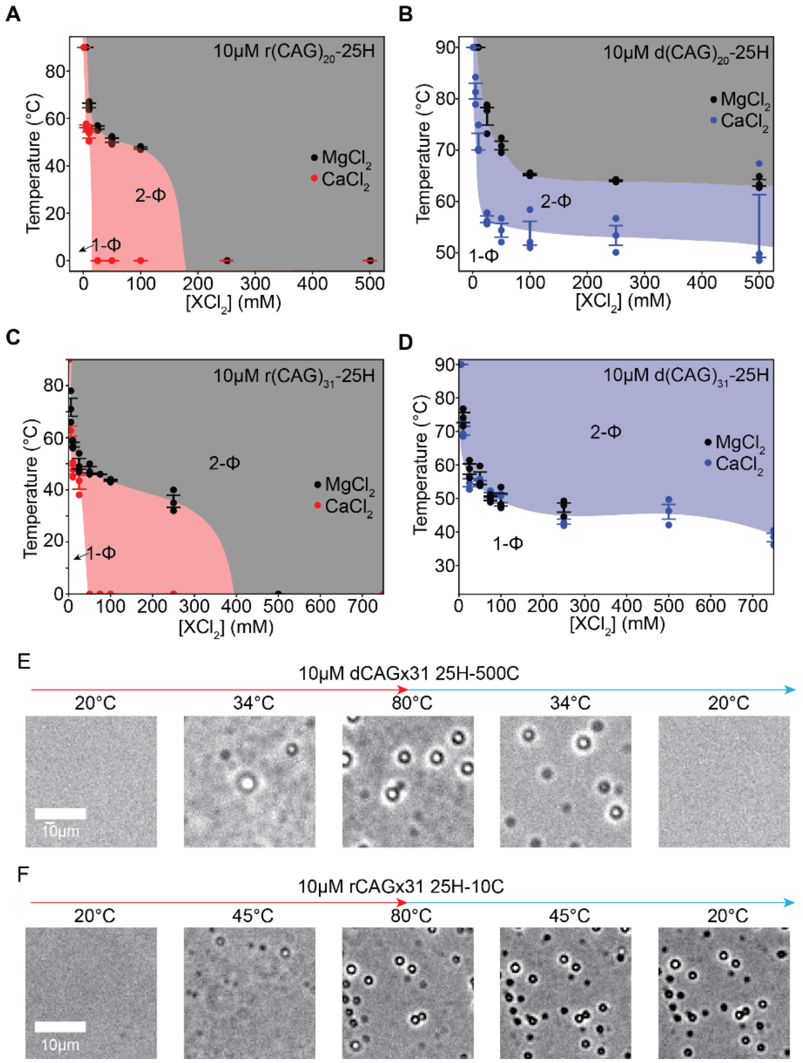


**Fig. S19. Comparison of the differential effects of calcium and magnesium on phase separation of (CAG)$_{20}$ and (CAG)$_{31}$ RNA and ssDNA**. (A) State diagram comparing [$MgCl_2$] (black) to [$CaCl_2$] (red) dependence for r(CAG)$_{20}$. Shading represents the 2-Φ regime for [$MgCl_2$] (grey) to [$CaCl_2$] (red). (B) State diagram comparing [$MgCl_2$] (black) to [$CaCl_2$] (red) dependence for d(CAG)$_{20}$. Shading represents the 2-Φ regime for [$MgCl_2$] (grey) to [$CaCl_2$] (blue). (C) State diagram comparing [$MgCl_2$] (black) to [$CaCl_2$] (red) dependence for r(CAG)$_{31}$. Shading represents the 2-Φ regime for [$MgCl_2$] (grey) to [$CaCl_2$] (red). (D) State diagram comparing [$MgCl_2$] (black) to [$CaCl_2$] (blue) dependence for d(CAG)$_{31}$. Shading represents the 2-Φ regime for [$MgCl_2$] (grey) to [$CaCl_2$] (blue). In all diagrams, red/grey is used to indicate irreversible phase separation while blue is used to indicate reversible phase separation. (E) Representative images of thermoresponsive phase separation of 10 μM d(CAG)$_{31}$ in a buffer containing 25 mM HEPES, pH 7.5 at RT, and 500 mM $CaCl_2$. The LCPT is 46.3 ± 2.2°C. (F) Representative images of thermoresponsive phase separation of 10 μM r(CAG)31in a buffer containing 25 mM HEPES, pH 7.5 at RT, and 10 mM $MgCl_2$. The LCPT is 47.7 ± 1.6°C. We note that RNA percolates at relatively lower [$Ca^{+2}$] while the DNA does not percolate at all conditions tested. Buffer notation used: the number in front of "H" indicates the [HEPES] in mM and the number in front of "C" indicates the [$CaCl_2$] in mM in the buffer.

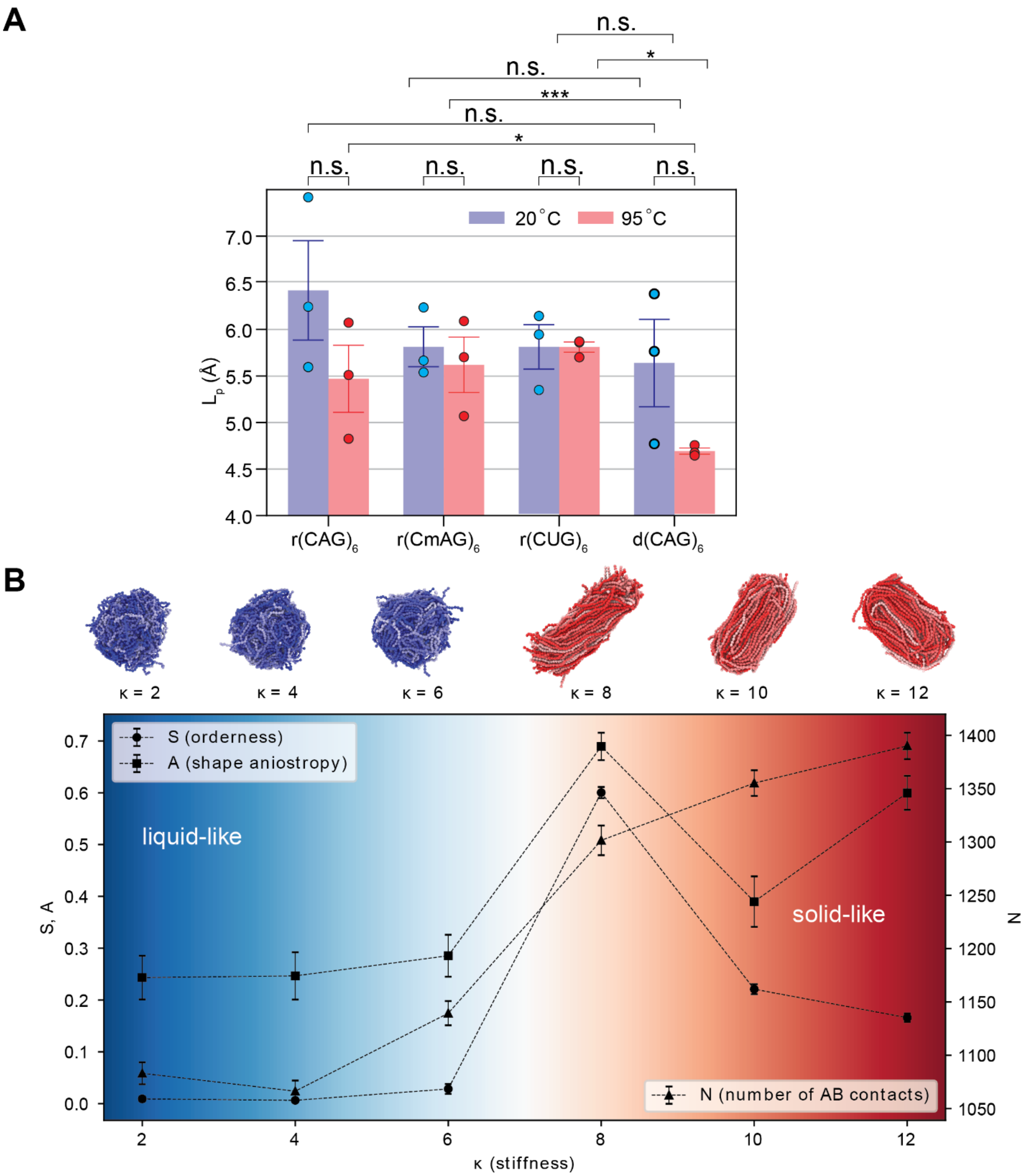


**Fig. S20. Effect of bending stiffness on condensate properties. (A)** A plot showing the persistence lengths of $(CAG)_6$ RNA and ssDNA for three independent simulations is shown. Values for the average persistence length are reported in Figure 1. P-values comparing the persistence lengths with columns numbered from right to left are p1:2 = 0.10791, p3:4 = 0.5, p5:6 = 0.31012, p7:8 = 0.058055, p1:7 = 0.16736, p2:8 = 0.049098, p3:7 = 0.37235, p4:8 = 0.018314, p5:7 = 0.37901, and p6:8 = 3.2285e-5. P-values are determined by a Student's T-test. Error bars represent s.e.m. for n = 3 data points. **(B)** Under the same conditions, minimal model simulations show that increasing the stiffness ($\kappa$) can induce a transition from a liquid-like condensate to a dynamically arrested condensate. This transition is characterized by an increase in orderness *S*, deviation of relative shape anisotropy *A,* and increase in the number of interchain crosslinks *N*. The errors are estimated using standard deviation obtained by measuring condensate properties using 1000 snapshots from the last 10,000,000 timesteps, which are sampled every 10,000 steps.

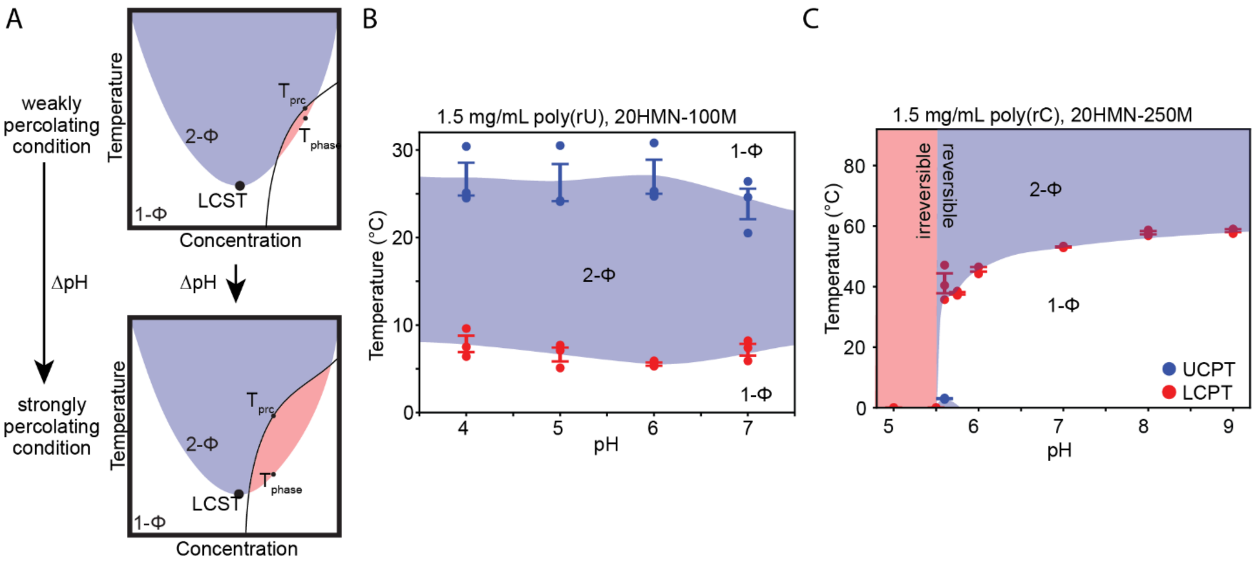


**Fig. S21. pH dependent percolation and phase separation of RNA. (A)** A schematic showing pH-dependent intersection of the percolation line with the binodal line (2-Φ regime). **(B)** A state diagram showing the pH dependence of the UCPTs and LCPTs of poly(rU) at 1.5 mg/mL in a buffer containing 20 mM each of HEPES, MES, and NaAc at the specified pH with 100 mM $MgCl_2$. The blue shading indicates the 2-Φ regime which is reversible. **(C)** A state diagram showing the pH dependence of the LCPTs of poly(rC) at 1.5 mg/mL in a buffer containing 20mM each of HEPES, MES and NaAc at the specified pH with 100 mM $MgCl_2$. The shading indicates the 2-Φ regime, and the color indicates reversibility (blue) or irreversibility (red). Buffer notation used: the number in front of "HMN" indicates the [HEPES]; [MES]; and [NaAc] in mM, and the number in front of "M" indicates [$MgCl_2$] in mM in each buffer. Error bars represent s.e.m. for n = 3 replicates.

**Supplementary Note 1**

**Small angle X-ray Scattering (SAXS) experimental design and optimization**

Our goal for SAXS experiments was to connect the coil-to-globule transition [27] at the single-chain level to the macroscopic LCST-type (Lower Critical Solution Temperature) phase behavior for RNA[17]. Explicitly, our expectation is that as the sample approaches the binodal curve from the single-phase regime as a function of increasing solution temperature, it may undergo a chain collapse known as a coil-to-globule transition (**Figure 1A**). The coil-to-globule transition of single chains has been related to UCST-type (Upper Critical Solution Temperature) phase transitions for systems where the intramolecular interactions that drive chain compaction also facilitate intermolecular multivalent interactions driving phase separation[28]. Additionally, we expect that RNA outside of the two-phase regime, but near the observed lower critical cloud point (LCPT) may form pre-percolation clusters [29,30] that may not be visible via brightfield or confocal microscopy but would be apparent in SAXS experiments as low-q bending of the q vs. I(q) curve.

In our experiments, we chose a buffer condition where SAXS measurements were performed at temperatures close to but below the LCPT for $(CAG)_{31}$ RNA and ssDNA derived from **Figure 1C**. Since $(CAG)_{31}$ was unreasonably long for atomistic simulations, for which we used $(CAG)_6$, we also performed SAXS measurements with $(CAG)_6$ RNA and ssDNA at a concentration that was approximately density matched with $(CAG)_{31}$ and in the same buffer conditions.

An additional consideration was the amount of RNA necessary to conduct the experiment. We titrated the concentration of $(CAG)_{31}$ RNA and ssDNA to determine the minimum amount of nucleic acid to be viable for SAXS measurements at the BioSAXS station at CHESS. We tried 1 μM, 5 μM, and 10 μM with only 10 μM (~0.1 mg/mL) samples providing an observable signal. During the pilot experimental run, we determined that a well-matched buffer was required and that overnight dialysis against the sample was sufficient. Furthermore, we found that the minimum volume of sample per trial needed was ~30 μL due to dithering. In experiments at elevated temperatures, we only heated the chamber with the capillary, and we allowed a few minutes for the sample to equilibrate to the temperature before acquisition.

For choice of temperatures, we chose 20 °C, 37 °C, and 50 °C to span as wide a range as possible in the single-phase regime with the limitation that the water bath could not exceed 50 °C and were slow to adjust to the temperature. Each measurement was replicated three times. Furthermore, each set of experimental data presented was collected during the same visit. However, certain experiments have not only been repeated in triplicate during a single visit but also repeated during subsequent visits including the data reported with $(CAG)_{31}$ RNA that was synthesized by two vendors, either IDT or Genscript.

**Small angle X-ray Scattering (SAXS) data controls**

We chose to perform several control experiments to verify the quality and accuracy of our results based on our expectations from SAXS data reported in the literature. These included a titration of the [NA] in the sample, a titration of $Mg^{2+}$ in the sample, and a check for $R_g$ dependence on temperature for RNA/ssDNA samples that were not observed to undergo a phase transition. Additionally, we have included the raw curve, the result of subtraction of the buffer, and a Kratky diagram for each data point from **Figure 1** as **Supplementary Note Figures SN1-SN4** and the same from **Supplementary Figure 2** as **Supplementary Note Figures SN5-SN8.** We also include example data from three different visits to CHESS (experimental runs) as **Supplementary Note Figure SN9** to verify the amelioration of low-q bending and the repeatability of the results with $r(CAG)_{31}$ as an example. In addition, we have conducted measurements on three separate visits to CHESS with three independently synthesized $(CAG)_{31}$ RNA samples from two vendors, (IDT and Genscript) which shows agreement with the chain collapse and $R_g$ values **(Supplementary Figure 3D)**.

We performed a titration of concentration for $(CAG)_6$ RNA and ssDNA (**Supplementary Figure 3.A)**, where we expected that due to the sample remaining in the single-phase (1-Φ) regime, the $R_g$ should remain constant [20]. Therefore, we repeated the temperature dependent $R_g$ measurements of $r(CAG)_6$ with the inclusion of three additional concentrations: 25 µM, 50 µM, 75 µM and 100 µM, as **Supplementary Note Figures SN10-SN21**, which also serves as an example of the repeatability of the data shown for $(CAG)_6$ in **Supplementary Note Figures SN5-SN8**. We observe that at increasing temperature, 20ºC, 37ºC, and 50 ºC, the $R_g$ values remain constant across concentration with only small variation from 25 µM – 100 µM (**Supplementary Figure S3.A**.)

To test the sensitivity of the experiment to ion concentration we performed a titration experiment with $Mg^{2+}$, which is expected to show a reduction in $R_g$ as the ion concentration is increased due to increased screening of the charge of the phosphate backbone. This has been shown previously for short homopolymers of $r(A)_{30}$ and $r(U)_{30}$ [31]. We observe that this is consistent with our observations for $r(CAG)_{31}$ shown in **Supplementary Figure S3.B** and **Supplementary Note Figures SN22-SN27**.

Finally, we conducted SAXS measurements on $r(CUU)_{31}$ and $d(CTT)_{31}$ where we observed no phase separation[17]. We rationalized that given the U/T-richness of these polymers, they may be quite far from the two-phase regime and thus there may not be any changes in the $R_g$ values upon heating/cooling within the same temperature range (20 to 50 ºC). We observe that the $R_g$ values remain constant for these two RNA/ssDNA controls (**Supplementary Figure 4; Supplementary Note Figures SN28-SN31)**.

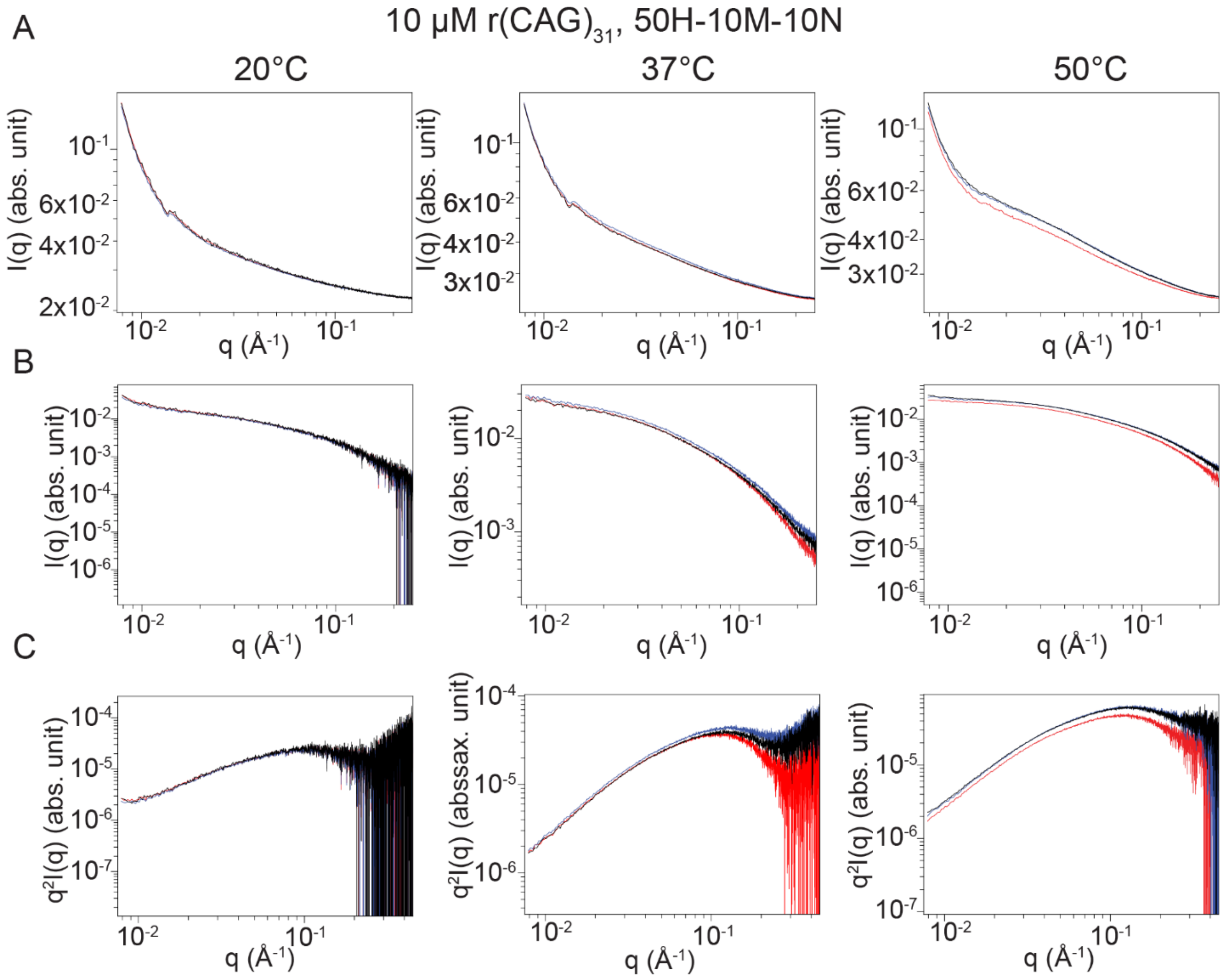


**Fig. SN1. Small Angle X-ray Scattering (SAXS) data for $(CAG)_{31}$ RNA. (A)** Temperature dependence of SAXS data for 10 µM $(CAG)_{31}$ RNA with three trials (red, blue, black) in a buffer containing 50 mM HEPES, pH 7.5 at RT, with 10 mM $MgCl_2$ and 10 mM NaCl. Data is averaged over ~100 frames and normalized to an absolute scale using water and glassy carbon. **(B)** Subtracted SAXS data from **A.** An acquisition of buffer solution including ions was averaged over ~100 frames and used for subtraction. **(C)** Kratky diagrams of the data from **B.** Buffer notation used: the number in front of "H" indicates the [HEPES] in mM, the number in front of "M" indicates the [$MgCl_2$] in mM, and the number in front of "N" indicates the [NaCl] in mM for each buffer.

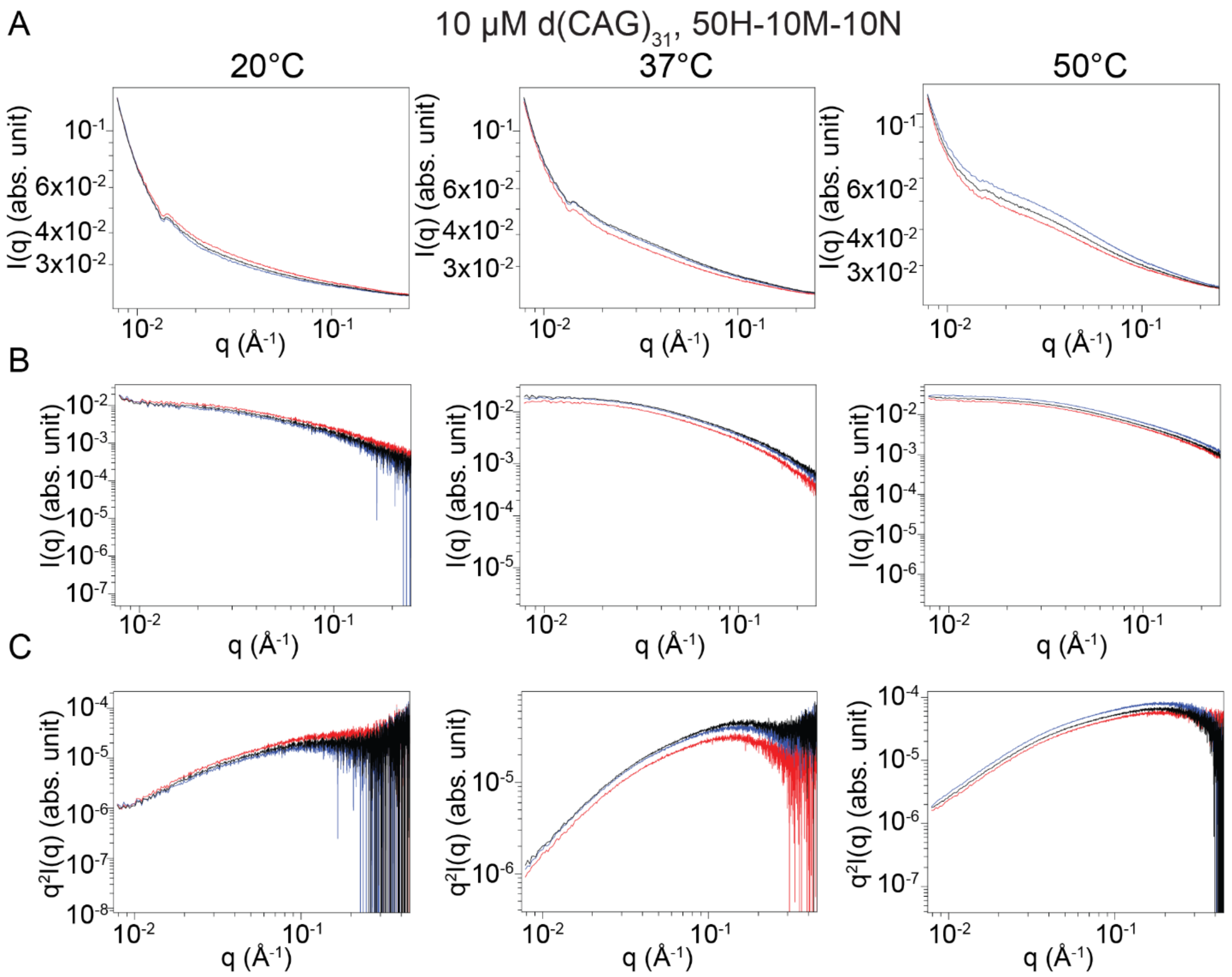


**Fig. SN2. Small Angle X-ray Scattering (SAXS) data for $(CAG)_{31}$ ssDNA. (A)** Temperature dependence of SAXS data for 10 μM $(CAG)_{31}$ DNA with three trials (red, blue, black) in a buffer containing 50 mM HEPES, pH 7.5 at RT, with 10 mM $MgCl_2$ and 10mM NaCl. Data is averaged over ~100 frames and normalized to an absolute scale using water and glassy carbon. **(B)** Subtracted SAXS data from **A.** An acquisition of buffer solution including ions was averaged over ~100 frames and used for subtraction. **(C)** Kratky diagrams of the data from **B.** Buffer notation used: the number in front of “H” indicates the [HEPES] in mM, the number in front of “M” indicates the [$MgCl_2$] in mM, and the number in front of “N” indicates the [NaCl] in mM for each buffer.

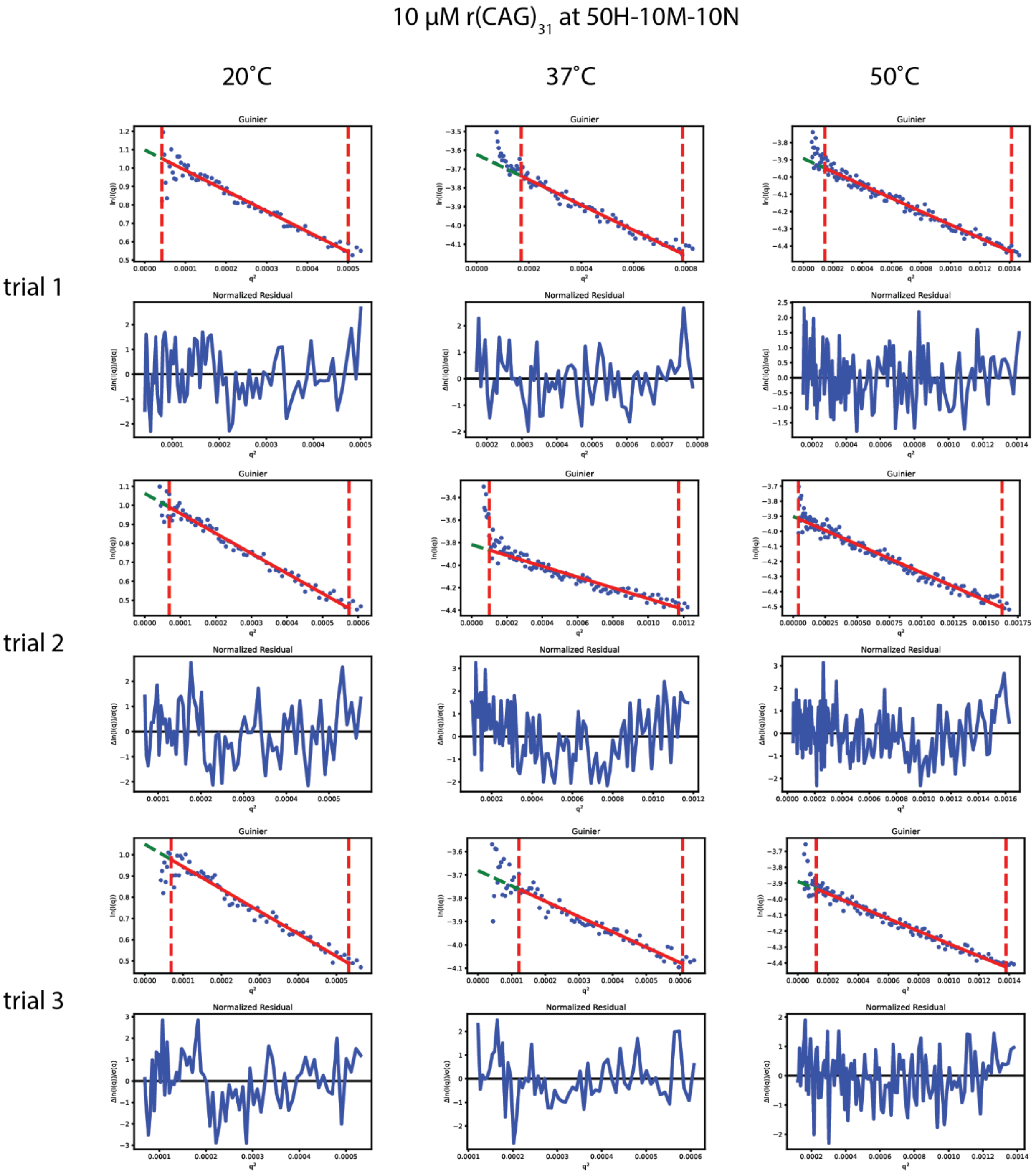


**Fig. SN3. Guinier analysis of Small Angle X-ray Scattering (SAXS) data for $(CAG)_{31}$ RNA.** Data was fit with a Guinier approximation for three temperatures and three repeats for SAXS data for 10 µM $(CAG)_{31}$ RNA with three trials (red, blue, black) in a buffer containing 50 mM HEPES, pH 7.5 at RT, with 10 mM $MgCl_2$ and 10 mM NaCl. Data is averaged over ~100 frames and normalized to an absolute scale using water and glassy carbon. Buffer notation used: the number in front of “H” indicates the [HEPES] in mM, the number in front of “M” indicates the [$MgCl_2$] in mM, and the number in front of “N” indicates the [NaCl] in mM for each buffer.

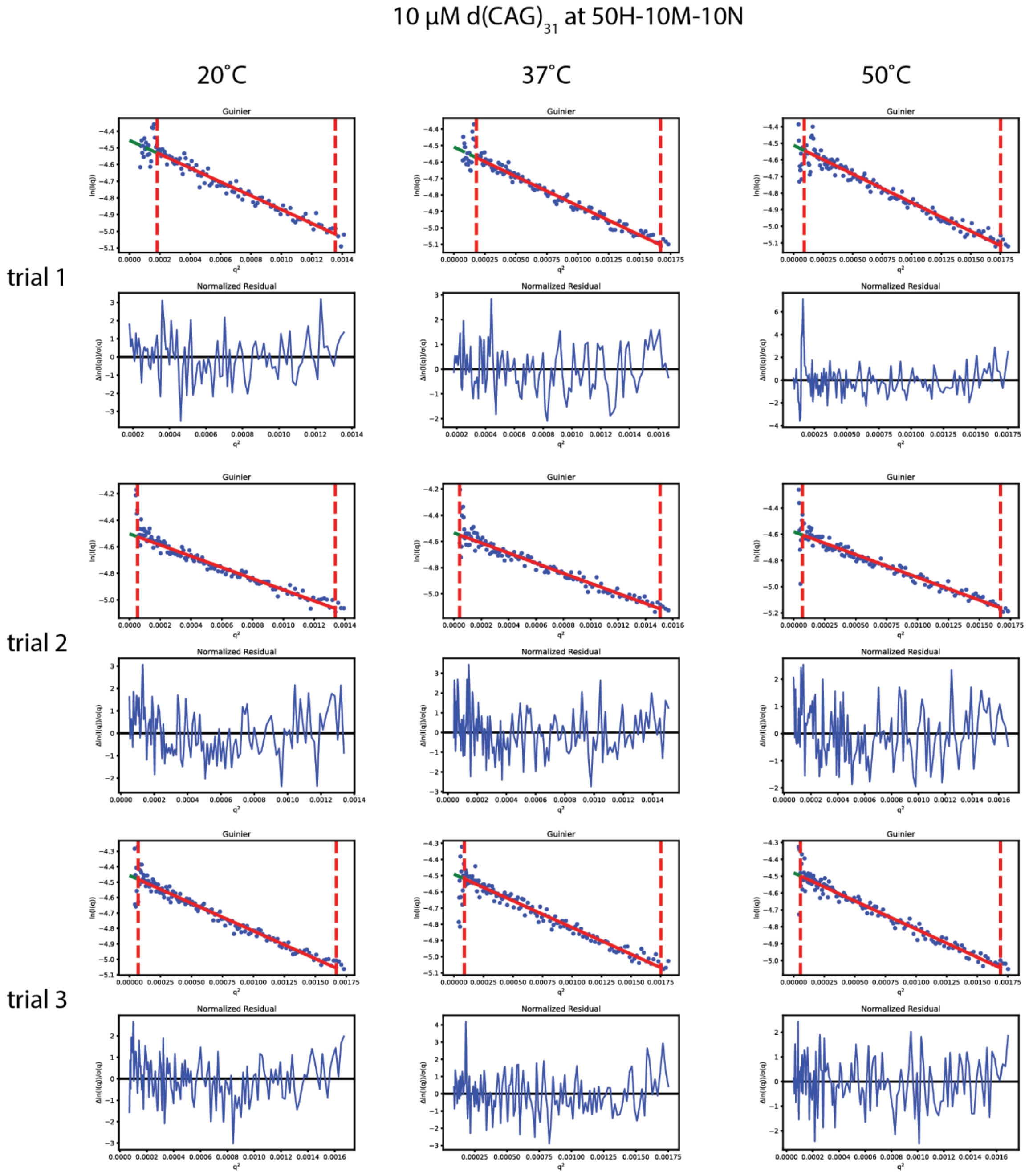


**Fig. SN4. Guinier analysis of Small Angle X-ray Scattering (SAXS) data for $(CAG)_{31}$ DNA.** Data was fit with a Guinier approximation for three temperatures and three repeats for SAXS data for 10 µM $(CAG)_{31}$ DNA with three trials (red, blue, black) in a buffer containing 50 mM HEPES, pH 7.5 at RT, with 10 mM $MgCl_2$ and 10 mM NaCl. Data is averaged over ~100 frames and normalized to an absolute scale using water and glassy carbon. Buffer notation used: the number in front of "H" indicates the [HEPES] in mM, the number in front of "M" indicates the [$MgCl_2$] in mM, and the number in front of "N" indicates the [NaCl] in mM for each buffer.

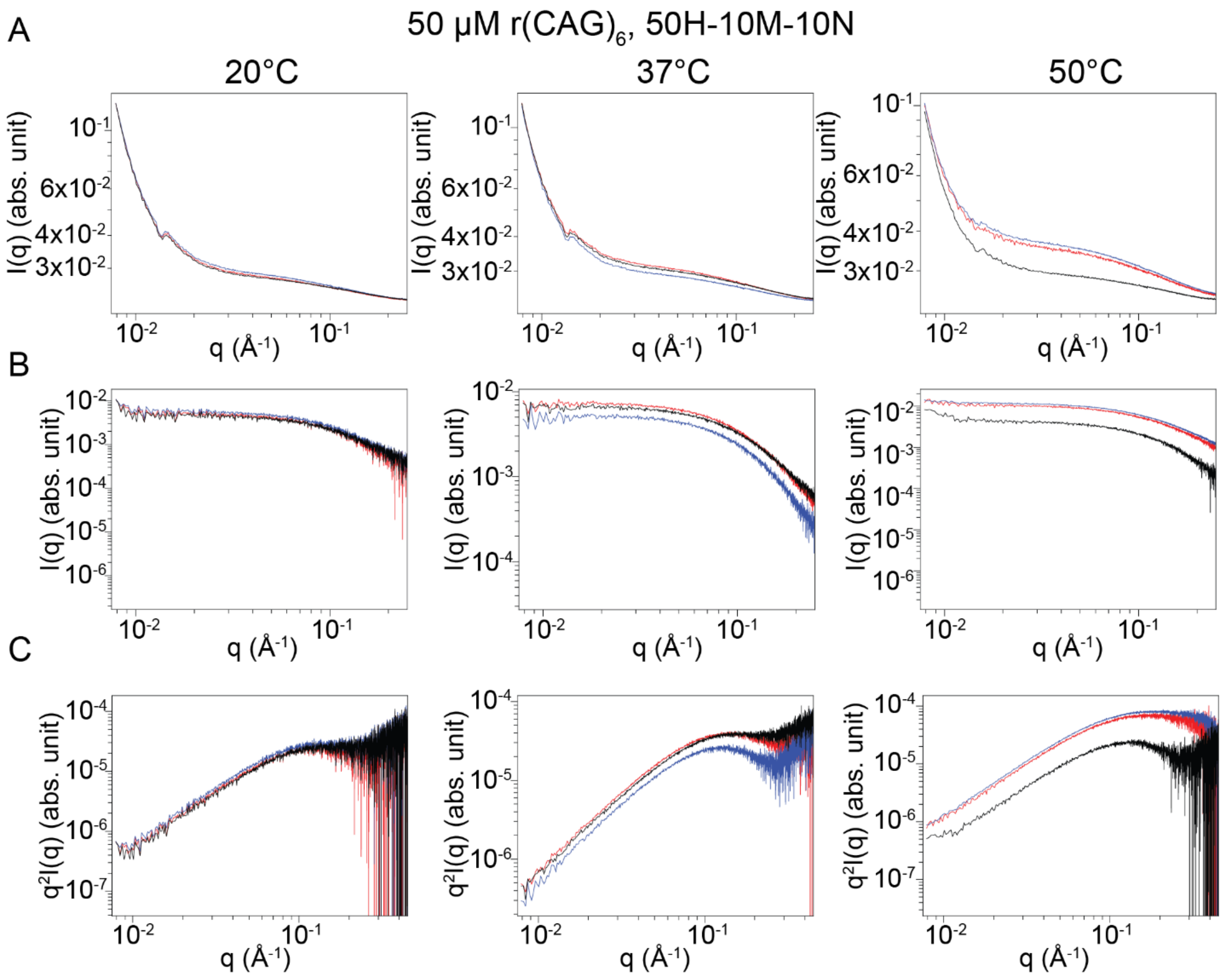


**Fig. SN5. Small Angle X-ray Scattering (SAXS) data for (CAG)$_6$ RNA. (A)** Temperature dependence of SAXS data for 50µM (CAG)$_6$ RNA with three trials (red, blue, black) in a buffer containing 50 mM HEPES, pH 7.5 at RT, with 10 mM $MgCl_2$ and 10 mM NaCl. Data is averaged over ~100 frames and normalized to an absolute scale using water and glassy carbon. **(B)** Subtracted SAXS data from **A.** An acquisition of buffer solution including ions was averaged over ~100 frames and used for subtraction. **(C)** Kratky diagrams of the data from **B.** Buffer notation used: the number in front of "H" indicates the [HEPES] in mM, the number in front of "M" indicates the [$MgCl_2$] in mM, and the number in front of "N" indicates the [NaCl] in mM for each buffer.

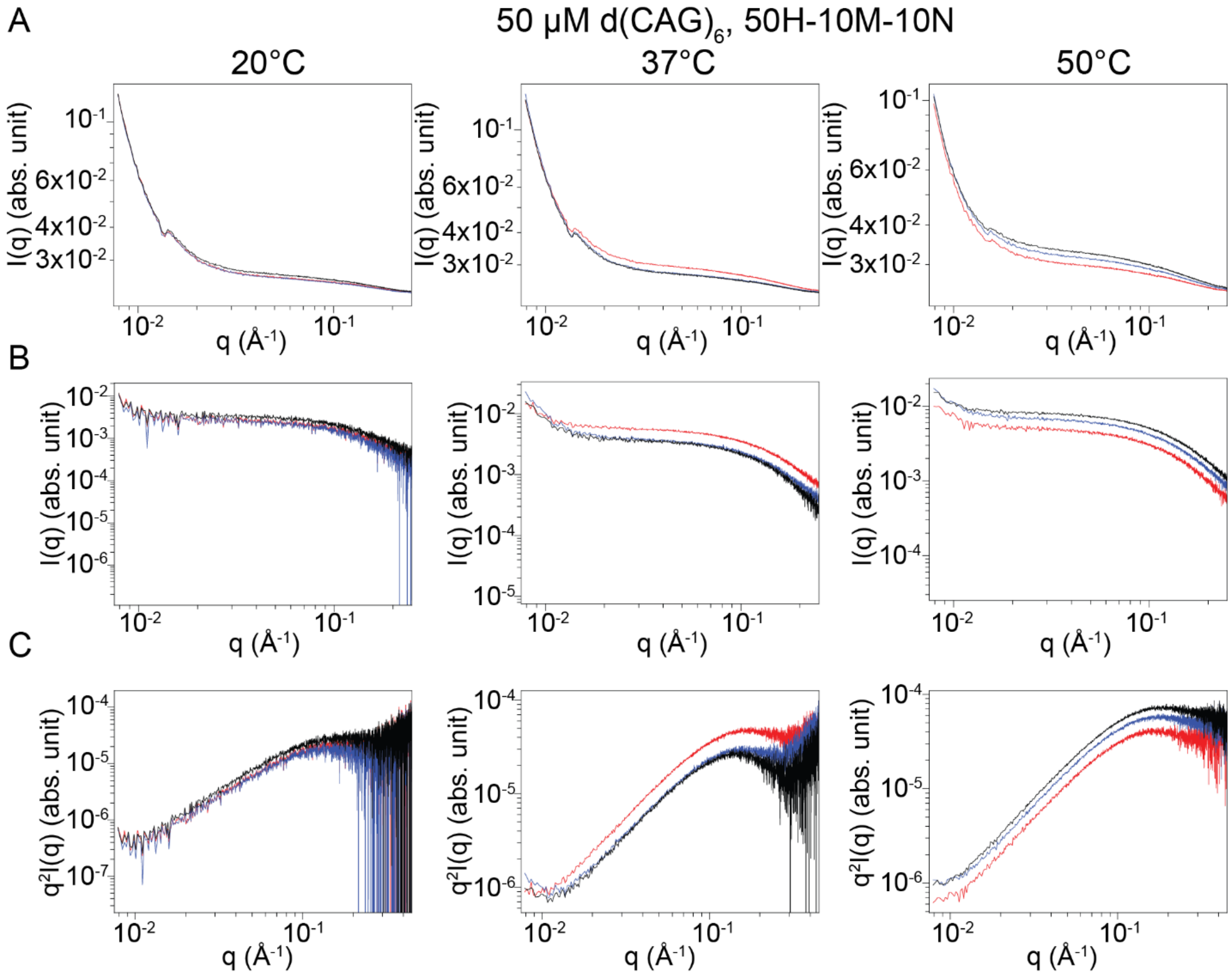


**Fig. SN6. Small Angle X-ray Scattering (SAXS) data for (CAG)$_6$ DNA. (A)** Temperature dependence of SAXS data for 50 μM (CAG)$_6$ DNA with three trials (red, blue, black) in a buffer containing 50 mM HEPES, pH 7.5 at RT, with 10 mM $MgCl_2$ and 10 mM NaCl. Data is averaged over ~100 frames and normalized to an absolute scale using water and glassy carbon. **(B)** Subtracted SAXS data from **A.** An acquisition of buffer solution including ions was averaged over ~100 frames and used for subtraction. **(C)** Kratky diagrams of the data from **B.** Buffer notation used: the number in front of "H" indicates the [HEPES] in mM, the number in front of "M" indicates the [$MgCl_2$] in mM, and the number in front of "N" indicates the [NaCl] in mM for each buffer.

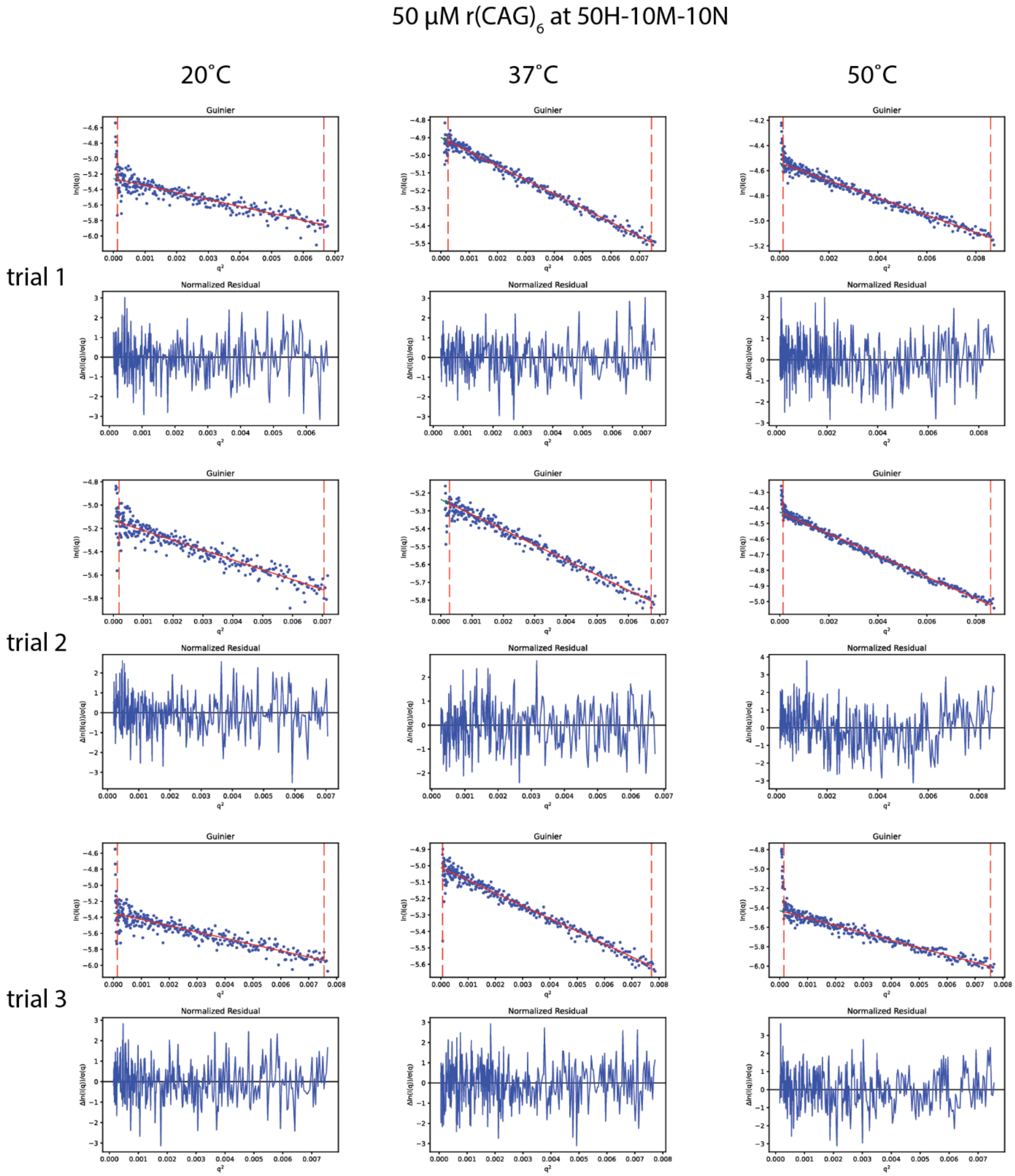


**Fig. SN7. Guinier analysis of Small Angle X-ray Scattering (SAXS) data for $(CAG)_6$ RNA.** Data was fit with a Guinier approximation for three temperatures and three repeats for SAXS data for 50 µM $(CAG)_6$ RNA with three trials (red, blue, black) in a buffer containing 50 mM HEPES, pH 7.5 at RT, with 10 mM $MgCl_2$ and 10 mM NaCl. Data is averaged over ~100 frames and normalized to an absolute scale using water and glassy carbon. Buffer notation used: the number in front of "H" indicates the [HEPES] in mM, the number in front of "M" indicates the [$MgCl_2$] in mM, and the number in front of "N" indicates the [NaCl] in mM for each buffer.

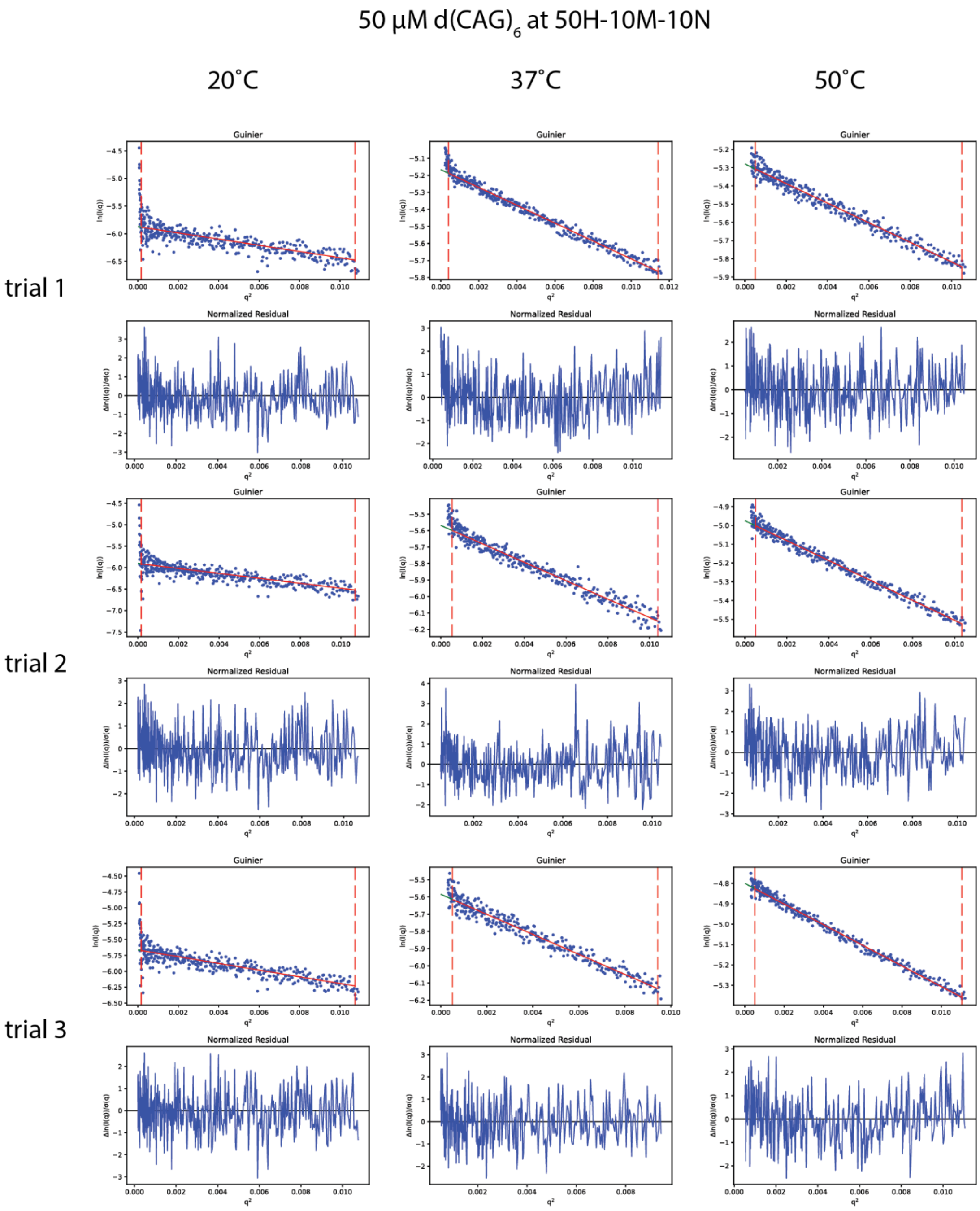


**Fig. SN8. Guinier analysis of Small Angle X-ray Scattering (SAXS) data for $(CAG)_6$ DNA.** Data was fit with a Guinier approximation for three temperatures and three repeats for SAXS data for 50 µM $(CAG)_6$ DNA with three trials (red, blue, black) in a buffer containing 50 mM HEPES, pH 7.5 at RT, with 10 mM $MgCl_2$ and 10 mM NaCl. Data is averaged over ~100 frames and normalized to an absolute scale using water and glassy carbon. Buffer notation used: the number in front of “H” indicates the [HEPES] in mM, the number in front of “M” indicates the [$MgCl_2$] in mM, and the number in front of “N” indicates the [NaCl] in mM for each buffer.

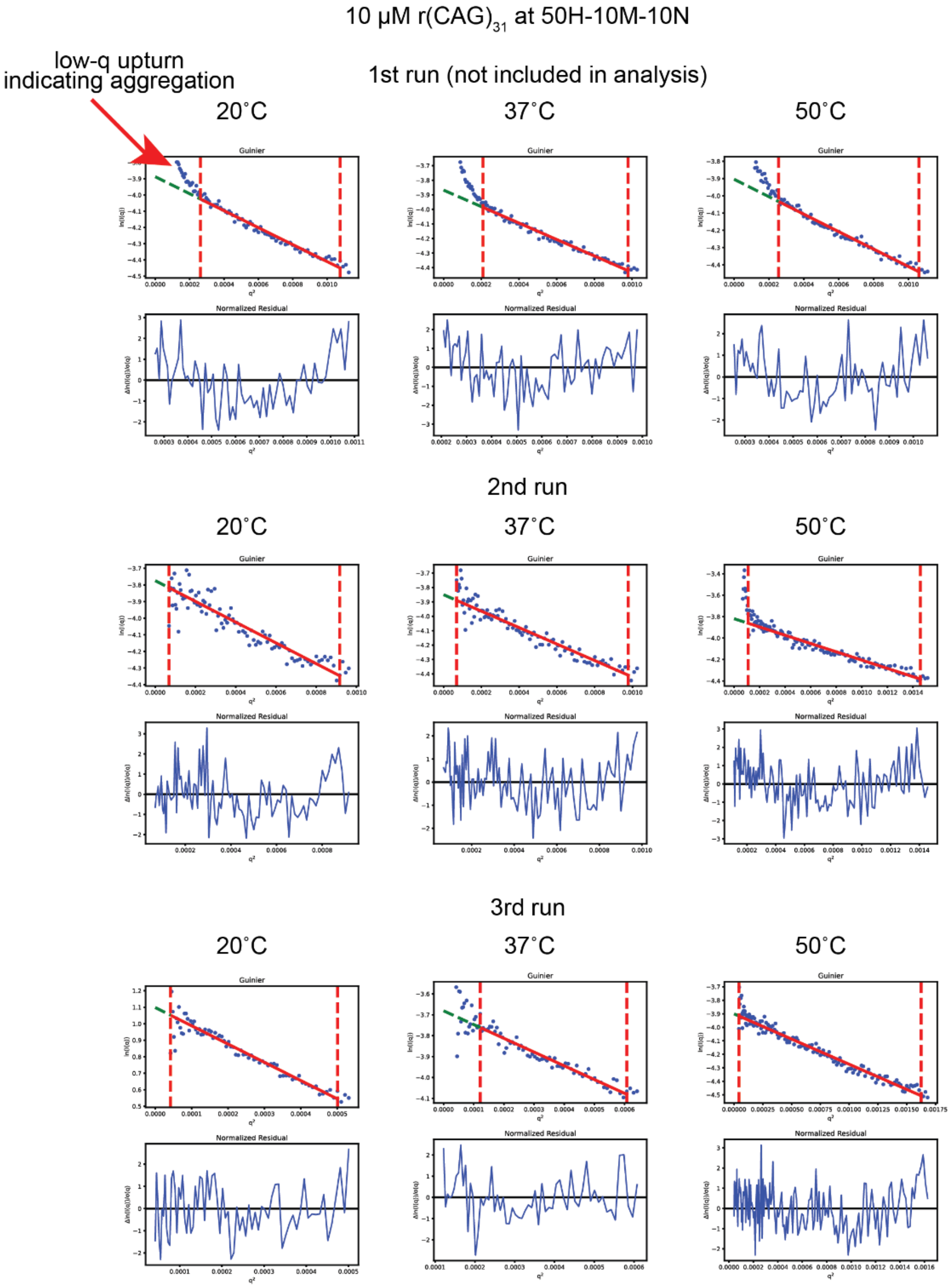


**Fig. SN9. Comparison of Guinier analysis over three different experiments for $(CAG)_{31}$ RNA.** Data was fit with a Guinier approximation for three temperatures for SAXS data for 10 μM $(CAG)_{31}$ RNA with three trials (red, blue, black) in a buffer containing 50 mM HEPES, pH 7.5 at RT, with 10 mM $MgCl_2$ and 10 mM NaCl. Data is averaged over ~100 frames and normalized to an absolute scale using water and glassy carbon. We note the upturn at low-q for the first experiment and have neglected to include that data in our analysis. In the subsequent measurements, we ameliorated the upturn, which is likely a result of RNA clustering, by lowering the beam intensity and spinning the sample prior to dialysis. Buffer notation used: the number in front of "H" indicates the [HEPES] in mM, the number in front of "M" indicates the [$MgCl_2$] in mM, and the number in front of "N" indicates the [NaCl] in mM for each buffer.

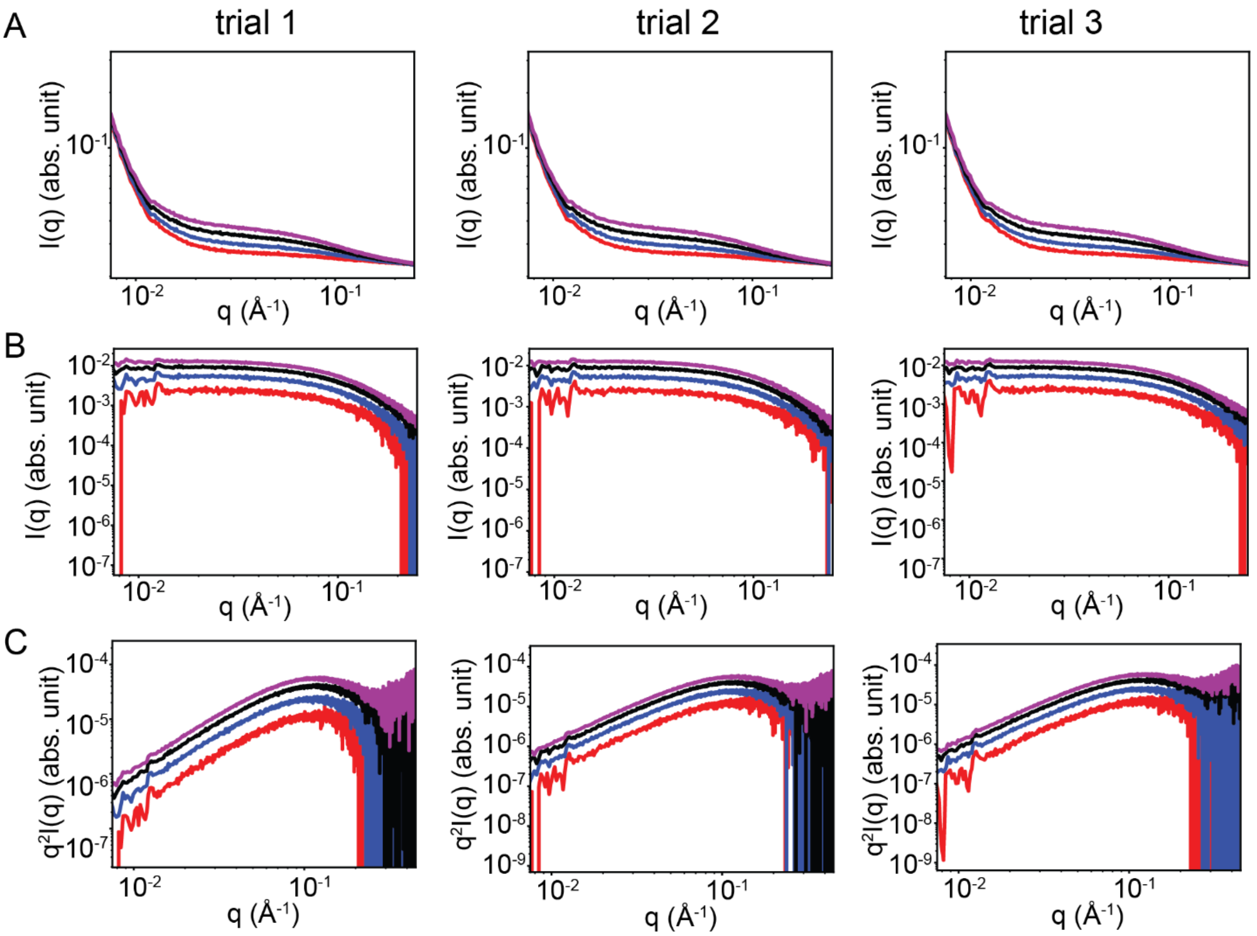


**Fig. SN10. Small Angle X-ray Scattering (SAXS) data for $(CAG)_6$ RNA at 20ºC. (A)** Temperature dependence of SAXS data for $(CAG)_6$ RNA at four concentrations (red, blue, black, purple; 25 µM, 50 µM, 75 µM and 100 µM respectively) in a buffer containing 50 mM HEPES, pH 7.5 at RT, with 10 mM $MgCl_2$ and 10 mM NaCl. Data is averaged over at least 30 frames and normalized to an absolute scale using water and glassy carbon. **(B)** Subtracted SAXS data from **A.** An acquisition of buffer solution including ions was averaged over at least 30 frames and used for subtraction. **(C)** Kratky diagrams of the data from **B.**

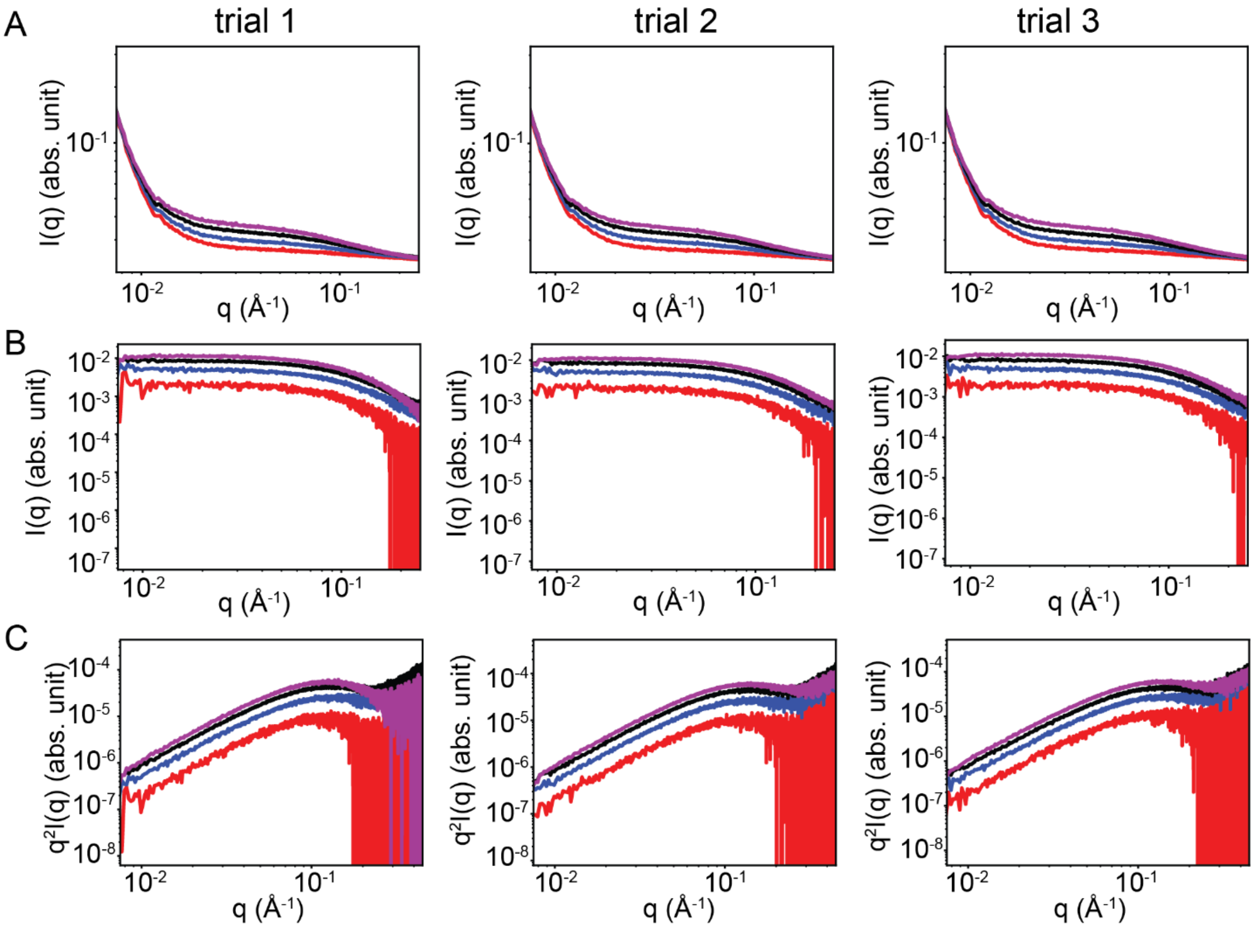


**Fig. SN11. Small Angle X-ray Scattering (SAXS) data for $(CAG)_6$ RNA at 37ºC. (A)** Temperature dependence of SAXS data for $(CAG)_6$ RNA at four concentrations (red, blue, black, purple; 25 µM, 50 µM, 75 µM and 100 µM respectively) in a buffer containing 50 mM HEPES, pH 7.5 at RT, with 10 mM $MgCl_2$ and 10 mM NaCl. Data is averaged over at least 30 frames and normalized to an absolute scale using water and glassy carbon. **(B)** Subtracted SAXS data from **A.** An acquisition of buffer solution including ions was averaged over at least 30 frames and used for subtraction. **(C)** Kratky diagrams of the data from **B.**

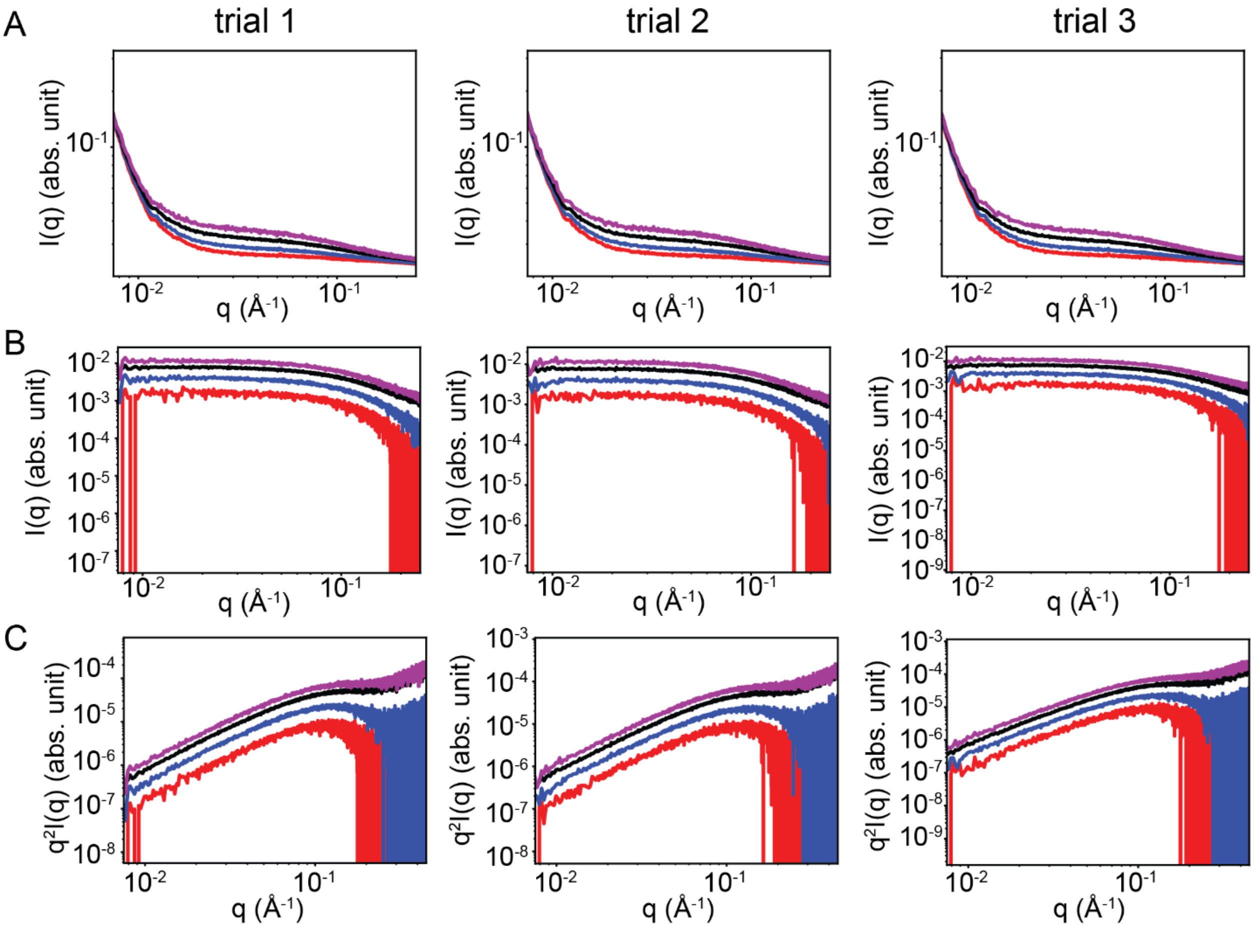


**Fig. SN12. Small Angle X-ray Scattering (SAXS) data for $(CAG)_6$ RNA at 50ºC. (A)** Temperature dependence of SAXS data for $(CAG)_6$ RNA at four concentrations (red, blue, black, purple; 25 µM, 50 µM, 75 µM and 100 µM respectively) in a buffer containing 50 mM HEPES, pH 7.5 at RT, with 10 mM $MgCl_2$ and 10 mM NaCl. Data is averaged over at least 30 frames and normalized to an absolute scale using water and glassy carbon. **(B)** Subtracted SAXS data from **A.** An acquisition of buffer solution including ions was averaged over at least 30 frames and used for subtraction. **(C)** Kratky diagrams of the data from **B.**

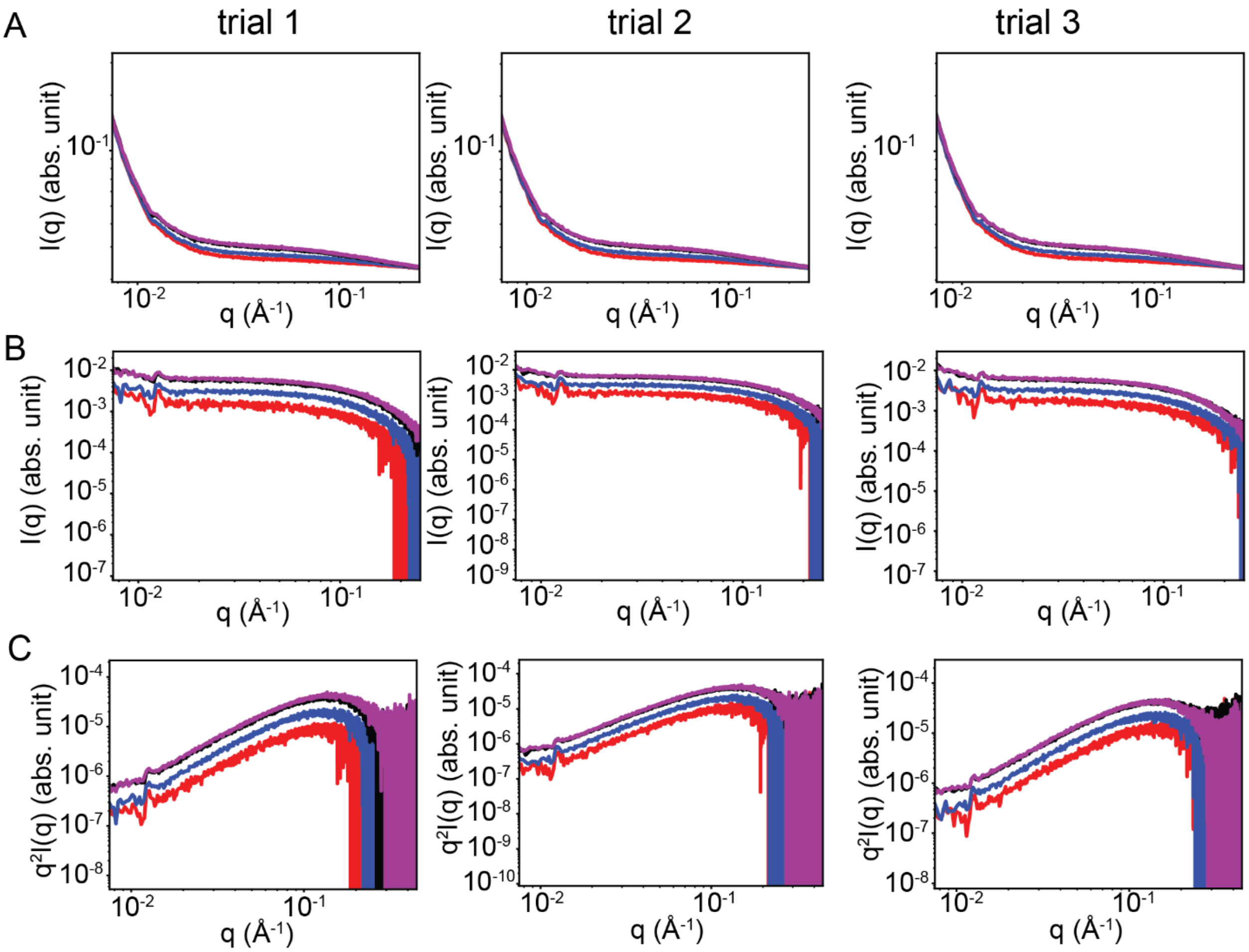


**Fig. SN13. Small Angle X-ray Scattering (SAXS) data for $(CAG)_6$ DNA at 20ºC. (A)** Temperature dependence of SAXS data for $(CAG)_6$ DNA at four concentrations (red, blue, black, purple; 25 μM, 50 μM, 75 μM and 100 μM respectively) in a buffer containing 50 mM HEPES, pH 7.5 at RT, with 10 mM $MgCl_2$ and 10 mM NaCl. Data is averaged over at least 30 frames and normalized to an absolute scale using water and glassy carbon. **(B)** Subtracted SAXS data from **A.** An acquisition of buffer solution including ions was averaged over at least 30 frames and used for subtraction. **(C)** Kratky diagrams of the data from **B.**

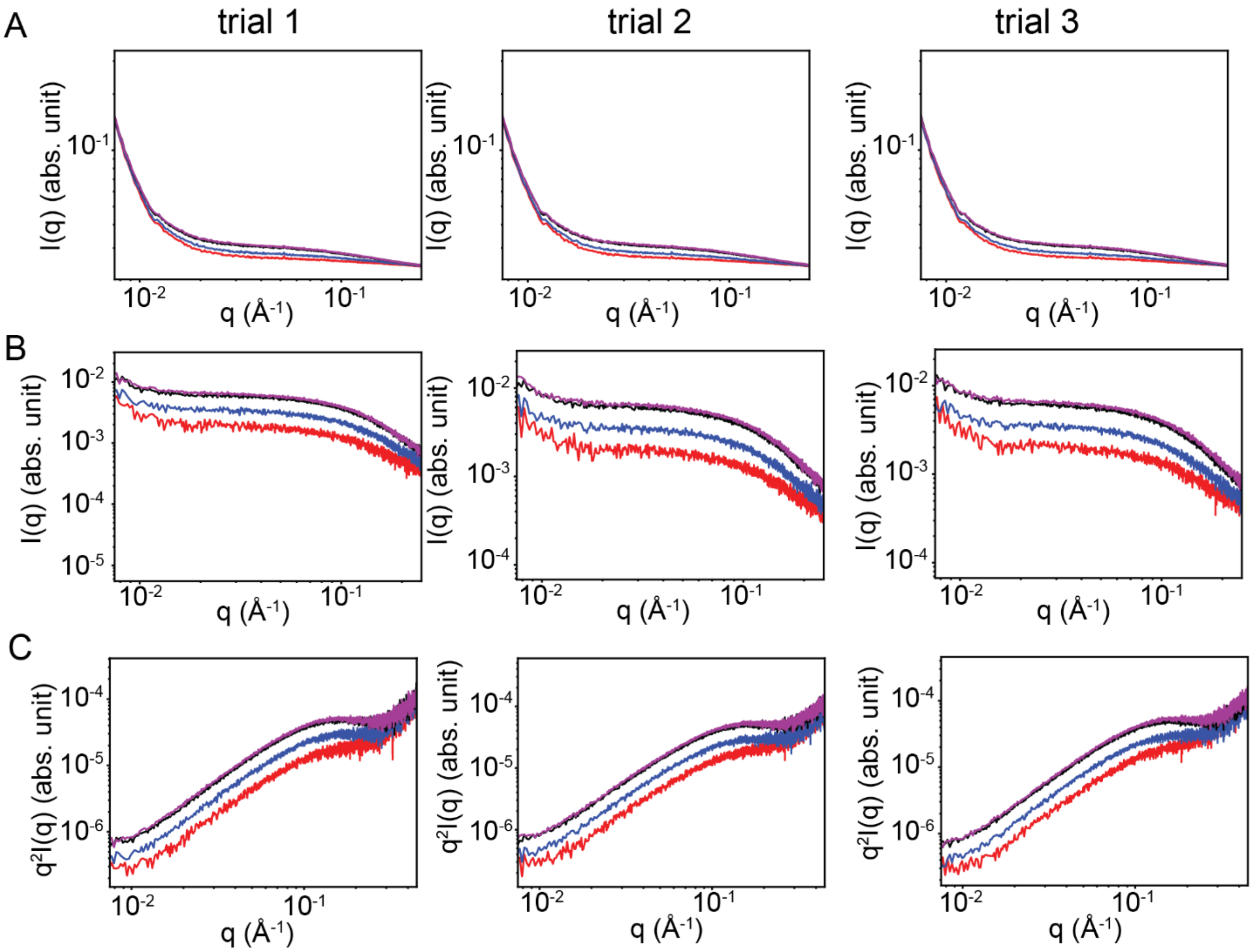


**Fig. SN14. Small Angle X-ray Scattering (SAXS) data for $(CAG)_6$ DNA at 37ºC. (A)** Temperature dependence of SAXS data for $(CAG)_6$ DNA at four concentrations (red, blue, black, purple; 25 μM, 50 μM, 75 μM and 100 μM respectively) in a buffer containing 50 mM HEPES, pH 7.5 at RT, with 10 mM $MgCl_2$ and 10 mM NaCl. Data is averaged over at least 30 frames and normalized to an absolute scale using water and glassy carbon. **(B)** Subtracted SAXS data from **A.** An acquisition of buffer solution including ions was averaged over at least 30 frames and used for subtraction. **(C)** Kratky diagrams of the data from **B.**

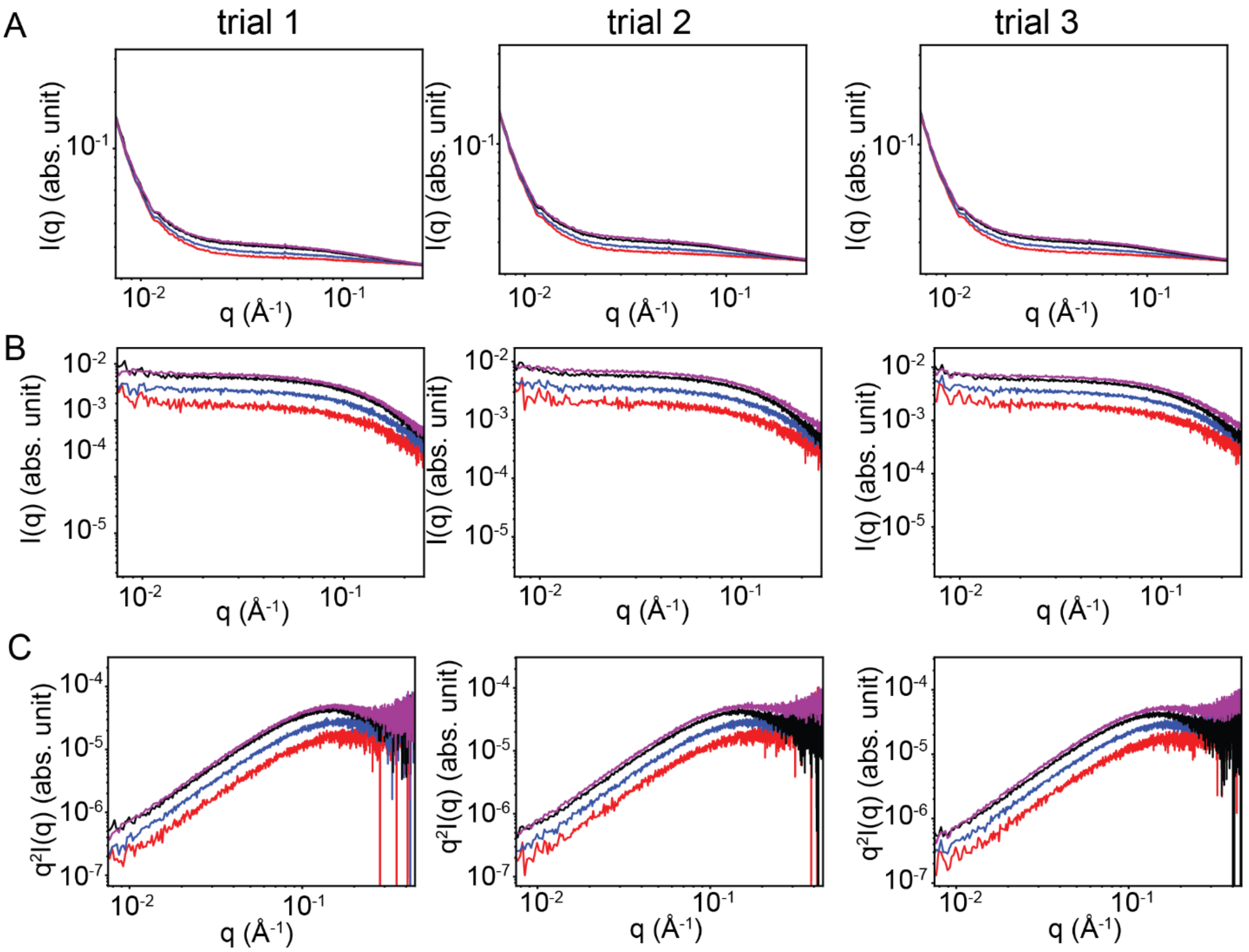


**Fig. SN15. Small Angle X-ray Scattering (SAXS) data for $(CAG)_6$ DNA at 50ºC. (A)** Temperature dependence of SAXS data for $(CAG)_6$ DNA at four concentrations (red, blue, black, purple; 25 μM, 50 μM, 75 μM and 100 μM respectively) in a buffer containing 50 mM HEPES, pH 7.5 at RT, with 10 mM $MgCl_2$ and 10 mM NaCl. Data is averaged over at least 30 frames and normalized to an absolute scale using water and glassy carbon. **(B)** Subtracted SAXS data from **A.** An acquisition of buffer solution including ions was averaged over at least 30 frames and used for subtraction. **(C)** Kratky diagrams of the data from **B.**

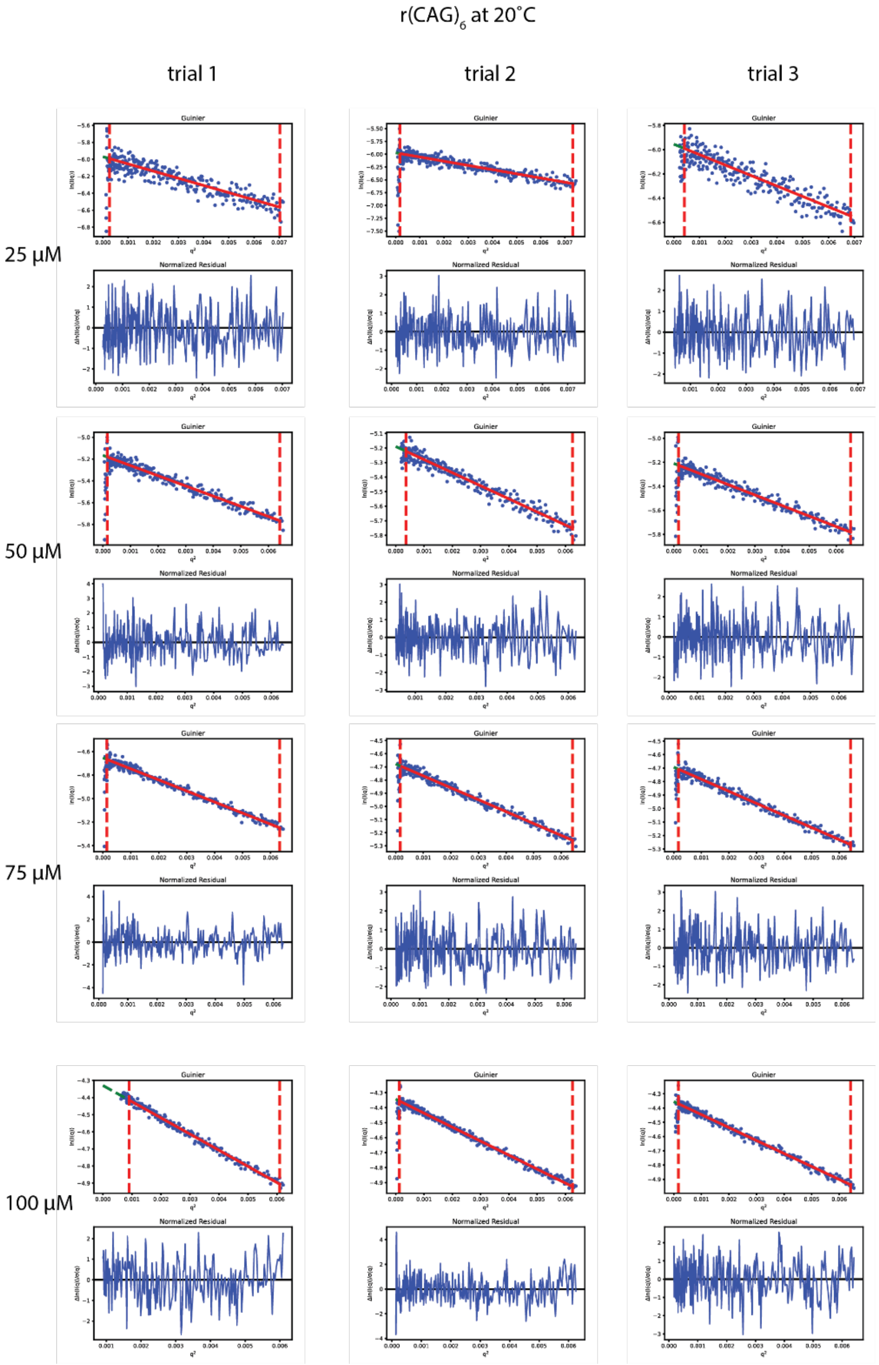


**Fig. SN16. Guinier analysis of Small Angle X-ray Scattering (SAXS) data for $(CAG)_6$ RNA at 20°C.** Data was fit with a Guinier approximation of SAXS data at four concentrations; 25 µM, 50 µM, 75 µM and 100 µM respectively for $(CAG)_6$ RNA with three trials in a buffer containing 50 mM HEPES, pH 7.5 at RT, with 10 mM $MgCl_2$ and 10 mM NaCl. Data is averaged over at least 30 frames and normalized to an absolute scale using water and glassy carbon.

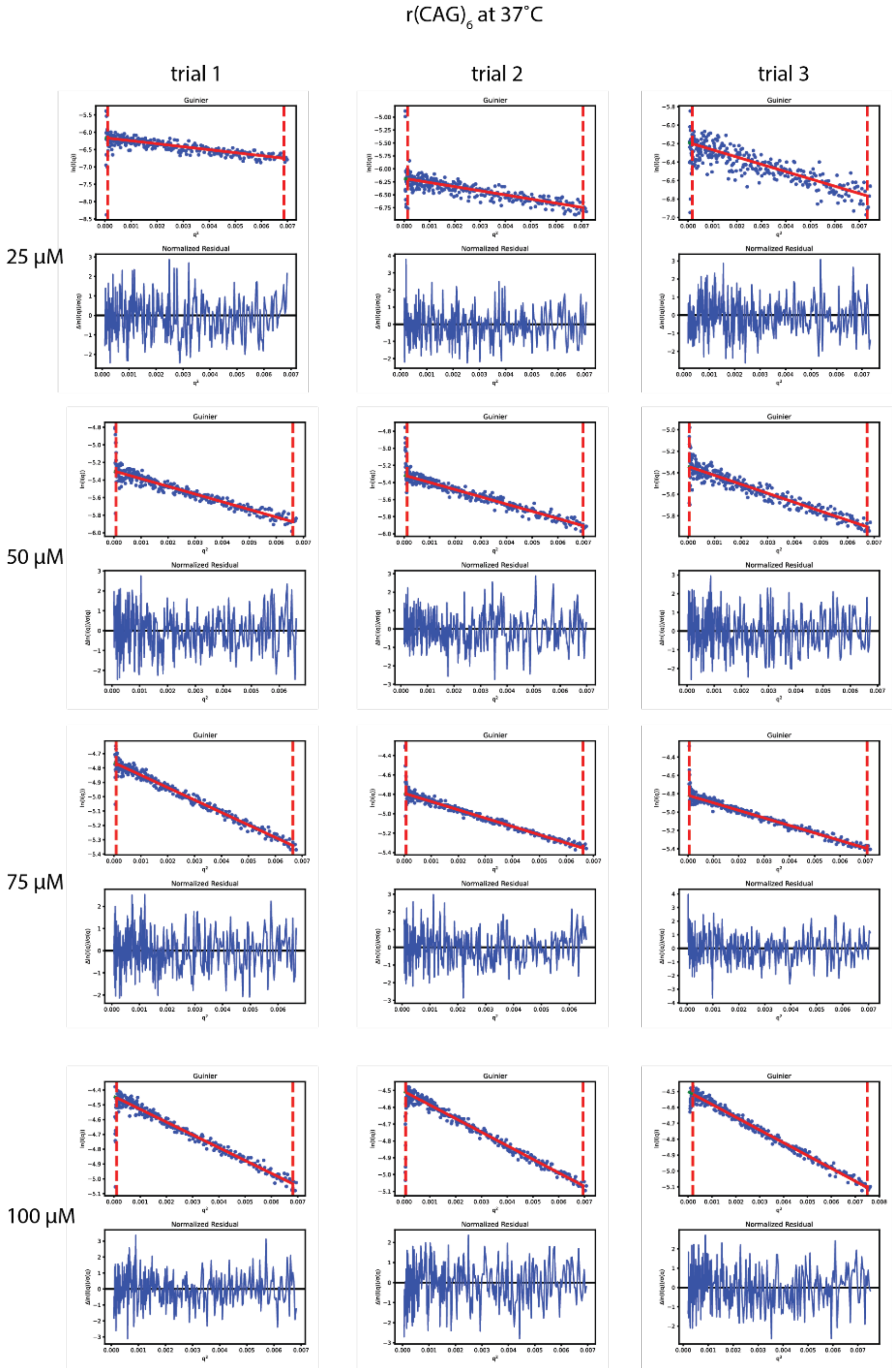


**Fig. SN17. Guinier analysis of Small Angle X-ray Scattering (SAXS) data for $(CAG)_6$ RNA at 37°C.** Data was fit with a Guinier approximation of SAXS data at four concentrations; 25 µM, 50 µM, 75 µM and 100 µM respectively for $(CAG)_6$ RNA with three trials in a buffer containing 50 mM HEPES, pH 7.5 at RT, with 10 mM $MgCl_2$ and 10 mM NaCl. Data is averaged over at least 30 frames and normalized to an absolute scale using water and glassy carbon.

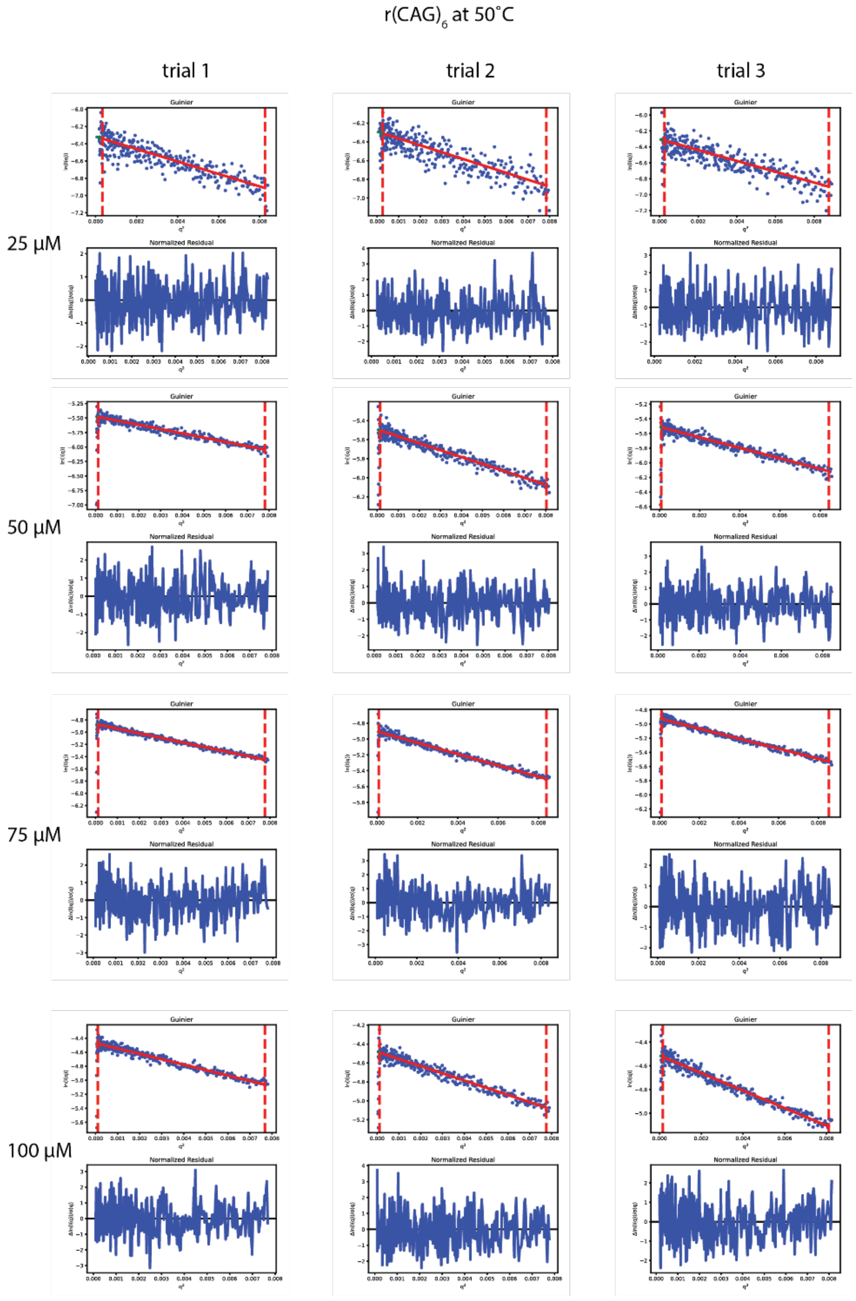


**Fig. SN18. Guinier analysis of Small Angle X-ray Scattering (SAXS) data for (CAG)$_6$ RNA at 50°C.** Data was fit with a Guinier approximation of SAXS data at four concentrations; 25 μM, 50 μM, 75 μM and 100 μM respectively for (CAG)$_6$ RNA with three trials in a buffer containing 50 mM HEPES, pH 7.5 at RT, with 10 mM $MgCl_2$ and 10 mM NaCl. Data is averaged over at least 30 frames and normalized to an absolute scale using water and glassy carbon.

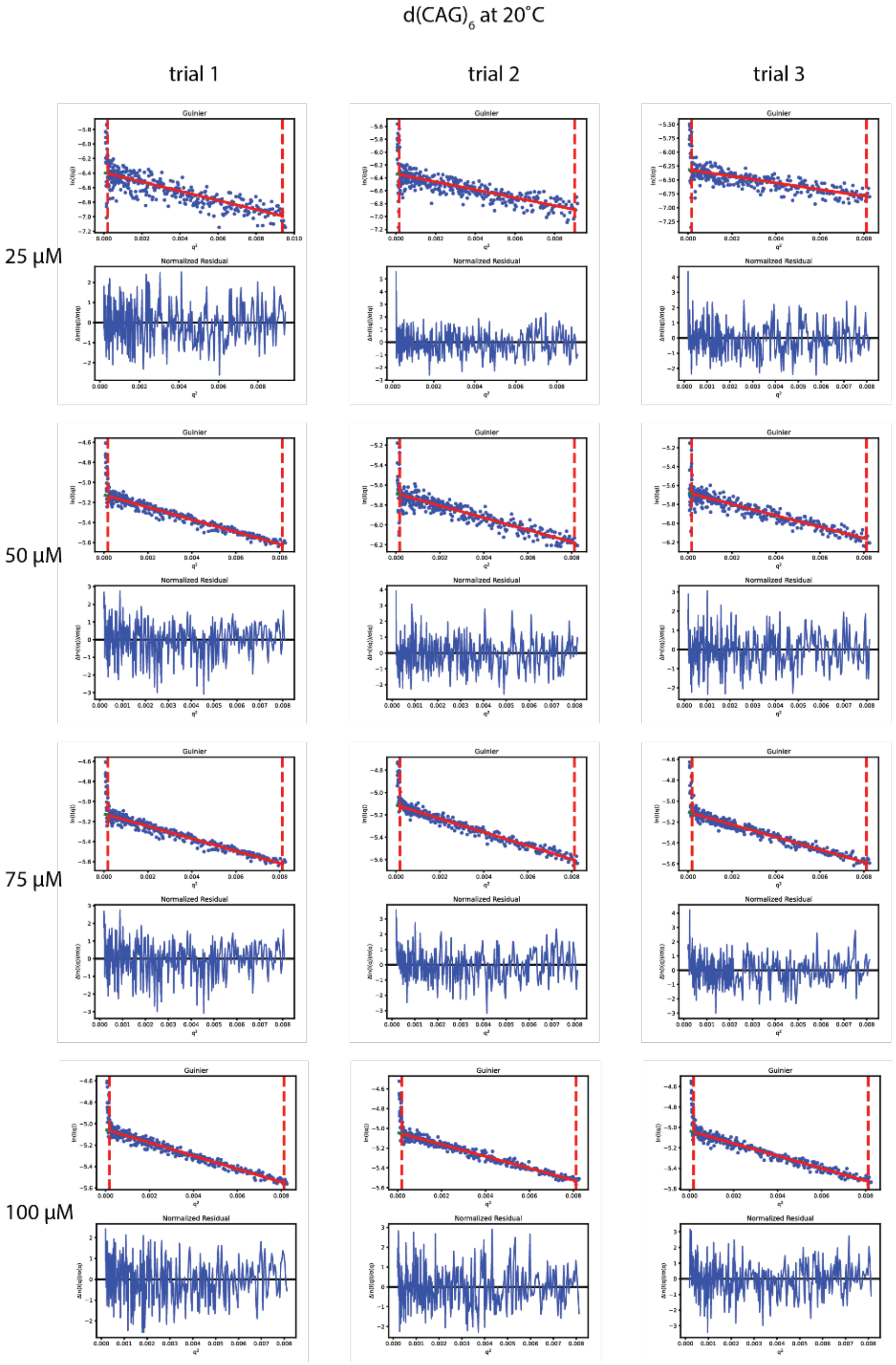


**Fig. SN19. Guinier analysis of Small Angle X-ray Scattering (SAXS) data for $(CAG)_6$ DNA at 20°C.** Data was fit with a Guinier approximation of SAXS data at four concentrations; 25 µM, 50 µM, 75 µM and 100 µM respectively for $(CAG)_6$ RNA with three trials in a buffer containing 50 mM HEPES, pH 7.5 at RT, with 10 mM $MgCl_2$ and 10 mM NaCl. Data is averaged over at least 30 frames and normalized to an absolute scale using water and glassy carbon.

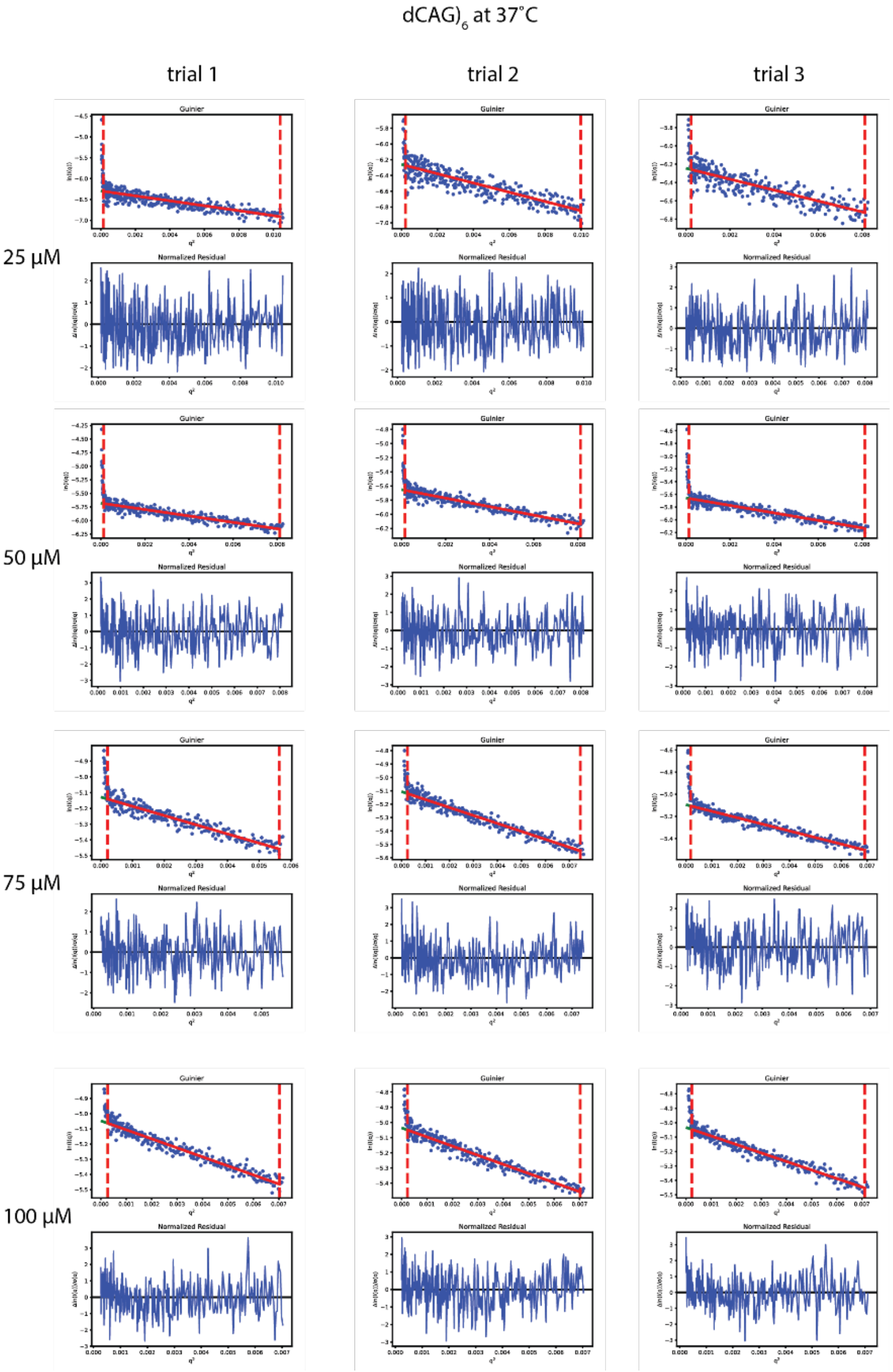


**Fig. SN20. Guinier analysis of Small Angle X-ray Scattering (SAXS) data for $(CAG)_6$ DNA at 37°C.** Data was fit with a Guinier approximation of SAXS data at four concentrations; 25 µM, 50 µM, 75 µM and 100 µM respectively for $(CAG)_6$ RNA with three trials in a buffer containing 50 mM HEPES, pH 7.5 at RT, with 10 mM $MgCl_2$ and 10 mM NaCl. Data is averaged over at least 30 frames and normalized to an absolute scale using water and glassy carbon.

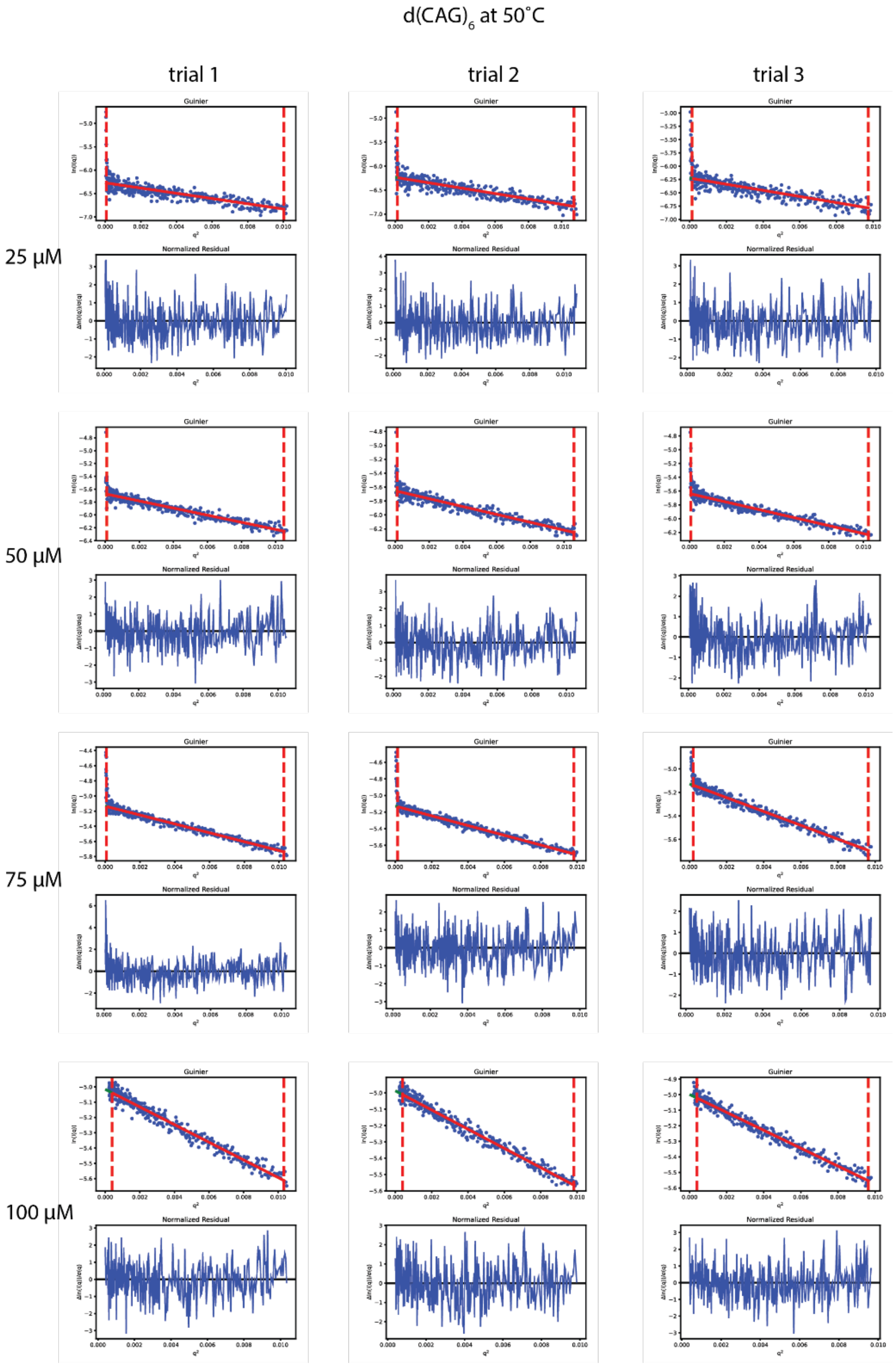


**Fig. SN21. Guinier analysis of Small Angle X-ray Scattering (SAXS) data for $(CAG)_6$ DNA at 50°C.** Data was fit with a Guinier approximation of SAXS data at four concentrations; 25 µM, 50 µM, 75 µM and 100 µM respectively for $(CAG)_6$ RNA with three trials in a buffer containing 50 mM HEPES, pH 7.5 at RT, with 10 mM $MgCl_2$ and 10 mM NaCl. Data is averaged over at least 30 frames and normalized to an absolute scale using water and glassy carbon.

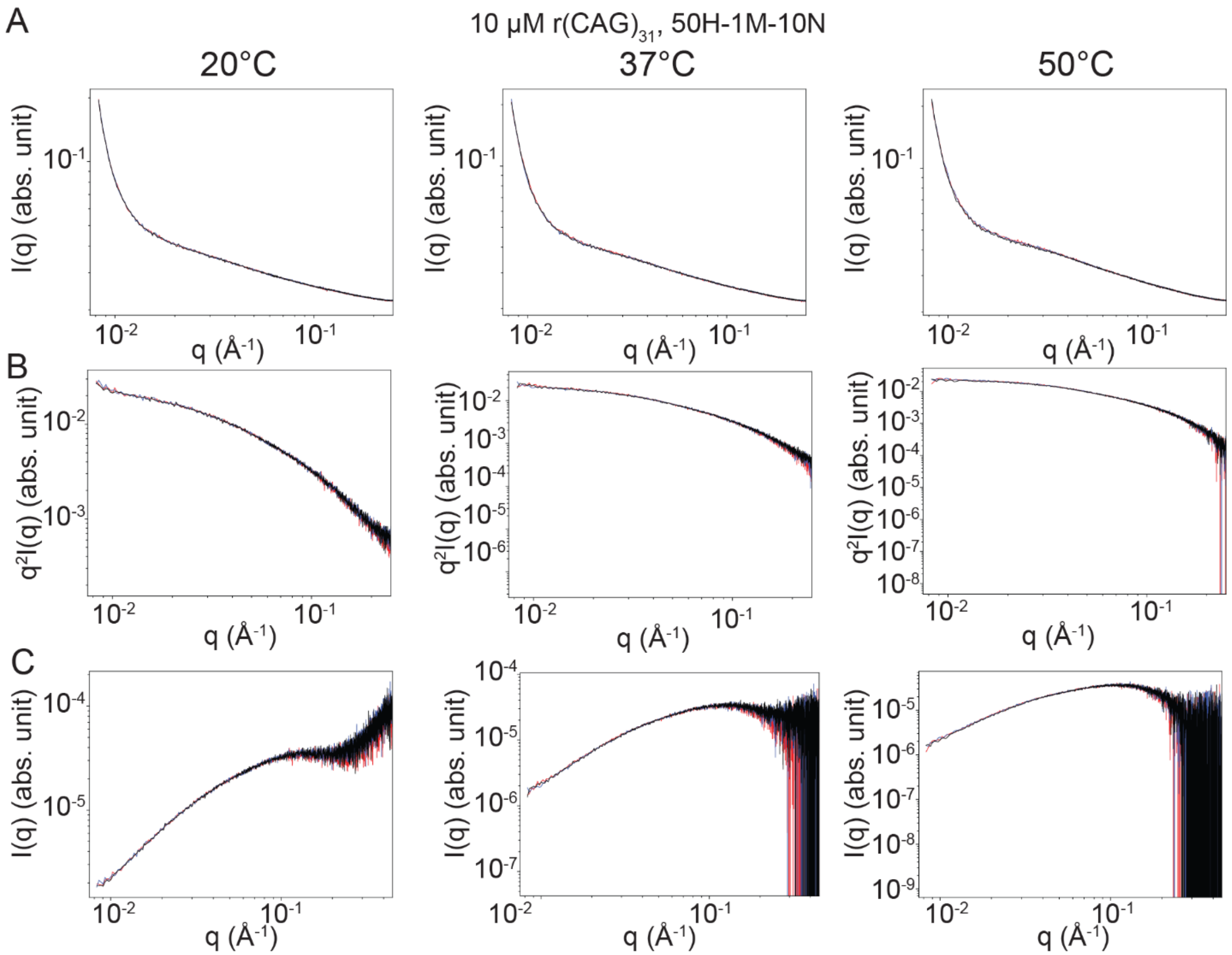


**Fig. SN22. Small Angle X-ray Scattering (SAXS) data for $(CAG)_{31}$ RNA at 1mM $MgCl_2$. (A)** Temperature dependence of SAXS data for 10 μM $(CAG)_{31}$ DNA with three trials (red, blue, black) in a buffer containing 50 mM HEPES, pH 7.5 at RT, with 1 mM $MgCl_2$ and 10 mM NaCl. Data is averaged over ~100 frames and normalized to an absolute scale using water and glassy carbon. **(B)** Subtracted SAXS data from **A.** An acquisition of buffer solution including ions was averaged over ~100 frames and used for subtraction. **(C)** Kratky diagrams of the data from **B.**

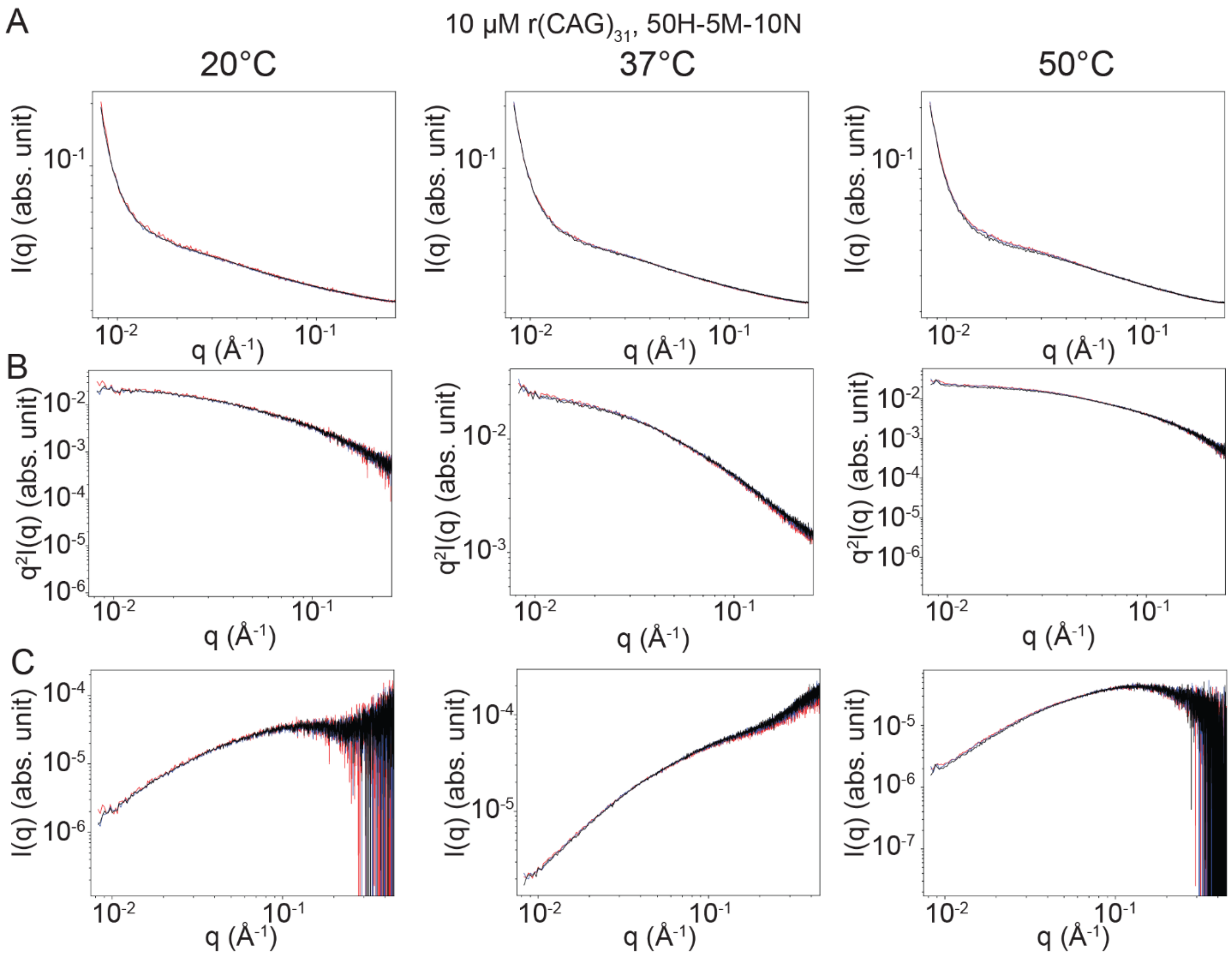


**Fig. SN23. Small Angle X-ray Scattering (SAXS) data for $(CAG)_{31}$ RNA at 5mM $MgCl_2$. (A)** Temperature dependence of SAXS data for 10 μM $(CAG)_{31}$ DNA with three trials (red, blue, black) in a buffer containing 50 mM HEPES, pH 7.5 at RT, with 5 mM $MgCl_2$ and 10 mM NaCl. Data is averaged over ~100 frames and normalized to an absolute scale using water and glassy carbon. **(B)** Subtracted SAXS data from **A.** An acquisition of buffer solution including ions was averaged over ~100 frames and used for subtraction. **(C)** Kratky diagrams of the data from **B.**

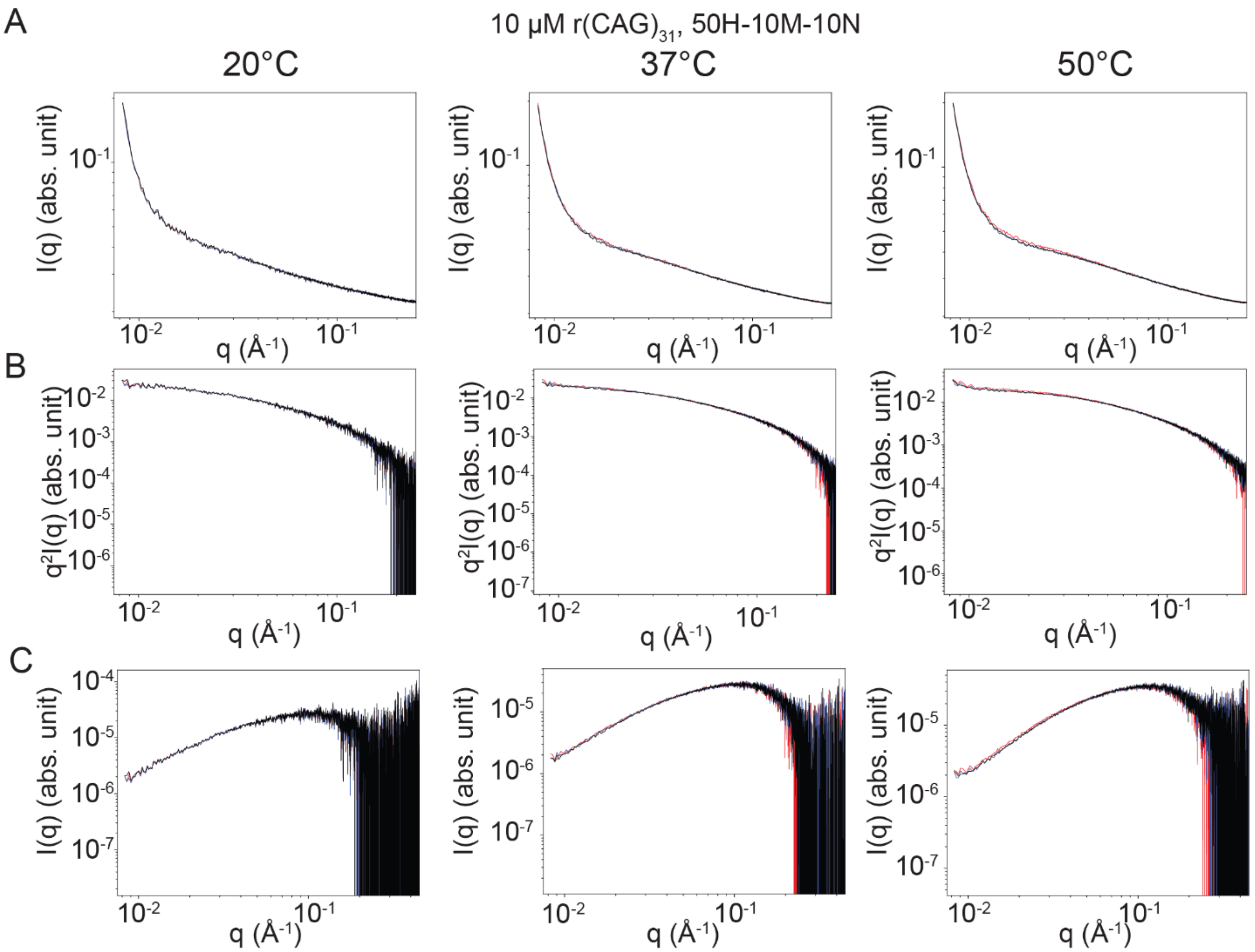


**Fig. SN24. Small Angle X-ray Scattering (SAXS) data for $(CAG)_{31}$ RNA at 10mM $MgCl_2$. (A)** Temperature dependence of SAXS data for 10 μM $(CAG)_6$ DNA with three trials (red, blue, black) in a buffer containing 50 mM HEPES, pH 7.5 at RT, with 10 mM $MgCl_2$ and 10 mM NaCl. Data is averaged over ~100 frames and normalized to an absolute scale using water and glassy carbon. Note this was collected independently from the data in Supplementary Note Figure SN1. **(B)** Subtracted SAXS data from **A.** An acquisition of buffer solution including ions was averaged over ~100 frames and used for subtraction. **(C)** Kratky diagrams of the data from **B.**

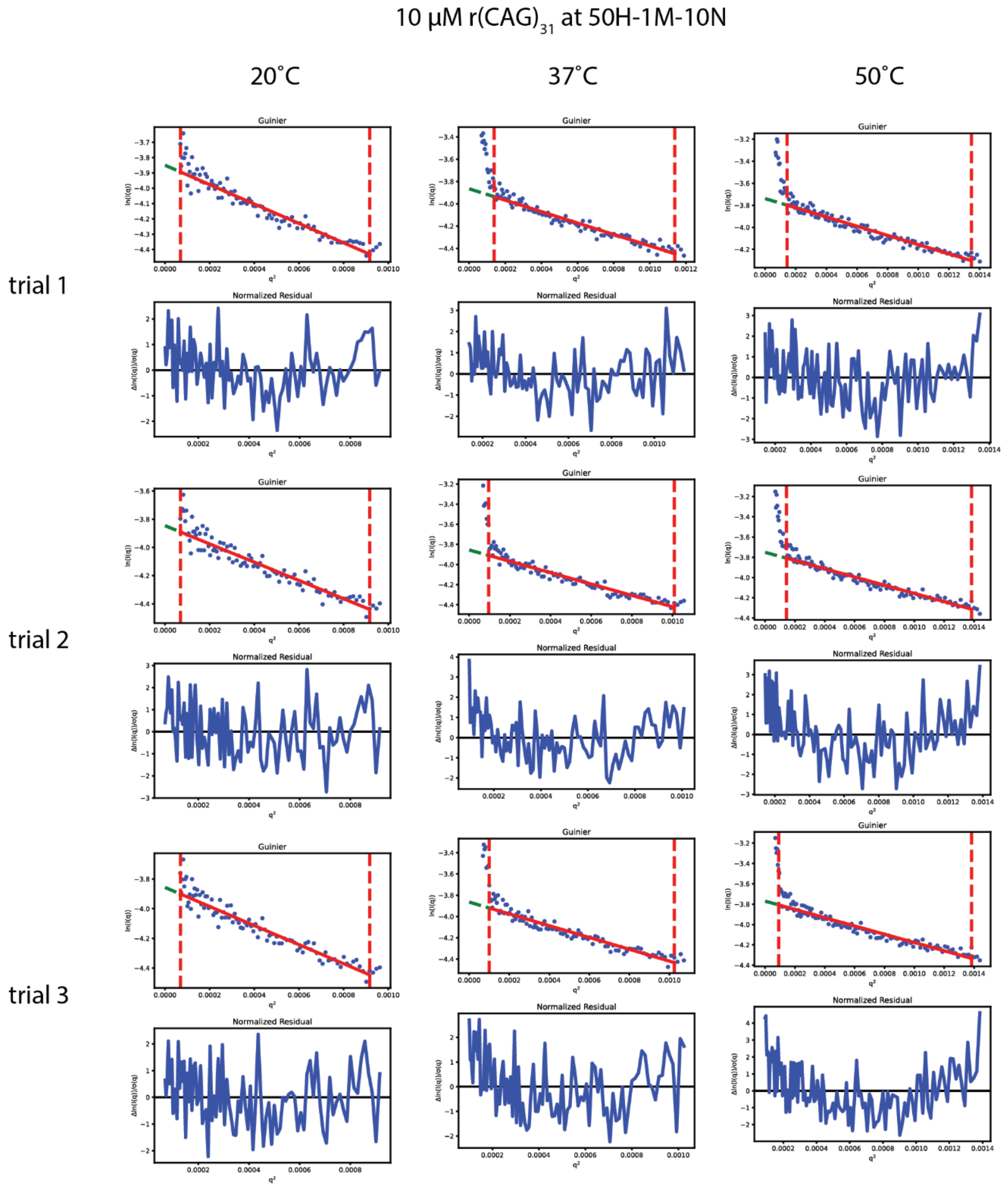


**Fig. SN25. Guinier analysis of Small Angle X-ray Scattering (SAXS) data for $(CAG)_{31}$ RNA for 1 mM $MgCl_2$.** Data was fit with a Guinier approximation for three temperatures and three repeats for SAXS data for 10 μM $(CAG)_{31}$ RNA with three trials in a buffer containing 50 mM HEPES, pH 7.5 at RT, with 1 mM $MgCl_2$ and 10 mM NaCl. Data is averaged over ~100 frames and normalized to an absolute scale using water and glassy carbon. Buffer notation used: the number in front of “H” indicates the [HEPES] in mM, the number in front of “M” indicates the [$MgCl_2$] in mM, and the number in front of “N” indicates the [NaCl] in mM for each buffer.

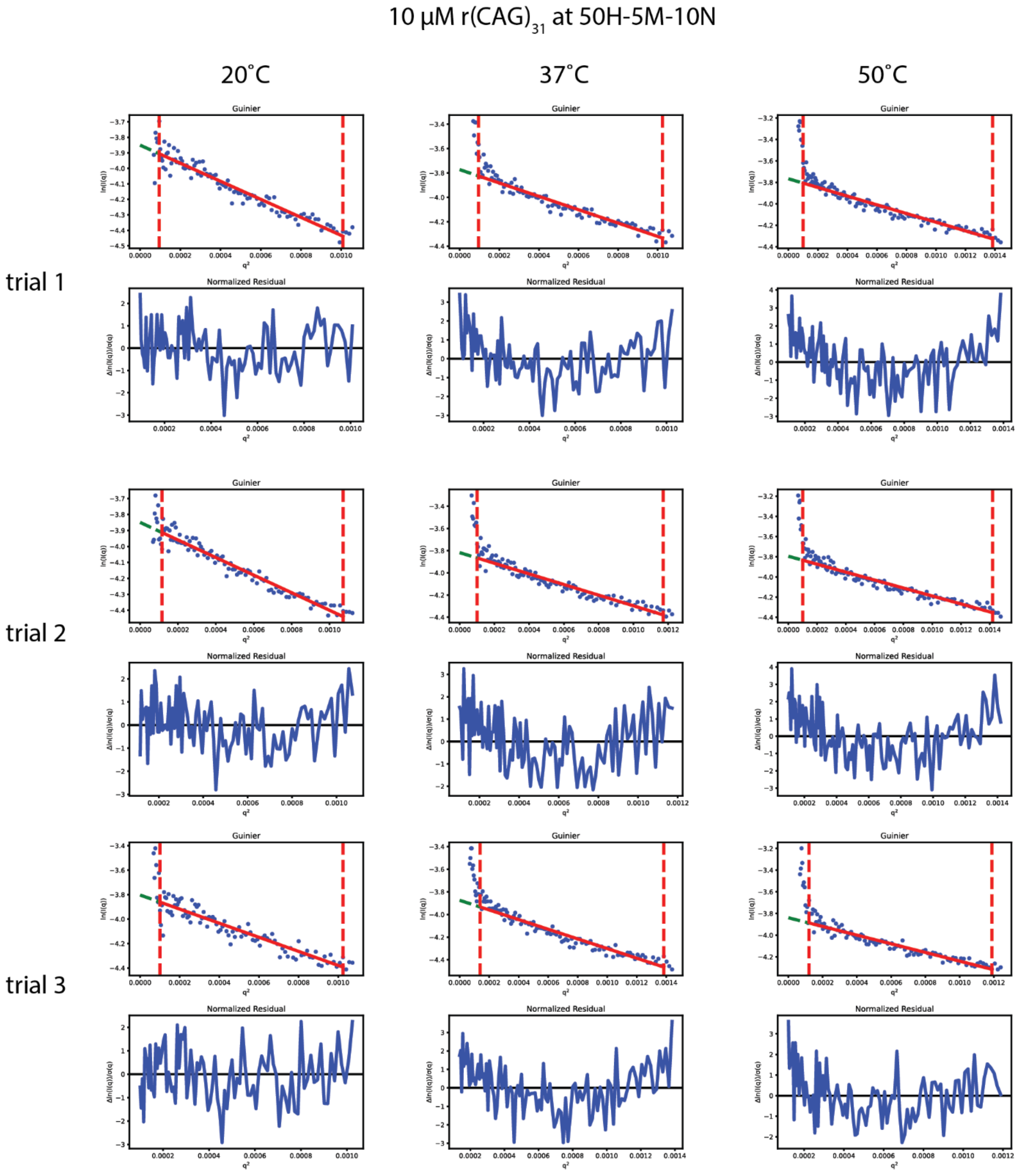


**Fig. SN26. Guinier analysis of Small Angle X-ray Scattering (SAXS) data for $(CAG)_{31}$ RNA for 5 mM $MgCl_2$.** Data was fit with a Guinier approximation for three temperatures and three repeats for SAXS data for 10 μM $(CAG)_{31}$ RNA with three trials in a buffer containing 50 mM HEPES, pH 7.5 at RT, with 5 mM $MgCl_2$ and 10 mM NaCl. Data is averaged over ~100 frames and normalized to an absolute scale using water and glassy carbon. Buffer notation used: the number in front of “H” indicates the [HEPES] in mM, the number in front of “M” indicates the [$MgCl_2$] in mM, and the number in front of “N” indicates the [NaCl] in mM for each buffer.

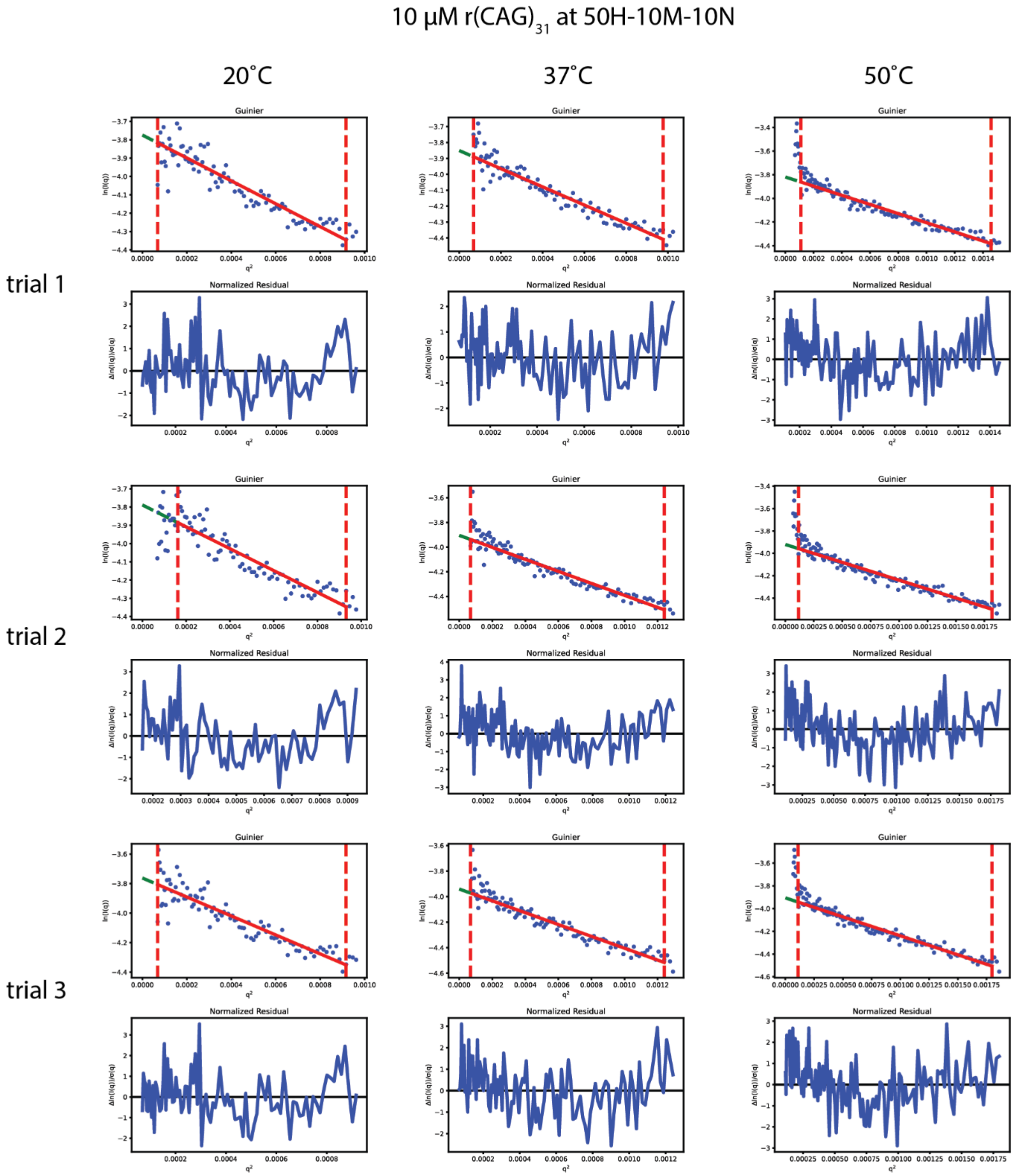


**Fig. SN27. Guinier analysis of Small Angle X-ray Scattering (SAXS) data for $(CAG)_{31}$ RNA for 10 mM $MgCl_2$.** Data was fit with a Guinier approximation for three temperatures and three repeats for SAXS data for 10 µM $(CAG)_{31}$ RNA with three trials in a buffer containing 50 mM HEPES, pH 7.5 at RT, with 10 mM $MgCl_2$ and 10 mM NaCl. Data is averaged over ~100 frames and normalized to an absolute scale using water and glassy carbon. Note this was collected independently from the data in Supplementary Note Figure SN1. Buffer notation used: the number in front of “H” indicates the [HEPES] in mM, the number in front of “M” indicates the [$MgCl_2$] in mM, and the number in front of “N” indicates the [NaCl] in mM for each buffer.

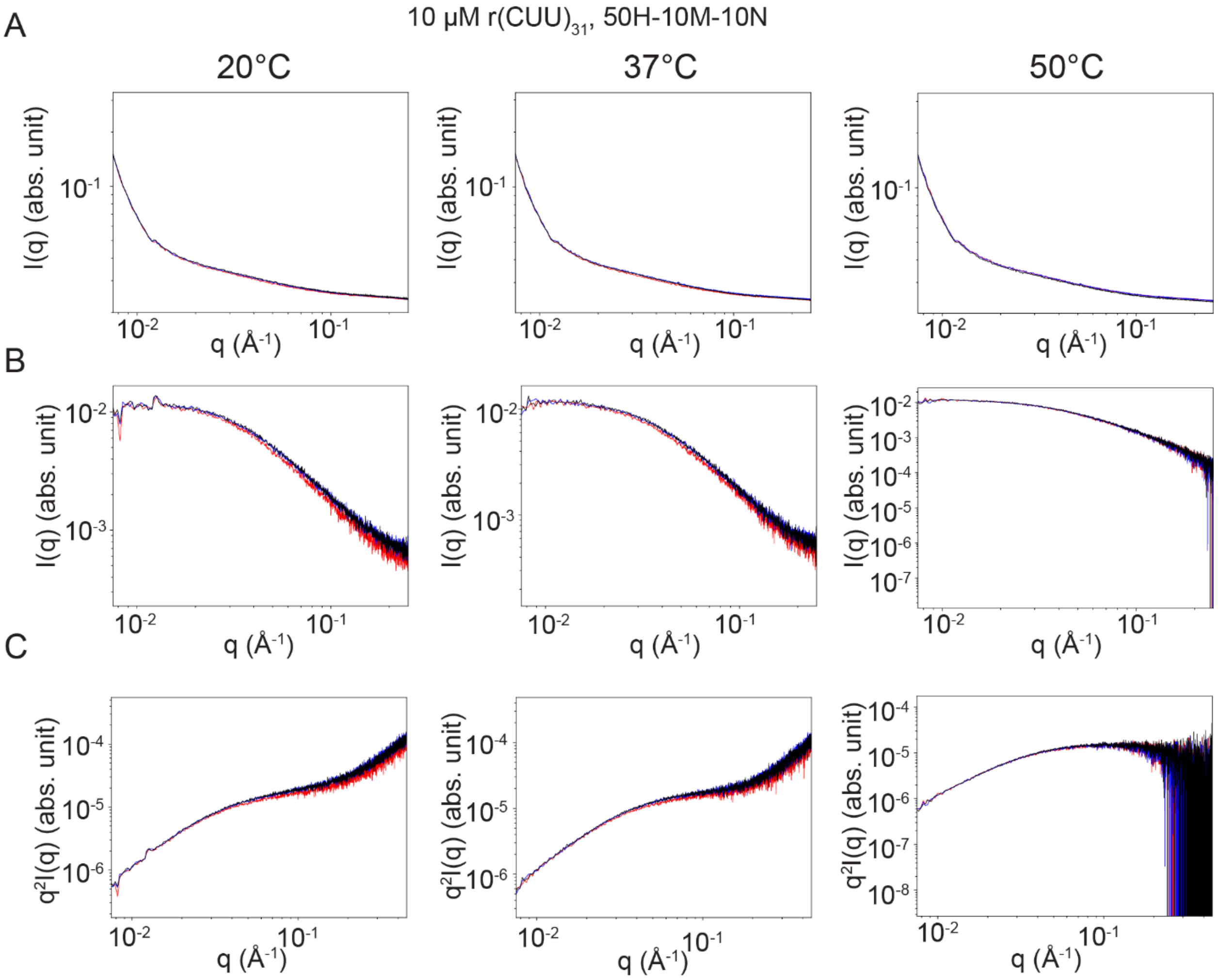


**Fig. SN28. Small Angle X-ray Scattering (SAXS) data for (CUU)$_{31}$ RNA. (A)** Temperature dependence of SAXS data for 10 μM (CUU)$_{31}$ DNA with three trials (red, blue, black) in a buffer containing 50 mM HEPES, pH 7.5 at RT, with 10 mM $MgCl_2$ and 10 mM NaCl. Data is averaged over ~100 frames and normalized to an absolute scale using water and glassy carbon. **(B)** Subtracted SAXS data from **A.** An acquisition of buffer solution including ions was averaged over ~100 frames and used for subtraction. **(C)** Kratky diagrams of the data from **B.** Buffer notation used: the number in front of "H" indicates the [HEPES] in mM, the number in front of "M" indicates the [$MgCl_2$] in mM, and the number in front of "N" indicates the [NaCl] in mM for each buffer.

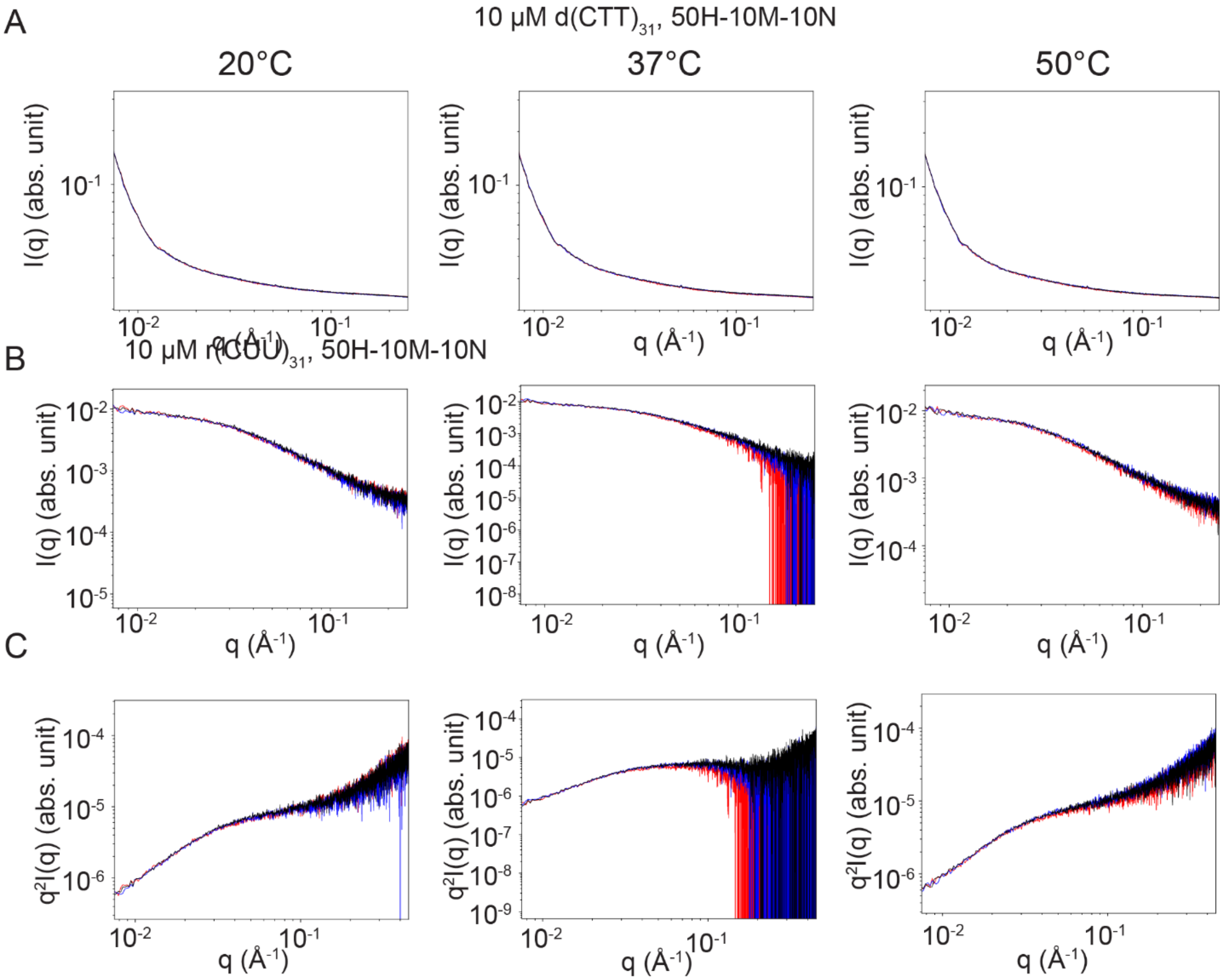


**Fig. SN29. Small Angle X-ray Scattering (SAXS) data for (CTT)$_{31}$ DNA. (A)** Temperature dependence of SAXS data for 10 μM (CTT)$_{31}$ DNA with three trials (red, blue, black) in a buffer containing 50 mM HEPES, pH 7.5 at RT, with 10 mM $MgCl_2$ and 10 mM NaCl. Data is averaged over ~100 frames and normalized to an absolute scale using water and glassy carbon. **(B)** Subtracted SAXS data from **A.** An acquisition of buffer solution including ions was averaged over ~100 frames and used for subtraction. **(C)** Kratky diagrams of the data from **B.** Buffer notation used: the number in front of "H" indicates the [HEPES] in mM, the number in front of "M" indicates the [$MgCl_2$] in mM, and the number in front of "N" indicates the [NaCl] in mM for each buffer.

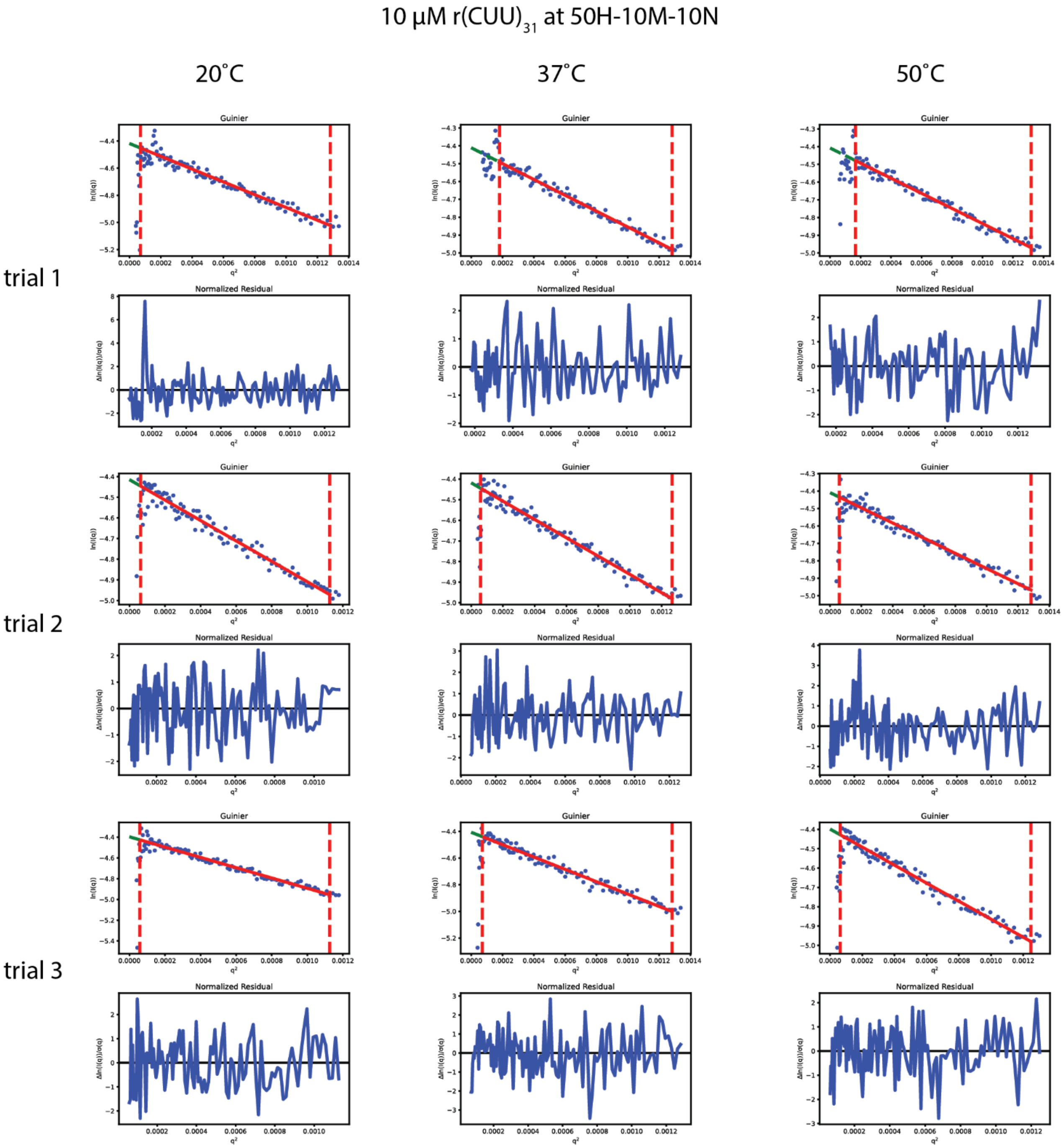


**Fig. SN30. Guinier analysis of Small Angle X-ray Scattering (SAXS) data for $(CUU)_{31}$ RNA for 5 mM $MgCl_2$.** Data was fit with a Guinier approximation for three temperatures and three repeats for SAXS data for 10 µM $(CUU)_{31}$ RNA with three trials in a buffer containing 50 mM HEPES, pH 7.5 at RT, with 10 mM $MgCl_2$ and 10 mM NaCl. Data is averaged over ~100 frames and normalized to an absolute scale using water and glassy carbon. Buffer notation used: the number in front of “H” indicates the [HEPES] in mM, the number in front of “M” indicates the $[MgCl_2]$ in mM, and the number in front of “N” indicates the [NaCl] in mM for each buffer.

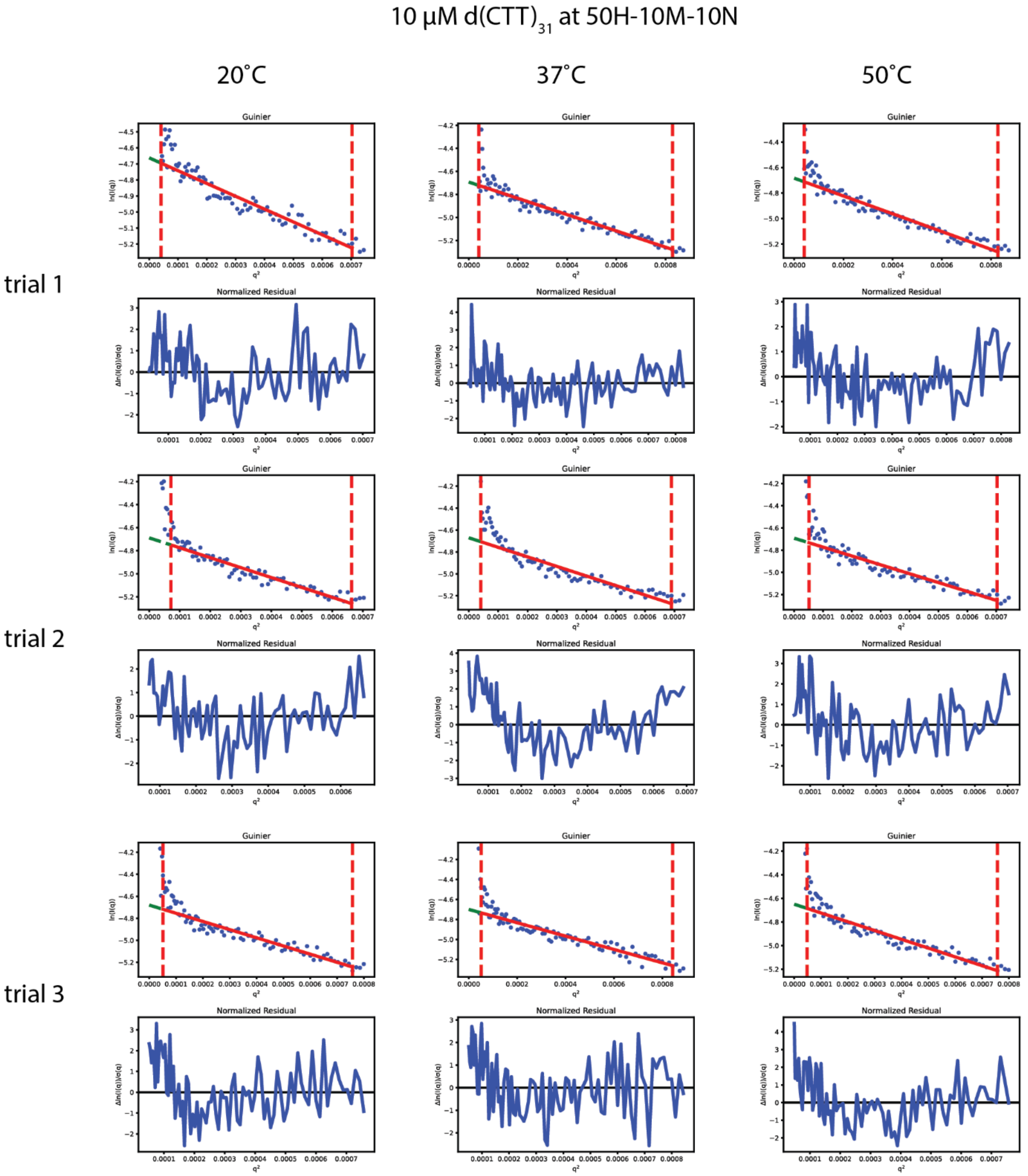


**Fig. SN31. Guinier analysis of Small Angle X-ray Scattering (SAXS) data for $(CTT)_{31}$ DNA** Data was fit with a Guinier approximation for three temperatures and three repeats for SAXS data for 10 µM $(CTT)_{31}$ DNA with three trials in a buffer containing 50 mM HEPES, pH 7.5 at RT, with 10 mM $MgCl_2$ and 10 mM NaCl. Data is averaged over ~100 frames and normalized to an absolute scale using water and glassy carbon. Buffer notation used: the number in front of "H" indicates the [HEPES] in mM, the number in front of "M" indicates the [$MgCl_2$] in mM, and the number in front of "N" indicates the [NaCl] in mM for each buffer.

| Name | Sequence | Vendor | Type |
|---|---|---|---|
| r(CAG)$_{31}$ | 5’-CAGCAGCAGCAGCAGCAGCAGCAGCAGCAGCAGCAGCAGCAGCAGCAGCAGCAGCAGCAGCAGCAG<br>CAGCAGCAGCAGCAGCAGCAGCAGCAG-3’ | IDT | RNA |
| r(CUG)$_{31}$ | 5’-CUGCUGCUGCUGCUGCUGCUGCUGCUGCUGCUGCUGCUGCUGCUGCUGCUGCUGCUGCUGCUGCUGCUG<br>CUGCUGCUGCUGCUGCUGCUGCUG-3’ | IDT | RNA |
| r(CGG)$_{31}$ | 5’-CGGCGGCGGCGGCGGCGGCGGCGGCGGCGGCGGCGGCGGCGGCGGCGGCGGCGGCGGCGGCGGCGG<br>CGGCGGCGGCGGCGGCGGCGGCGGCGG-3’ | IDT | RNA |
| r(CCG)$_{31}$ | 5’-CCGCCGCCGCCGCCGCCGCCGCCGCCGCCGCCGCCGCCGCCGCCGCCGCCGCCGCCGCCGCCG<br>CCGCCGCCGCCGCCGCCGCCGCCGCCGCCG-3’ | IDT | RNA |
| d(CAG)$_{31}$ | 5’-CAGCAGCAGCAGCAGCAGCAGCAGCAGCAGCAGCAGCAGCAGCAGCAGCAGCAGCAGCAG<br>CAGCAGCAGCAGCAGCAGCAGCAGCAGCAGCAG-3’ | IDT | DNA |
| d(CTG)$_{31}$ | 5’-CTGCTGCTGCTGCTGCTGCTGCTGCTGCTGCTGCTGCTGCTGCTGCTGCTGCTGCTGCTGCTGCTG<br>CTGCTGCTGCTGCTGCTGCTGCTGCTG-3’ | IDT | DNA |
| d(CGG)$_{31}$ | 5’-CGGCGGCGGCGGCGGCGGCGGCGGCGGCGGCGGCGGCGGCGGCGGCGGCGGCGGCGG<br>CGGCGGCGGCGGCGGCGGCGGCGGCGGCGGCGGCGG-3’ | IDT | DNA |
| d(CCG)$_{31}$ | 5’-CCGCCGCCGCCGCCGCCGCCGCCGCCGCCGCCGCCGCCGCCGCCGCCGCCGCCG<br>CCGCCGCCGCCGCCGCCGCCGCCGCCGCCGCCGCCGCCG-3’ | IDT | DNA |
| r(CAG)$_{20}$ | 5’-CAGCAGCAGCAGCAGCAGCAGCAGCAGCAGCAGCAGCAGCAGCAGCAGCAGCAGCAGCAG3’ | IDT | RNA |
| d(CAG)$_{20}$ | 5’-CAGCAGCAGCAGCAGCAGCAGCAGCAGCAGCAGCAGCAGCAGCAGCAGCAGCAGCAGCAG3’ | IDT | DNA |
| r(CmAG)$_{20}$ | 5’-CmAGCmAGCmAGCmAGCmAGCmAGCmAGCmAGCmAGCmAGCmAGCmAGCmAGCmAGCmAG<br>CmAGCmAGCmAGCmAGCmAG3’ | Gene Link | RNA |
| (TERRA)$_{10}$ | 5’-UUAGGGUUAGGGUUAGGGUUAGGGUUAGGGUUAGGGUUAGGGUUAGGGUUAGGGUUAGGG-3’ | IDT | RNA |
| (htelo)$_{10}$ | 5’-TTAGGGTTAGGGTTAGGGTTAGGGTTAGGGTTAGGGTTAGGGTTAGGGTTAGGGTTAGGG-3’ | IDT | DNA |
| d(CAG)$_{47}$ | 5’-CAGCAGCAGCAGCAGCAGCAGCAGCAGCAGCAGCAGCAGCAGCAGCAGCAGCAGCAGCAGCAGCAG<br>CAGCAGCAGCAGCAGCAGCAGCAGCAGCAGCAGCAGCAGCAGCAGCAGCAGCAGCAGCAGCAGCAGCAG<br>CAGCAG-3’ | IDT | DNA |
| r(CUG)$_{47}$ | 5’-CUGCUGCUGCUGCUGCUGCUGCUGCUGCUGCUGCUGCUGCUGCUGCUGCUGCUGCUGCUGCUGCUGCUG<br>CUGCUGCUGCUGCUGCUGCUGCUGCUGCUGCUGCUGCUGCUGCUGCUGCUGCUGCUGCUGCUGCUG<br>CUGCUG-3’ | in vitro transcription | RNA |
| r(GGGGCC)$_5$ | GGGGCCGGGGCCGGGGCCGGGGCCGGGGCC | IDT | RNA |
| d(GGGGCC)$_5$ | GGGGCCGGGGCCGGGGCCGGGGCCGGGGCC | IDT | DNA |
| rsc(CAG)$_{31}$ | GGGAA GCAGG GCGAG GGCAG CGGAC GCCGG ACAAC GGCGC CGCAA GCAGC CAGAC GAAAA GCAAG GACCC AACGG<br>AAACC CAAGC CCGCG CACAA | In vitro transcription | RNA |
| dsc(CAG)$_{31}$ | GGGAA GCAGG GCGAG GGCAG CGGAC GCCGG ACAAC GGCGC CGCAA GCAGC CAGAC GAAAA GCAAG GACCC AACGG<br>AAACC CAAGC CCGCG CACAA | IDT | DNA |
| tRNA$^{met}$ | AGCGGCGUAGGGAACUGGGAGUAUCCCGCAGGGCUCAUAACCCUGAGGUCCCUGGUUCAAAUCCAGGCGCCGCUACCA | Genscript | RNA |
| tRNA$^{met}$-2’Ome | AGCGGCGUAGGGAACUGGGAGUAUCCCGCAmGGGmCUCAUAACCCUGAGGUCCCUGGmUUmCAAAUCCAGGCGCCGCUACCA | Genscript | RNA |

**Table S1.** Table of nucleic acid sequences used in experiments.

**Movie Captions**

**Movie S1.**
Phase separation of $r(CAG)_{31}$. A sample of 100 μM $r(CAG)_{31}$ in 25 mM Tris-HCl, (pH 7.5 at 25°C) and 50 mM $MgCl_2$ underwent thermal cycling via temperature-controlled microscopy. $r(CAG)_{31}$ phase separated irreversibly with a LCPT of 40.1 ± 0.96°C (n = 3 replicates).

**Movie S2.**
Phase separation of $d(CAG)_{31}$. A sample of 100 μM $d(CAG)_{31}$ in 25 mM Tris-HCl, (pH 7.5 at 25°C) and 50 mM $MgCl_2$ underwent thermal cycling via temperature-controlled microscopy. $d(CAG)_{31}$ phase separated reversibly with a LCPT of 55.3± 0.75°C (n = 3 replicates).

**Movie S3.**
Phase separation of $d(CAG)_{20}$. A sample of 100 μM $d(CAG)_{20}$ in 25 mM Tris-HCl, (pH 7.5 at 25°C) and 50 mM $MgCl_2$ underwent thermal cycling via temperature-controlled microscopy. $d(CAG)_{20}$ phase separated reversibly with a LCPT of 72.4 ± 1.71°C (n = 3 replicates).

**Movie S4.**
Phase separation of $d(CAG)_{47}$. A sample of 100 μM $d(CAG)_{47}$ in 25 mM Tris-HCl, (pH 7.5 at 25°C) and 50 mM $MgCl_2$ underwent thermal cycling via temperature-controlled microscopy. $d(CAG)_{47}$ phase separated reversibly with a LCPT of 46.3 ± 3.0°C (n = 3 replicates).

**Movie S5.**
Phase separation of $r(CAG)_{31}$. A sample of 100 μM $r(CAG)_{31}$ in 25 mM Tris-HCl, (pH 7.5 at 25°C), 50 mM $MgCl_2$, and 25 mM NaCl underwent thermal cycling via temperature-controlled microscopy. $r(CAG)_{31}$ phase separated irreversibly with a LCPT of 41.3 ± 1.5°C (n = 3 replicates).

**Movie S6.**
Phase separation of $d(CAG)_{31}$. A sample of 100 μM $d(CAG)_{31}$ in 25 mM Tris-HCl, (pH 7.5 at 25°C), 50 mM $MgCl_2$, and 25 mM NaCl underwent thermal cycling via temperature-controlled microscopy. $d(CAG)_{31}$ phase separated reversibly with a LCPT of 50.1± 2.5°C (n = 3 replicates).

**Movie S7.**
Phase separation of $r(CUG)_{31}$. A sample of 100 μM $r(CUG)_{31}$ in 25 mM Tris-HCl, (pH 7.5 at 25°C), 50 mM $MgCl_2$, and 25 mM NaCl underwent thermal cycling via temperature-controlled microscopy. $r(CUG)_{31}$ phase separated reversibly with a LCPT of 59.2 ± 2.4°C (n = 3 replicates).

**Movie S8.**
Phase separation of $d(CTG)_{31}$. A sample of 100 μM $d(CTG)_{31}$ in 25 mM Tris-HCl, (pH 7.5 at 25°C), 50 mM $MgCl_2$, and 25 mM NaCl underwent thermal cycling via temperature-controlled microscopy. $d(CTG)_{31}$ phase separated reversibly with a LCPT of 72.9 ± 2.0°C (n = 3 replicates).

**Movie S9.**
Phase separation of $r(CGG)_{31}$. A sample of 100 μM $r(CGG)_{31}$ in 25 mM Tris-HCl, (pH 7.5 at 25°C), 50 mM $MgCl_2$, and 25 mM NaCl underwent thermal cycling via temperature-controlled microscopy. $r(CGG)_{31}$ phase separated irreversibly with a LCPT of 50.5 ± 0.73°C (n = 3 replicates).

**Movie S10.**

Phase separation of d(CGG)$_{31}$. A sample of 100 μM d(CGG)$_{31}$ in 25 mM Tris-HCl, (pH 7.5 at 25°C), 50 mM $MgCl_2$, and 25 mM NaCl underwent thermal cycling via temperature-controlled microscopy. d(CGG)$_{31}$ phase separated irreversibly with a LCPT of 60.1 ± 0.87°C (n = 3 replicates).

**Movie S11.**

Phase separation of r(CCG)$_{31}$. A sample of 100 μM r(CCG)$_{31}$ in 25 mM Tris-HCl, (pH 7.5 at 25°C), 50 mM $MgCl_2$, and 25 mM NaCl underwent thermal cycling via temperature-controlled microscopy. r(CCG)$_{31}$ phase separated reversibly with a LCPT of 51.8 ± 0.74°C (n = 3 replicates).

**Movie S12.**

Phase separation of d(CCG)$_{31}$. A sample of 100 μM d(CCG)$_{31}$ in 25 mM Tris-HCl, (pH 7.5 at 25°C), 50 mM $MgCl_2$, and 25 mM NaCl underwent thermal cycling via temperature-controlled microscopy. d(CCG)$_{31}$ phase separated reversibly with a LCPT of 52.4 ± 0.96°C (n = 3 replicates).

**Movie S13.**

Phase separation of (TERRA)$_{10}$. A sample of 50 μM (TERRA)$_{10}$ in 25 mM Tris-HCl, (pH 7.5 at 25°C) and 6.25 mM $MgCl_2$ underwent thermal cycling via temperature-controlled microscopy. (TERRA)$_{10}$ phase separated irreversibly with a LCPT of 65.0 ± 1.8°C (n = 3 replicates). This video is remade from Mahendran et al., 2025 [32,33].

**Movie S14.**

Reversible phase separation of (htelo)$_{10}$. A sample of 50 μM (htelo)$_{10}$ in 25 mM Tris-HCl, (pH 7.5 at 25°C) and 50 mM $MgCl_2$ underwent thermal cycling via temperature-controlled microscopy. d(telo)$_{10}$ phase separated reversibly with a LCPT of 60.4 ± 0.48°C (n = 3 replicates).

**Movie S15.**

Irreversible phase separation of (htelo)$_{10}$. A sample of 50 μM (htelo)$_{10}$ in 25 mM Tris-HCl, (pH 7.5 at 25°C) and 100 mM $MgCl_2$ underwent thermal cycling via temperature-controlled microscopy. d(telo)$_{10}$ phase separated irreversibly with a LCPT of 53.7 ± 0.78°C (n = 3 replicates).

**Movie S16.**

Percolation of (TERRA)$_{10}$. A sample of 50 μM (TERRA)$_{10}$ in 25 mM Tris-HCl, (pH 7.5 at 25°C) and 50 mM $MgCl_2$ underwent thermal cycling via temperature-controlled microscopy. (TERRA)$_{10}$ clusters underwent shape relaxation upon heating from irregular clusters to spherical condensates (n = 3 replicates).

**Movie S17.**

Phase separation of r(CmAG)$_{20}$. A sample of 10 μM r(CmAG)$_{20}$ in 50 mM HEPES, (pH 7.5 at 25°C) and 50 mM $MgCl_2$ underwent thermal cycling via temperature-controlled microscopy. r(CmAG)$_{20}$ phase separated reversibly with a LCPT of 57.7 ± 0.81°C (n = 3 replicates).

**Movie S18.**

Phase separation of r(CAG)$_{20}$. A sample of 10 μM r(CAG)$_{20}$ in 50 mM HEPES, (pH 7.5 at 25°C) and 50 mM $MgCl_2$ underwent thermal cycling via temperature-controlled microscopy. r(CAG)$_{20}$ phase separated irreversibly with a LCPT of 51.4 ± 2.49°C (n = 3 replicates).

**Movie S19.**

$Ca^{+2}$ ion dependent phase separation of $r(CAG)_{31}$. A sample of 10 μM $r(CAG)_{31}$ in 25 mM HEPES, (pH 7.5 at 25°C) and 10 mM $CaCl_2$ underwent thermal cycling via temperature-controlled microscopy. $r(CAG)_{31}$ phase separated irreversibly with a LCPT of 44.9 ± 1.6°C (n = 3 replicates).

**Movie S20.**

Reversible condensation of poly(rC). A sample of 1.5 mg/mL poly(rC) in 20 mM HEPES-MES-NaAc (pH 5.75 at 25°C) and 250 mM $MgCl_2$ underwent thermal cycling via temperature-controlled microscopy. Poly(rC) phase separated reversibly with a LCPT of 38.5 ± 0.4°C (n = 3 replicates).

**Movie S21.**

Irreversible condensation of poly(rC). A sample of 1.5 mg/mL poly(rC) in 20 mM HEPES-MES-NaAc (pH 5.6 at 25°C) and 250 mM $MgCl_2$ underwent thermal cycling via temperature-controlled microscopy. Poly(rC) phase separated irreversibly with a LCPT of 35.7 ± 3.3°C (n = 3 replicates).

**Movie S22.**

Percolation of poly(rC) condensates. A sample of 1.5 mg/mL poly(rC) in 20 mM HEPES-MES-NaAc, (pH 5.5 at 25°C) and 250 mM $MgCl_2$ underwent thermal cycling via temperature-controlled microscopy. Poly(rC) clusters underwent shape relaxation upon heating (n = 3 replicates).

**Movie S23.**

Phase separation of poly(rU). A sample of 1.5 mg/mL poly(rU) in 25 mM Tris-HCl, (pH 7.5 at 25°C) and 400 mM $MgCl_2$ underwent thermal cycling via temperature-controlled microscopy. Poly(rU) phase separated reversibly with a UCPT of 24.2 ± 0.5°C and a LCPT of 1.1 ± 0.3°C (n = 3 replicates). This video is reproduced from Wadsworth et al., 2023 [17].
